# Drug Contraindications in Comorbid Diseases: a Protein Interactome Perspective

**DOI:** 10.1101/2022.01.11.475465

**Authors:** Kalyani B. Karunakaran, Madhavi K. Ganapathiraju, Sanjeev Jain, Samir K. Brahmachari, N. Balakrishnan

## Abstract

Adverse drug reactions (ADRs) are leading causes of death and drug withdrawals and frequently cooccur with comorbidities. However, systematic studies on the effects of drugs in comorbidities are lacking. Drug interactions with the cellular protein-protein interaction (PPI) network give rise to ADRs. We selected 6 comorbid disease pairs, identified the drugs used in the treatment of the individual diseases ‘A’ and ‘B’– 44 drugs in anxiety and depression, 128 in asthma and hypertension, 48 in chronic obstructive pulmonary disease and heart failure, 58 in type 2 diabetes and obesity, 58 in Parkinson’s disease and schizophrenia, and 84 in rheumatoid arthritis and osteoporosis – and categorized them based on whether they aggravate the comorbid condition. We constructed drug target networks (DTNs) and examined their enrichment among genes in disease A/B PPI networks, expressed across 53 tissues and involved in ~1000 pathways. To pinpoint the biological features characterizing the DTNs, we performed principal component analysis and computed the Euclidean distance between DTN component scores and feature loading values. DTNs of disease A drugs not contraindicated in B were affiliated with proteins common to A/B networks or uniquely found in the B network, similarly regulated common pathways, and disease-B specific pathways and tissues. DTNs of disease A drugs contraindicated in B were affiliated with common proteins or those uniquely found in the A network, differentially regulated common pathways, and disease A-specific pathways and tissues. Hence, DTN enrichment in pathways, tissues, and PPI networks of comorbid diseases will help identify drugs contraindications in comorbidities.

## 1. Introduction

Comorbidity is the phenomenon in which one or more diseases co-exist with a primary disease in patients. Comorbidities are the norm rather than exceptions among chronic conditions and pose a significant threat to the physical and psychosocial wellbeing of patients [1]. Comorbidities increase with age, and the risk of mortality increases with the number of comorbidities. A longitudinal study (1992-2006) has shown that the mortality risk increased by 25% in patients with 3-4 chronic comorbidities and by 80% in those with 5 or more comorbidities, both in comparison with individuals having no chronic conditions [2]. The prevalence of comorbidities increases from 10% in 0-19 year-olds to 78% in individuals aged 80 or more [3]. The prevalence of comorbidity in women of age groups of 18-44 years, 45-64 years, and ≥65 years was 68%, 95%, and 99% and in men, it was 72%, 89%, and 97% [4]. As per the US National Comorbidity Survey Replication (NCS-R) survey, 73.8-98.2% of the respondents reported having at least one comorbid condition along with a primary condition [1]. The most striking finding from this report was that the estimates of individual disease burden based on the respondents’ perception of their health condition decreased substantially when adjusted for comorbidity [1]. This effect was particularly magnified for neurological disorders, chronic pain, anxiety disorders, major depressive disorder, and diabetes, all of which contribute immensely to the global disease burden [1]. For example, anxiety disorders collectively affect 284 million people (63% females, 2.5-7% variation by country) around the world, and are among the most prevalent mental health and neurodevelopmental disorders (WHO and IHME, 2017) [5].

Disease comorbidity may increase the likelihood of experiencing adverse drug reactions [6–8]. Drugs that are beneficial in the treatment of one disease may aggravate or even cause comorbid conditions, giving rise to adverse drug reactions, e.g. beta-blockers that treat hypertension and heart disease may aggravate asthma [6], trimethoprim and sulfamethoxazole to treat AIDS may increase the patient’s susceptibility to Stevens-Johnson syndrome and toxic epidermal necrolysis [7]; malaria patients with AIDS and osteoarthritis treated with artemisinin based combination antimalarial therapy were 3 times more likely to experience adverse side effects [8]. Serious adverse drug reactions constitute the fourth leading cause of death in the U.S. with 100,000 deaths per year, and about 2 million patients in the U.S. experiencing adverse drug reactions per year [9]. Patient fatalities have led to the withdrawal of 19 drugs from the U.S. market during 1998-2007 [9]. These aspects highlight the importance of reexamining drug design, and the need to develop drugs in light of disease mechanisms governing comorbidities.

Network medicine is an integrative framework for examining the mechanistic effects of disease-associated genes within the context of the human protein-protein interaction (PPI) network (or the ‘interactome’) [10]. The emerging network medicine paradigm in systems biology has prompted systematic data-driven investigations of the effects of drugs on diseases. It captures the essence of the Fourth Paradigm, i.e. Data-Intensive Scientific Discovery [11, 12]. This framework allows data capture and combines theory and computation to facilitate the translation of biological data into biologically insightful and clinically actionable results. The primary applications of this framework are uncovering disease-associated genes, identifying biomarkers that will improve disease screening, clinical diagnosis, and patient stratification, and prioritizing drug targets and pathways for therapeutic intervention [12].

Drugs that target proteins may perturb the PPI network to elicit the intended therapeutic response or an unintended adverse event or side effect [13]. The extensive interconnectivity of the network components suggests that perturbations at the genomic or proteomic level that affect PPIs may disrupt cellular functions and affect other proteins in the neighborhood network, posing deeper implications for several aspects of the disease such as comorbidity and phenotypic responses to drugs [10].

Although the side effects or adverse events precipitated by drugs in specific diseases have been investigated within the framework of the PPI network [14–19], the effects of multiple drugs and their contraindications on comorbid conditions remain largely unexplored. Some studies have provided key insights on the influence of disease-associated PPI networks, biological pathways, and tissues on drug action. Pairs of drugs used for the same disease have shown significant adverse events when the network modules of their protein targets overlap with each other or with a network of disease-associated genes (‘overlapping exposure’, statistical significance p-value ≤ 0.007), e.g. the anti-hypertensive drug nadroparin increased hyperkalemia, an adverse effect of spironolactone, another anti-hypertensive drug [20]. The targets of both cancer and non-cancer drugs were enriched by 1.8 folds among tissue-specific proteins (p-value = 2E-06), and this enrichment became magnified to 2.3 folds when the targets of non-cancer drugs were considered alone [21]. Drugs that are currently in the market are twice as likely to act on tissue-specific proteins than on housekeeping proteins [22].

In this study, we attempt to elucidate the mechanisms underlying drug contraindications in pairs of comorbid diseases. Our findings suggest the relationship between the PPI networks of disease-associated proteins and drug targets, and the pathway membership and tissue specificity of the drug target networks as critical biological factors influencing adverse drug reactions in comorbidities.

## 2. Methods

### 2.1 Compilation of drugs indicated for specific diseases

The Drug Bank database [23] (version 5.1.8) was used to compile the lists of drugs indicated for each of the 14 diseases. After compiling these lists, we used the TWOSIDES database [24] (version 0.1) – a publicly available database of drugs and associated adverse events – to categorize these drugs with respect to their effects on the disease pairs, specifically, (a) drugs effective in disease A and not contraindicated in disease B, (b) drugs effective in disease B and not contraindicated in disease A, (c) drugs effective in disease A and contraindicated in disease B, and (d) drugs effective in disease B and contraindicated in disease A. Drugs associated with specific adverse effects (belonging to (c) and (d)) were identified using their ‘condition concept names’ (descriptions of adverse events). The lists of the condition concept names used for identifying the drugs belonging to the 4 groups for each of the disease pairs can be found in **Additional File 1: Table S1**, and the drug lists can be found in **Additional File 2: Table S2**.

### 2.2 Construction of drug target protein-protein interaction (PPI) networks

The proteins targeted by the drugs (**Additional File 3: Table S3**) belonging to the 4 categories were retrieved from the Drug Bank database [23] using the DGIdb (drug gene interaction database) web portal [25]. The PPIs of these drug targets in the human interactome were compiled from Human Protein Reference Database (HPRD; version 9) [26] and the Biological General Repository for Interaction Datasets (BioGRID; version 4.3.194) [27] using the Cytoscape plugin, Bisogenet [28]. The network building options were: organism - *Homo sapiens*, biorelation type - *protein-protein interaction*, data sources - *BioGRID and HPRD*, method - *input nodes and its neighbors upto a distance of 1*.

### 2.3 Compilation of disease-associated genes

The genes associated with each of the 14 diseases in the 3 non-comorbid pairs and 6 comorbid pairs were compiled from the DisGeNET database [29] (version 7). The non-comorbid pairs were (I) Multiple sclerosis (DisGeNET ID: C0026769) – Peroxisomal disorders (C0282528), (II) Schizophrenia (C0036341) – Rheumatoid arthritis (C0003873), (III) Asthma (C0004096) – Schizophrenia (C0036341). The comorbid pairs were (IV) Anxiety (C0003467) – Depression (C0011570), (V) Asthma (C0004096) – Hypertension (C0085580), (VI) Chronic obstructive pulmonary disorder (COPD) (C0024117) – Heart failure (C0018801), (VII) Type 2 diabetes (C0011860) – Obesity (C0028754), (VIII) Rheumatoid arthritis (C0003873) – Osteoporosis (C0029456) and (IX) Parkinson’s disease (C0030567) – Schizophrenia (C0036341) (**Additional File 4: Table S4**). 100 top-ranking genes associated with each of the diseases were curated based on their gene-disease association scores (GDA). Although the range of the GDA scores among the 100 top-ranking genes varied across our selected diseases (multiple sclerosis (0.11-0.5), peroxisomal disorders (0.01-0.32), schizophrenia (0.43-0.9), rheumatoid arthritis (0.33-0.7), asthma (0.29-0.7), anxiety (0.1-0.5), mental depression (0.34-0.6), essential hypertension (0.03-0.063), chronic obstructive airway disease (0.11-0.9), heart failure (0.3-0.6), non-insulin-dependent diabetes mellitus (0.4-1), obesity (0.4-1), osteoporosis (0.13-0.9) and Parkinson’s disease (0.23-0.7)), a minimum GDA of ≥ 0.01 was chosen to ensure that at least one publication has linked the gene in question with the disease. Note that ‘association’ of a gene with a disease here does not imply causality in most cases and may only indicate an association with disease susceptibility or an endophenotype.

### 1.4 Construction of disease protein-protein interaction (PPI) networks

The PPI networks of the proteins encoded by the disease-associated genes were assembled by extracting their protein interactors from the PPI repositories BioGRID [27] and HPRD [26] using BisoGenet [28] and the network building options specified before. The input nodes for the construction of each of the disease networks were the 100 top-ranking genes compiled from the DisGeNET database.

### 2.5 Calculation of network similarity measures

Matching node ratio (N_M_) was measured as the ratio of the total number of common nodes shared between the two PPI networks of a comorbid pair and the total number of unique nodes in the two disease networks [30].

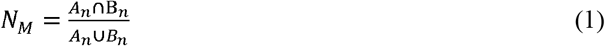

*A_n_* = Number of nodes in the PPI network of disease A
*B_n_* = Number of nodes in the PPI network of disease B

Matching link ratio (L_M_) was measured as the ratio of the total number of common links (i.e. edges) shared between the two PPI networks of a comorbid pair and the total number of unique links in the two disease networks [30].

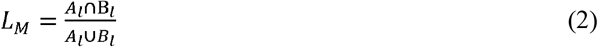

*A_1_* = Number of links in the PPI network of disease A
*B_1_* = Number of links in the PPI network of disease B

The same formula shown above was also used to calculate the matching link ratio for common links of path length 2 and path length 3. Links of specific path lengths were retrieved using the Cytoscape application called NetworkAnalyzer [31, 32].

### 2.6 Calculation of comorbid associations

Relative risk (RR_AB_) measures comorbidity by comparing the observed prevalence of a pair of comorbid diseases (A and B) in the population with the expected number, which is calculated based on the prevalence of the individual diseases A and B in the population.

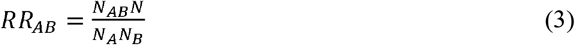

*N_A_* = Total number of patients diagnosed with disease A
*N_B_* = Total number of patients diagnosed with disease B
*N_AB_* = Total number of patients diagnosed with both disease A and disease B
*N* = Total number of patients in the population

For the calculation of relative risks of disease pairs, we downloaded the HuDiNe dataset (http://sbi.upf.edu/data/hudine/) containing processed hospital claims data of 13,039,018 U.S. individuals who had applied for support from the U.S. Medicare program during 1990-1993 [33]. Comorbidity data was available for five out of our six comorbid disease pairs and two out of the three non-comorbid pairs in HuDiNe. Specifically, data was not available for Anxiety – Depression and Multiple sclerosis – Peroxisomal disorders. Hence, N_A_, N_B_ and N_AB_ were extracted for seven out of the nine disease pairs. The diseases were specified in the form of their ICD-9 codes (at three digits level): asthma (ICD-9: 493), hypertension (ICD-9: 401), type 2 diabetes (ICD-9: 250), obesity (ICD-9: 278), chronic obstructive pulmonary disease (ICD-9: 496), heart failure (ICD-9: 428), Parkinson’s disease (ICD-9: 332), schizophrenia (ICD-9: 295), rheumatoid arthritis (ICD-9: 714) and osteoporosis (ICD-9: 733). The population size N was considered to be 13,039,018, i.e. the total number of individuals represented in the HuDiNe dataset.

### 2.7 Pathway enrichment analysis

WebGestalt [34] was used to compute the distribution of genes involved in specific signalling pathways in the drug target networks, and compare it with the background distribution of genes belonging to this pathway among all the genes associated with any pathway in the selected database (Reactome) [35]. Statistical significance of the enrichment was computed using Fisher’s exact test and corrected using the Benjamini-Hochberg method for multiple test adjustment.

### 2.8 Gene expression enrichment analysis

The enrichment of the drug target networks in genes expressed in specific tissues was computed using RNA-sequencing data from 53 postnatal human tissues extracted from GTEx [36] (version 8). Genes with high or medium expression (transcripts per million (TPM) ≥ 9) in 53 tissues were included, provided that they were not housekeeping genes, i.e. genes detected in all the tissues with transcripts per million ≥ 1, as identified in the Human Protein Atlas [37]. TPM is a metric for quantifying gene expression; it directly measures the relative abundance of transcripts. The GMT files served as inputs for a gene over-representation analysis (GSEA) based on hypergeometric distribution. The following GWAS datasets were selected in TSEA-DB [38] for identification of disease-specific tissues (trait IDs are given in parentheses): anxiety (4679), depression (5315), chronic obstructive pulmonary disease (571), heart failure (5333), asthma (5259), hypertension (169), type 2 diabetes (4628), obesity (1031), Parkinson’s disease (4607), schizophrenia (5215), rheumatoid arthritis (4614) and osteoporosis (746).

BaseSpace Correlation Engine (https://covid-19.ce.basespace.illumina.com/c/nextbio.nb) was used to identify the correlations between the gene expression profile induced by maprotiline in PC3 cells (Broad Connectivity Map (CMAP 2.0) [39]), the expression profile associated with major depressive disorder and generalized anxiety disorder (GSE98793 [40]) and the expression profile of adrenal cortex. The software uses a non-parametric rank-based approach to compute the extent of enrichment of a particular set of genes (or ‘bioset’) in another set of genes [41].

### 2.9 Principal component analysis

Principal component analysis (PCA) was used to capture relationships between the drug target networks and the disease networks/biological pathways/tissues. For each disease pair, negative log-transformed p-values indicating the statistical enrichment of the disease networks/biological pathways/tissues in the 4 drug target networks were assembled into a data matrix containing disease networks/biological pathways/tissues as rows and drug target networks as columns; each cell in the matrix contained a −log_10_P value. Following the established approach [42], log transformation was performed to reduce the influence of extreme values on the obtained PCs. PCA was performed with a web-based tool called ClustVis (https://biit.cs.ut.ee/clustvis/) [43]. The data matrix was pre-processed such that 70% missing values were allowed across the rows and columns. The −log_10_P values in the matrix were further centred using the unit variance scaling method, in which the values are divided by standard deviation so that each row or column has a variance of one; this ensures that they assume equal importance while finding the components. The method called singular value decomposition (SVD) with imputation was used to extract principal components. In this method, missing values are predicted and iteratively filled using neighbouring values during SVD computation, until the estimates of missing values converge. The factor/component loadings corresponding to the disease networks/pathways/tissues that contributed to the selected principal components were also extracted. Component loadings are correlation coefficients between the variables in rows and the factors (i.e. PC1, PC2 etc.). The squared value of a component loading gives the percentage of the variance explained by a particular original variable, and essentially its contribution to the principal components. Finally, for each of the disease pairs, the Euclidean distance between the principal component scores of each of the drug target networks were computed for all the component loading values pertaining to the particular biological modality. This resulted in a list of the specific disease protein sets/pathways/tissues that may be closely related to each of the different drug target networks.

## 2. Results

To identify potential mechanisms of adverse drug interactions within comorbid diseases, we systematically studied pairs of comorbid diseases (‘disease A’ and ‘disease B’) and their FDA-approved drugs. We separated the drugs into two groups, namely, disease A drugs that are (a) contraindicated and (b) not contraindicated in disease B, and disease B drugs that are (c) contraindicated and (d) not contraindicated in disease A We then constructed the interactomes of the proteins targeted by these drugs and examined these drug target interactomes in the context of three biological factors, namely, (i) proteins exclusive to interactomes of diseases A and B and those that are in their intersection, and (ii) biological pathways and (iii) tissues associated with these drug target interactomes.

Specifically, we selected three pairs of non-comorbid diseases as negative controls and six pairs of comorbid diseases for our analysis. The non-comorbid pairs were: (I) Multiple sclerosis – Peroxisomal disorders [44], (II) Schizophrenia – Rheumatoid arthritis [45–47], (III) Asthma – Schizophrenia [48]. The comorbid pairs were (IV) Anxiety – Depression [49], (V) Asthma – Hypertension [50, 51], (VI) Chronic obstructive pulmonary disorder (COPD) – Heart failure [52, 53], (VII) Type 2 diabetes – Obesity [54, 55], (VIII) Rheumatoid arthritis – Osteoporosis [56] and (IX) Parkinson’s disease – Schizophrenia [57].

The drugs indicated for use in each of the diseases were retrieved from Drug Bank (version 5.1.8) [23]. For each pair, we categorized the drugs into the four groups (a-d) mentioned earlier, based on their clinical activity in the diseases, collected from the TWOSIDES database (version 0.1) [24], a compendium of drugs and their contraindications (see **Additional File 2: Table S2**). Drugs contributing to specific adverse effects were collected by manually selecting relevant ‘condition concept names’ (**Additional File 1: Table S1**). For example, to identify the anxiolytic drugs that may cause depression, the condition concept names, *depression*, *major depression*, *depressive symptom*, *depression suicidal*, *depression postoperative*, *postpartum depression*, *depressive delusion*, and *agitated depression*, were selected. The list of anxiolytic drugs was then compared with the list of drugs associated with these condition concept names. The matching drugs were compiled into groups ‘a’ and ‘c’, for example, “drugs effective in anxiety and contraindicated in depression”. Similarly, groups ‘b’ and ‘d’ drugs were compiled. The proteins targeted by the drugs belonging to groups a and b were retrieved by querying the Drug Bank database through the DGIdb drug-genee interaction database) web portal [25] (see **Additional File 3: Table S3**). Finally, the protein-protein interaction (PPI) networks of the drug targets were assembled by extracting their protein interactors from the PPI repositories BioGRID [27] (version 4.3.194) and HPRD [26] (version 9) using a Cytoscape plugin called BisoGenet [28].

The methodology of our study is illustrated in **Fig. 1**. To characterize the 4 classes of drug target networks (DTNs), we examined 3 types of data that may reflect their biological profiles, namely (i) disease PPI networks, (ii) biological pathways and (iii,) tissue gene expression. Specifically, we conducted gene overrepresentation analyses based on hypergeometric distribution to check the enrichment of the DTNs among proteins that are unique to/shared between networks of disease A and disease B, genes showing high/moderate expression in 53 tissues across the human body, and proteins involved in ~1000 biological pathways. Overlaps computed in this manner with each of the 3 types of biological data were considered to be statistically significant at p-value < 0.05 after multiple test adjustments with the Benjamini-Hochberg method.

**Figure 1:**
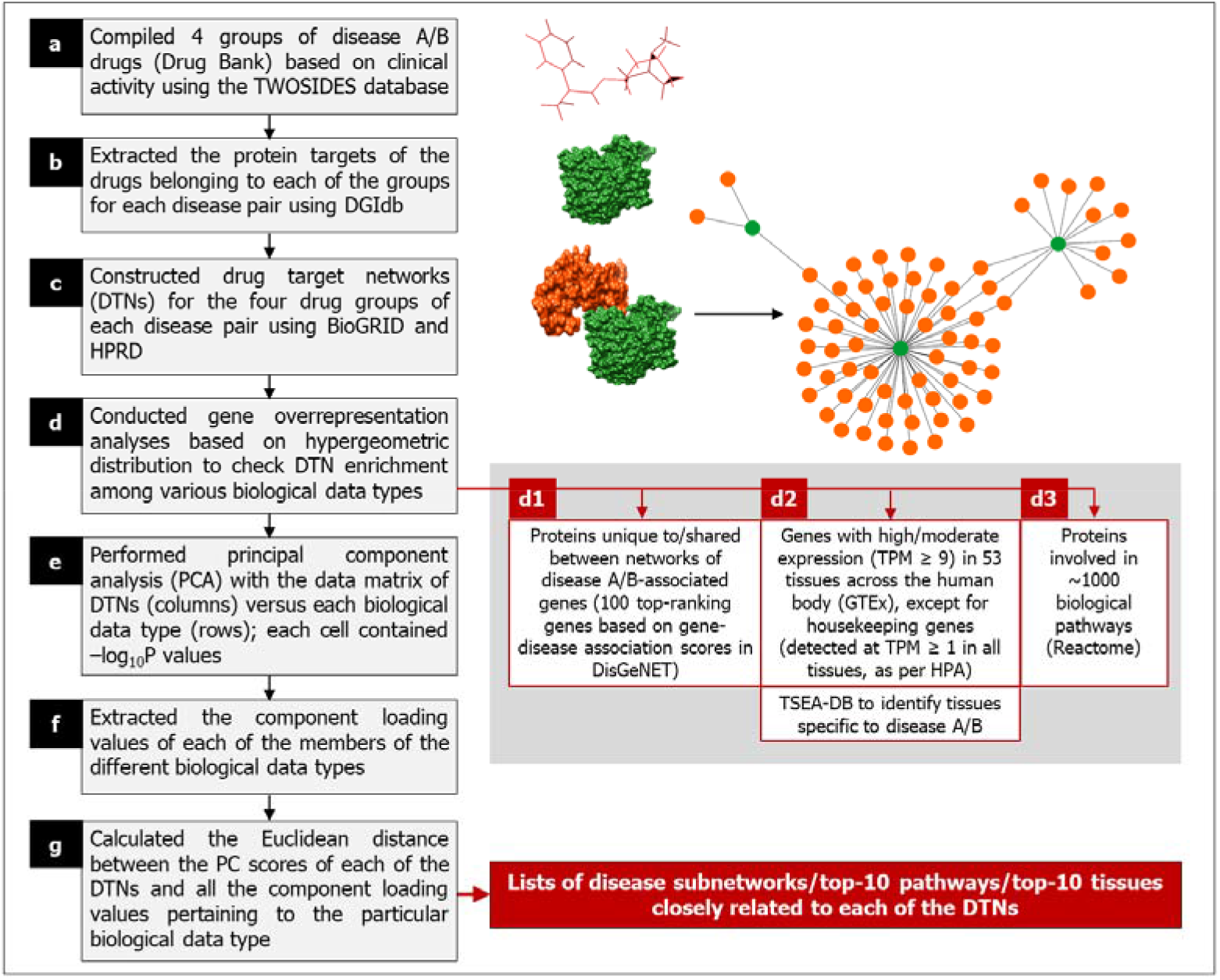
Framework for characterizing the drugs that target comorbid disease pairs. Our methodology to characterize drug target networks (DTNs) contained seven steps: (a) Retrieval of the drugs indicated for use against each of the diseases using Drug Bank and their categorization into four groups based on their clinical activity in the comorbid diseases, namely, disease A drugs not contraindicated in disease B, disease B drugs not contraindicated in disease A, disease A drugs contraindicated in disease B and disease B drugs contraindicated in disease A. (b) Identification of the proteins collectively targeted by the drugs in each of the groups by querying Drug Bank through DGIdb. (c) Construction of DTNs using the protein targets as input nodes and assembling their immediate neighbors in the human protein-protein interaction network up to a distance of 1, based on data from the PPI repositories BioGRID and HPRD. (d) Performing gene enrichment analysis with the four DTNs (corresponding to each of the disease pairs) in 3 biological data types: (d1) disease protein-protein interaction networks, (d2) tissue gene expression and (d3) biological pathways. (e) Generation of a data matrix containing the enriched disease protein sets/tissues/pathways as rows, DTNs as columns and log-transformed p-values in each of the cells, and using the matrix as an input for principal component analysis. (f) Extraction of component loading values of each of the enriched disease protein sets/tissues/pathways that correspond to each of the principal components. (g) Calculation of the Euclidean distance between the principal component scores of each of the DTNs and the component loading values of the disease protein sets/tissues/pathways. These steps resulted in the identification of the top disease protein sets, tissues and pathways that were closely associated with each of the DTNs. Databases: BioGRID (Biological General Repository for Interaction Datasets), DGIdb (Drug Gene Interaction database), DisGeNET (Disease Gene association NETwork), Drug Bank, GTEx (Genotype-Tissue Expression), HPRD (Human Protein Reference Database), Reactome, TSEA-DB (Tissue-Specific Enrichment Analysis DataBase) and TWOSIDES. Abbreviations: DTN – Drug Target Network, PCA – Principal Component Analysis and TPM – Transcripts Per Million.

As a first step towards identifying the specific biological data modalities (disease subnetworks/pathways/tissues) that were relatively more ‘closer’ to each of the different types of DTNs in terms of Euclidean distance, we generated a data matrix of the DTNs (columns) versus the various members of the biological data modality (rows) (for example, for the data modality ‘disease subnetwork’, the members would be ‘common to both the networks’, ‘unique to disease A network’ and ‘unique to disease B network’ and for the data modality ‘tissue’, the members would be ‘amygdala’, ‘aorta’, ‘lungs’ etc.). Each cell contained the negative of log-transformed p-values. −log_10_ transformed p-values have been used as inputs for PCA in previous studies [58, 59]. Following the established approach [42], log transformation was performed to reduce the influence of extreme values on the obtained PCs. Single value decomposition (SVD) with imputation and unit variance scaling was applied to this matrix to extract principal components that explained the variance observed with each of the data modalities across the DTNs. Principal component analysis (PCA) has been applied to matrices containing gene-level association scores in several studies [59]. PCA is primarily used to capture systematic variations underlying datasets. All the principal components generated after this analysis were considered for our study, since they may together reveal underlying clustering patterns among the different DTNs. Following this, we extracted the component loading values of each of the members of the different data modalities, which correspond to each of the principal components representing the relationships among the DTNs. Component loadings are values depicting the correlation of the original variables in our data matrix — negative log of p-values of enrichment for specific disease subnetworks/pathways/tissues — with each of the extracted principal components. Lastly, we calculated the Euclidean distance between the principal component scores of each of the DTNs specifically in the context of each data modality and all the corresponding component loading values. This yielded a list of the specific disease subnetworks/pathways/tissues that are presumably closely related to each of the different DTNs.

### 3.1 Disease network similarity and comorbid associations

Relative risk is an experiential measure of comorbidity as it compares the observed prevalence of a pair of comorbid diseases in the population with the expected number, which is calculated based on the prevalence of the individual diseases in the population. We then explored whether this information was embedded in the disease networks, i.e., whether the relative risk of comorbidity of the disease pairs would be reflected in the similarity of the disease networks. For each of the comorbid pairs, we computed four established network similarity measures, namely, matching node ratio (NM) for all the nodes shared between the two disease networks, and the matching link ratio (L_M_) [30] for all the (i) shared links (i.e. edges), (ii) shared links of path length 2 (connecting two nodes via one intermediate node) and (iii) shared links of path length 3 (connecting two nodes via two intermediate nodes) between the two disease networks.

We computed the relative risk for each of the disease pairs observed in hospital claims data of 13,039,018 U.S. individuals who had filed for support from the Medicare program during the period of 1990-1993, made available as the HuDiNe dataset [33]. The ICD-9 codes corresponding to pairs of diseases diagnosed as primary and secondary conditions, along with the number of individuals who were diagnosed with diseases A or B or both (NA, NB and NAB, respectively) were available (see **Methods**). Comorbidity data was available for five out of our six comorbid disease pairs (i.e. except for Anxiety – Depression) and two out of the three non-comorbid pairs (i.e. except for Multiple sclerosis – Peroxisomal disorders in HuDiNe.

For each of the diseases considered, the top 100 genes associated with the disease were curated from the DisGeNET database (version 7) [29] based on their gene-disease association (GDA) scores (see **Additional File 4: Table S4**). The GDA score ranges from 0 to 1 and is computed for a gene based on the number of publications supporting its association with the disease, and the number and types of database sources (levels of curation (expert-curated/computationally-predicted) and the model organisms in which the association was validated). The 100 top-ranking genes collected in this manner were used as starting points for the construction of disease networks. Here also, the network is assembled by extracting PPIs from BioGRID and HPRD using a Cytoscape plugin BisoGenet, similar to assembling of DTNs. Then, we systematically conducted network overlap analyses with each of the 9 disease pairs and identified the proteins (a) shared between the two disease networks, (b) unique to disease A and (c) unique to disease B (**Table 1**).

**Table 1:** Overlap of the disease networks. The table shows the statistics of the overlaps shared between the two diseases in each of the nine disease pairs that were examined in our study.

| **Disease pair** | **# Proteins in disease A network** | **# Proteins in disease B network** | **# Shared proteins** | **p-value of overlap** | **Odds ratio of overlap** | **% Shared proteins in disease A network** | **% Shared proteins in disease B network** |
| --- | --- | --- | --- | --- | --- | --- | --- |
| Multiple sclerosis (A) – Peroxisomal disorders (B) | 2418 | 727 | 284 | 5.97E-70 | 2.9 | 12% | 39% |
| Schizophrenia (A) – rheumatoid arthritis (B) | 2662 | 2424 | 918 | 6.86E-208 | 2.56 | 34.5% | 38% |
| Asthma (A) – Schizophrenia (B) | 3041 | 2662 | 1084 | 1.36E-228 | 2.41 | 36% | 41% |
| Anxiety (A) – Depression (B) | 3342 | 3054 | 1732 | 1.86E-628 | 3.06 | 52% | 57% |
| Asthma (A) – Hypertension (B) | 3041 | 2515 | 1371 | 1.85E-500 | 3.23 | 45% | 54.50% |
| Chronic obstructive pulmonary disease (A) –heart failure (B) | 3736 | 2922 | 1505 | 3.12E-371 | 2.48 | 40% | 51.50% |
| Type 2 diabetes (A) – Obesity (B) | 2471 | 2490 | 1232 | 3.66E-503 | 3.6 | 50% | 49% |
| Rheumatoid arthritis (A) – osteoporosis (B) | 2424 | 3681 | 1206 | 1.30E-270 | 2.43 | 50% | 33% |
| Parkinson’s disease (A) – schizophrenia (B) | 3200 | 2662 | 1232 | 2.88E-310 | 2.6 | 38.50% | 46% |

The relative risk between diseases was proportional to the matching node and link ratios (**Fig. 2**). The control disease pairs showed low relative risks and smaller disease network overlaps, whereas three out of five comorbid disease pairs showed high relative risks and larger network overlaps, namely, Asthma – Hypertension, COPD – Heart failure and Type 2 diabetes – Obesity. However, this trend was not seen in the comorbid pairs, Rheumatoid arthritis – Osteoporosis and Parkinson’s disease – Schizophrenia. Specifically, their higher relative risks (compared with other comorbid pairs), were not accompanied by a corresponding increase in the network overlaps. ~85% of the human interactome awaits experimental discovery [60]. Hence, two factors may have led to the underestimation of the network overlaps. Firstly, the inherent incompleteness of these disease networks [60]. Secondly, the tendency of incomplete networks to exhibit small overlaps [60].

**Figure 2:**
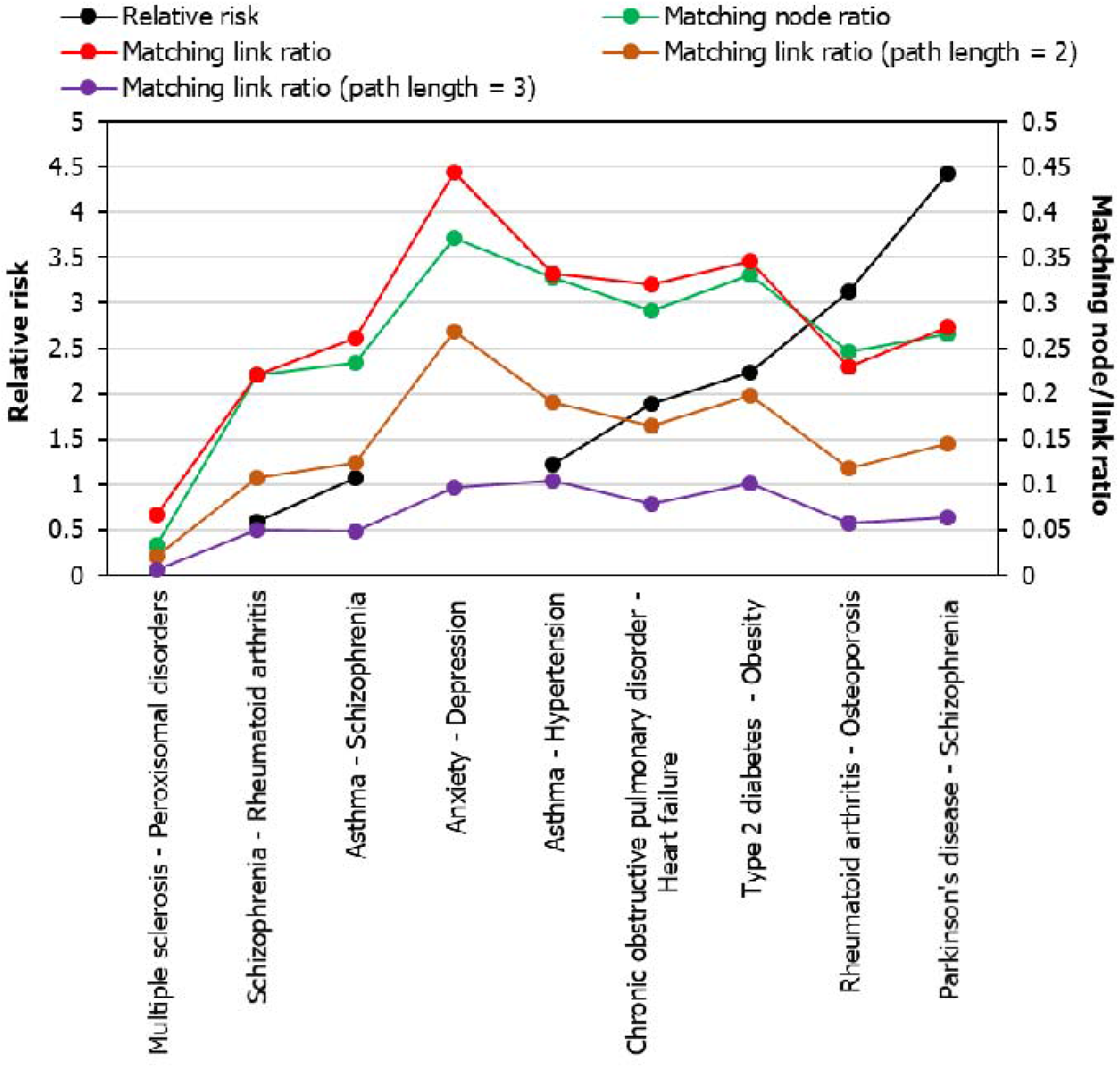
Comparison of disease network similarity measures and comorbid associations. The graph shows the relationship between relative risk (black data points) and four measures of network similarity, namely, matching node ratio (green data points), matching link ratio of all shared edges (red data points), matching link ratio of all shared

### 3.2 Druggability of disease networks

Next, we tested the potential of each of the disease subnetworks to be acted upon by drugs or their susceptibility to pharmacological modulation (druggability), by examining their enrichment among a group of 4,463 proteins deemed to be druggable [61], similar to the approach followed in a previous study [62]. These proteins are bound with high affinity at specific binding sites by drugs that follow the Lipinski’s ‘rule-of-five’, i.e. orally bioavailable drugs with specific molecular characteristics that influence their pharmacokinetic ability to enter systemic circulation and act on their target sites (**Table 2**) [63].

**Table 2:** Overlaps of the disease protein sets with druggable targets. -log_10_P values computed for each of the nine tested disease pairs using a hypergeometric test. The −log_10_P values indicate the statistical significance of the overlaps shared by each of the disease protein sets (top column headings) with a group of 4463 druggable proteins. *, ** and *** indicate low, medium and high levels of statistical significance. ^□^, ^□□^ and ^□□□^ indicate non-significant overrepresentation, non-significant underrepresentation and significant underrepresentation respectively.

| Disease pairs | Common to both the networks | Unique to disease A network | Unique to disease B network |
| --- | --- | --- | --- |
| Multiple sclerosis (A) – peroxisomal disorders (B) | 7.38** | 19.52*** | 2.09* |
| Schizophrenia (A) – Rheumatoid arthritis | 13.26** | 14.36*** | 2.4* |
| Asthma (A) – schizophrenia (B) | 19.18*** | 9.41** | 0.89□ |
| Anxiety (A) – Depression (B) | 5.57** | 0.001$\square\square$ | 12.05*** |
| Asthma (A) – Hypertension (B) | 31.34*** | 3.19* | 9.59** |
| Chronic obstructive pulmonary disease (A) – heart failure (B) | 34.73*** | 1.06$\square\square$ | 9.16** |
| Type 2 diabetes (A) – Obesity (B) | 18.65*** | 1.47* | 7.05** |
| Rheumatoid arthritis (A) – Osteoporosis (B) | 21.96*** | 7.17** | 1.97* |
| Parkinson's disease (A) – Schizophrenia (B) | 19.93*** | 0.27□ | 0.3□ |

We found that the proteins shared between the two diseases were the most significantly enriched for druggable targets in 5 out of the 6 tested comorbid pairs (**Table 2**). In case of the sixth pair, namely anxiety and depression, the proteins that are exclusive to the depression network were found to be more enriched for druggable targets. In 2 out of the 5 disease pairs that shared many common drug targets, the drug target proteins were significantly enriched in G protein-coupled receptor activity (p-value <0.05) (**Table 3**).

**Table 3:** Enrichment of Gene Ontology molecular functions among druggable targets and proteins unique to the depression network. The odds ratio of enrichment of two specific Gene Ontology molecular functions among druggable proteins, proteins unique to the depression network and proteins common to the anxiety and depression networks have been shown. p-values indicating the statistical significance of these enrichments have been shown in parentheses. Note that druggable proteins show higher enrichment for transmitter-gated channel activity compared to G protein-coupled peptide receptor activity, in terms of odds ratio of enrichment. The overrepresentation of a more druggable class (glutamate-gated Ca^2+^ channel activity) among proteins unique to the depression network (and not among the common proteins) would have altered the enrichment pattern for anxiety and depression in comparison with other the disease pairs (as shown in Table 2).

| **Gene Ontology Molecular Function** \ **Protein sets** | 4463 druggable proteins – odds ratio (p-value) | Proteins common to anxiety and depression networks – odds ratio (p-value) | Proteins unique to depression network – odds ratio (p-value) |
| --- | --- | --- | --- |
| Transmitter-gated channel activity | **4.2 (< 1E-15)** | **8.65 (0.037)** | - |
| G protein-coupled peptide receptor activity | 3.8 (< 1E-15) | 3.6 (6.6E-03) | **10.46 (9.3E-07)** |

Based on these observations and the finding in the previous section that relative risk varies in tandem with network similarity measures, we speculated that contraindications in comorbidities may arise from drug action on druggable proteins shared between the networks of comorbid diseases (**Table 2**). This led to two corollaries: (i) the target networks of the group ‘a’ and ‘c’ drugs (effective in disease A and contraindicated in disease B or vice versa) may show the highest enrichment for the proteins/pathways/tissues shared between the two disease networks and (ii) the target networks of the groups ‘b’ and ‘d’ drugs (effective in disease A and *not* contraindicated in disease B or vice versa) may show the highest enrichment for proteins/pathways/tissues unique to disease A (or B respectively).

### 3.3 Disease networks and drug target networks

To test these corollaries, we systematically computed the overlaps between three groups of disease proteins, namely, proteins that are (a) common to disease A and disease B networks, (b) unique to disease A network and (c) unique to disease B network, and four classes of DTNs, namely, target networks of drugs effective in disease A and (a) contraindicated and (b) *not* contraindicated in disease B, and target networks of drugs effective in disease B (c) contraindicated and (d) *not* contraindicated in disease A (**Table 4**); previous studies have examined the overlaps between the PPI networks of drug targets and disease-associated proteins [20, 64]. For each of the six disease pairs, we created a data matrix of DTNs (columns) versus disease subnetworks (rows), which contains −log(p-values) indicating the statistical significance of these enrichments. This data matrix was used as the input for PCA. In order to identify the specific disease subnetworks that were the nearest to each of the DTNs, we calculated the Euclidean distance between the PC scores of each of the DTNs across all the extracted axes and the corresponding component loading values of all the disease subnetworks across these axes (following the methodology depicted in **Fig. 1**). By counting the two disease subnetworks that were the closest to each of the different DTNs, we identified two predominant patterns.

**Table 4:**
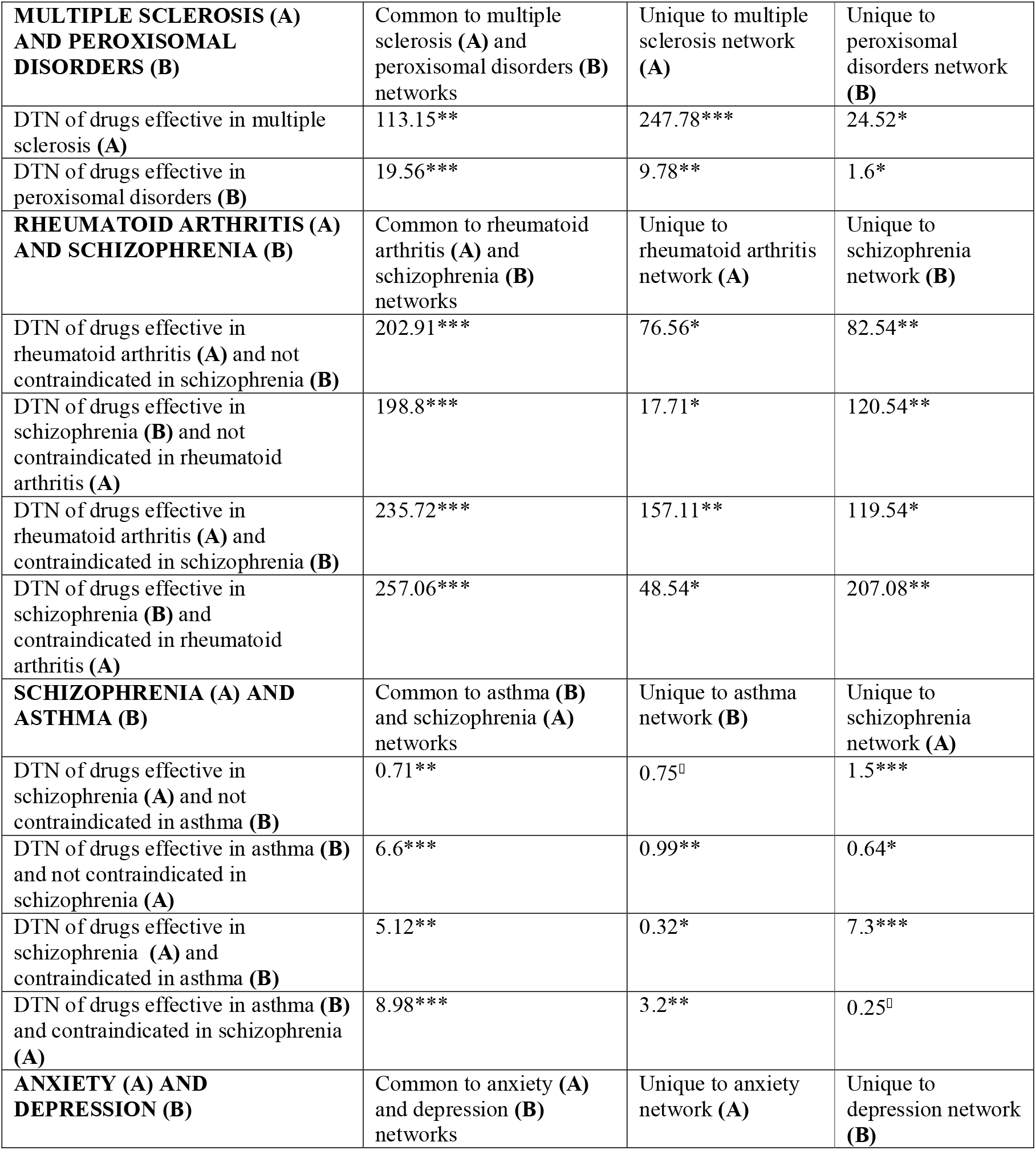
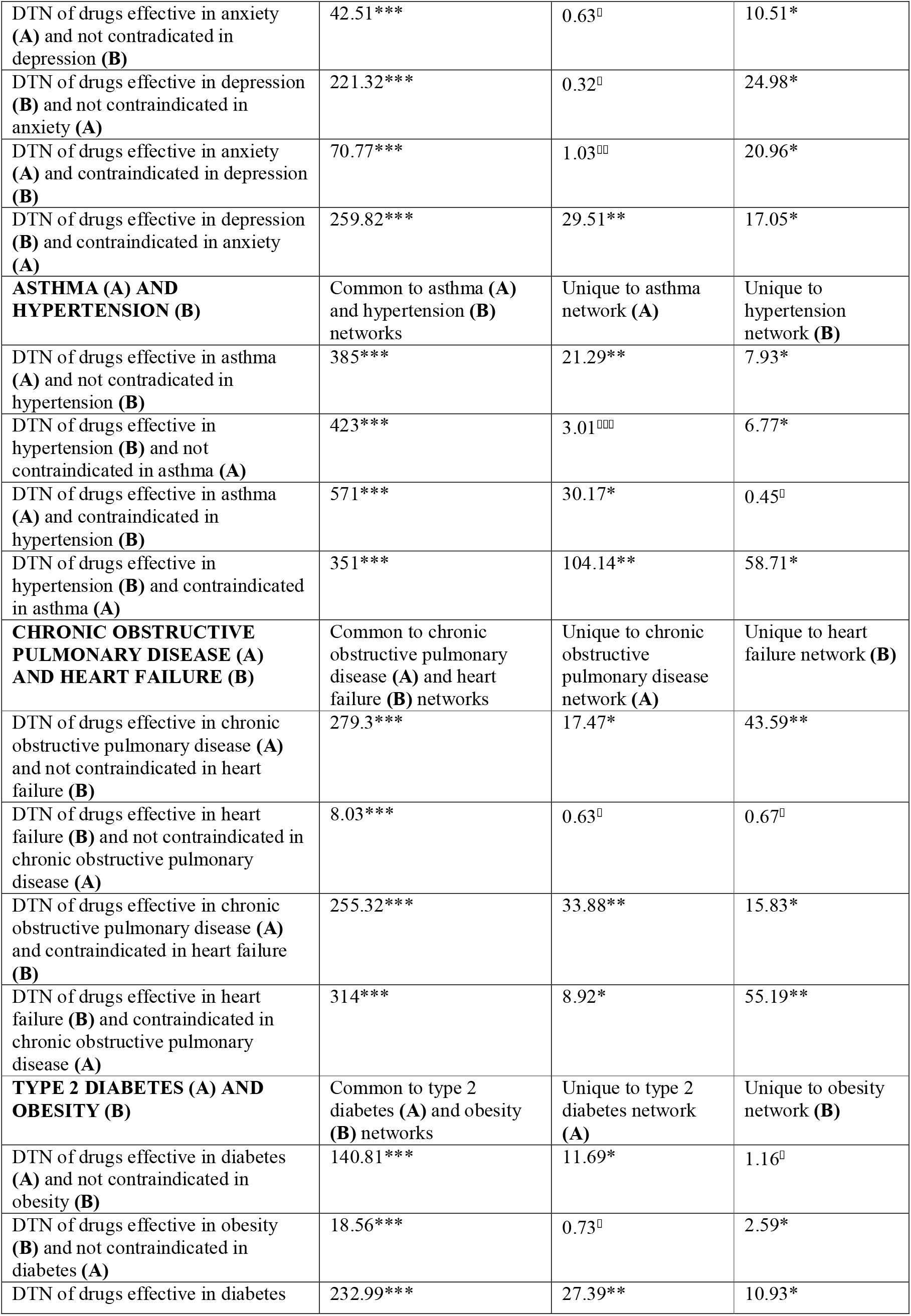
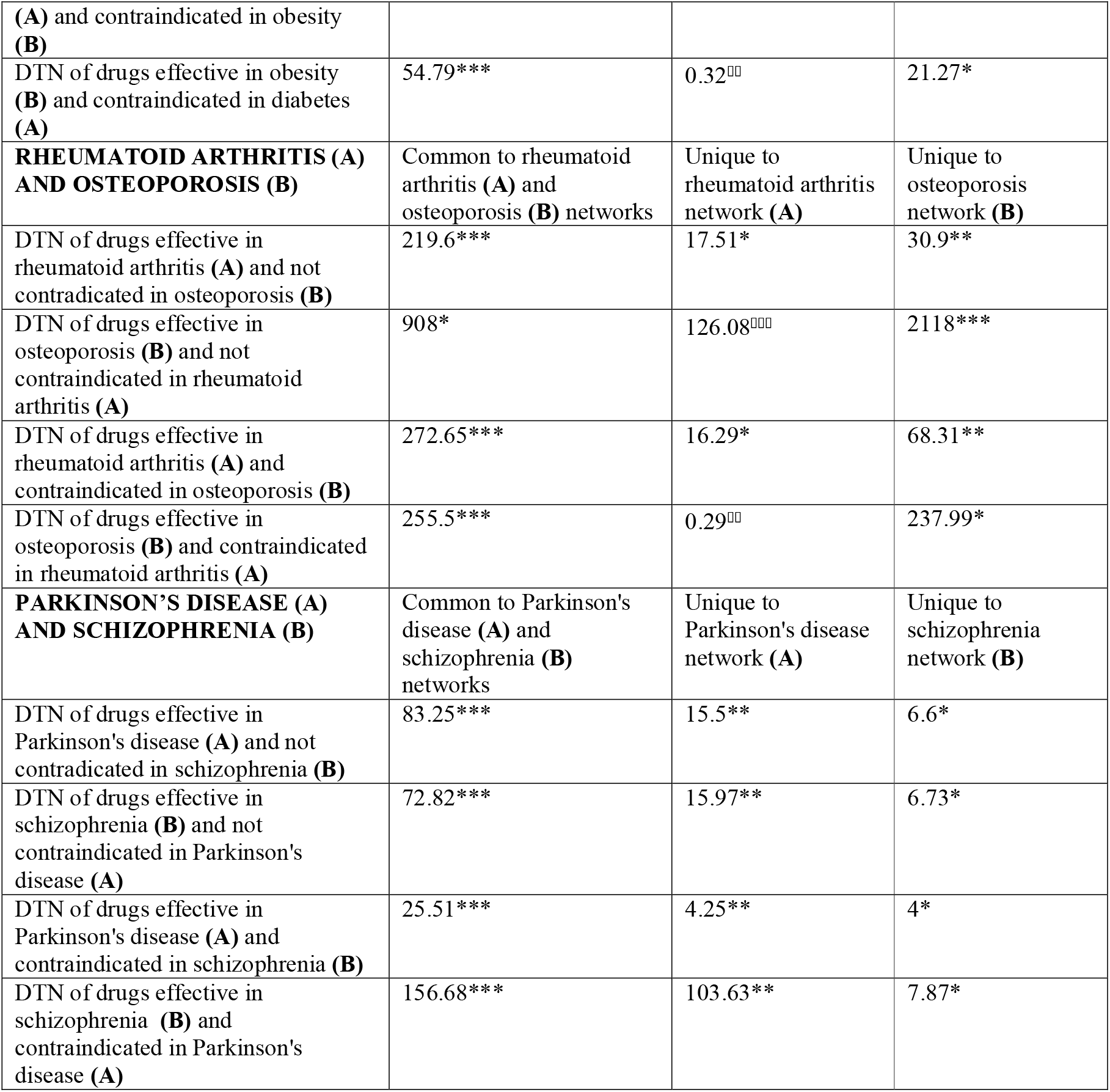
Overlaps of the disease protein sets with the four classes of drug target networks. -log_10_P values computed for each of the nine tested disease pairs using a hypergeometric test. The −log_10_P values indicate the statistical significance of the overlaps shared between each of the disease protein sets (top column headings) and the target networks of the four classes of drugs (row headings). *, ** and *** indicate low, medium and high levels of statistical significance. ^□^, ^□□^ and ^□□□^ indicate non-significant overrepresentation, non-significant underrepresentation and significant underrepresentation respectively.

In 10 out of the 12 cases, the DTNs of drugs used for a specific disease and not contraindicated in a comorbid condition were found to be closest/second closest to the proteins uniquely found in the network of the comorbid condition. Additionally, in 9 out of the 12 cases, they were closest/second closest to the proteins shared between the networks of both the diseases. In contrast, the DTNs of drugs used for a specific disease and contraindicated in a comorbid condition were found to be closest/second closest to the proteins uniquely found in the network of the disease for which these drugs were primarily used in 8 out of the 12 cases. Additionally, in 9 out of the 12 cases, they were closest/second closest to the proteins shared between the networks of both the diseases.

These observations led us to speculate two scenarios. Firstly, disease A drugs that are not contraindicated in disease B may target proteins unique to the disease B subnetwork involved in mechanisms that are either inconsequential/beneficial for disease B, but whose modulation is certainly beneficial for the treatment of disease A. Alternatively, they may target common mechanisms that are dysregulated in a similar manner in both the diseases and pharmacologically modulate them in a similar direction. Secondly, disease A drugs may become contraindicated in disease B when they target either (a) common mechanisms that are pharmacologically oppositely modulated in a manner that benefits disease A but aggravates disease B or (b) mechanisms unique to disease A that aggravate disease B. Additionally, we hypothesized that biological processes such as signalling pathways that function at a higher level than disease subnetworks could be regulating the action of drugs under comorbid conditions.

### 3.4 Biological pathways and drug target networks

We identified the pathway associations of the DTNs using the gene set analysis toolkit called WebGestalt [34]. WebGestalt computes statistical significance enrichment of a functional group (e.g., a Reactome pathway) in an input gene list using Fisher’s exact test using the Benjamini-Hochberg method for multiple test adjustment. For each of the 6 disease pairs, a data matrix of DTNs (columns) versus Reactome pathways (rows) containing corresponding enrichments was used as inputs for PCA, and the Euclidean distance between the PC scores of each of the DTNs across all the extracted axes and the corresponding component loading values of all the pathways across these axes were computed. For each of the disease pairs, we retrieved the top-10 pathways closest to each of the DTNs out of all the pathways enriched in the DTNs (**Additional Files 5-10: Figures S1-S6)**. Confirming our earlier suspicions, we noted that disease A DTN classes without contraindications in disease B were nearest to pathways possibly underlying both the diseases or uniquely associated with B, which are similarly regulated, i.e. upregulated or downregulated together, in the two comorbid diseases. On the other hand, disease A DTN classes with contraindication in disease B were nearest to pathways underlying both the diseases or unique to disease A that are differentially regulated, i.e. upregulated in one disease and downregulated in the other or vice versa.

G alpha (12/13) signalling events’ and ‘muscarinic acetylcholine receptors’ were identified among the top-10 pathways that were close to anxiety drugs not contraindicated in depression in our study (**Fig. 3**). Adrenergic receptor signalling could be regulated via G α(12/13); Gr12 and Gα 13 have been shown to mediate alpha-1 adrenergic receptor-induced JNK activation in rat cardiomyocytes [65]. The drug maprotiline was among our list of anxiety drugs without contraindications in depression (**Fig. 4**). Corroborating this, clinical data suggested that the drug is effective in alleviating anxiety symptoms co-occurring with depression [66]. Maprotiline acts an inhibitor of SLC6A2 (sodiumdependent noradrenaline transporter) and inhibits noradrenaline reuptake in the brain. It also acts as an antagonist to alpha-1 adrenergic receptors (ADRA1A, ADRA1B and ADRA1D) and alpha-2 adrenergic autoreceptors and heteroreceptors (ADRA2A, ADRA2B and ADRA2C), and enhances central noradrenergic and serotonergic functions, which have been linked to alleviation of anxiety and depression [67]. Maprotiline also acts as a weak antagonist to muscarinic acetylcholine receptors (CHRM1, CHRM2, CHRM3, CHRM4 and CHRM5); enhanced cholinergic signaling has been linked to both anxiety and depression [68]. It is notable that maprotiline targets a higher number of proteins associated uniquely with depression (ADRA2A, HTR2C, SLC6A2 and CHRM2) in the adrenergic, serotonergic and cholinergic systems (**Fig. 4**). It targets only one receptor associated with both anxiety and depression (HTR2A), and no gene uniquely associated with anxiety (**Fig. 4**). These observations are in line with our findings with the overlap of DTNs with disease subnetworks, i.e. DTNs of disease A drugs that are not contraindicated in disease B (e.g. maprotiline) are closely associated with proteins uniquely found in the disease B network (i.e. depression in this specific example). Since maprotiline has been discontinued from usage since 2020 in U.S. [69], note that we are citing this drug only as a demonstrative example. Drugs acting on the serotonergic system is known to be effective in both short-term and long-term treatment of patients with major depressive disorder and anxiety disorders [70]. ‘Serotonin receptors’ was identified among the top-10 pathways that were close to depression drugs not contraindicated in anxiety (**Fig. 3**). This may suggest the broadspectrum efficacy of drugs acting on serotonin receptors in both the conditions. Two such drugs in our study displayed antagonistic activity on serotonin receptors – flupentixol [71] and mirtazapine [72] (both acting on HTR2A and HTR2C) – and have been used to treat depression accompanied by anxiety symptoms (**Fig. 4**).

**Figure 3:**
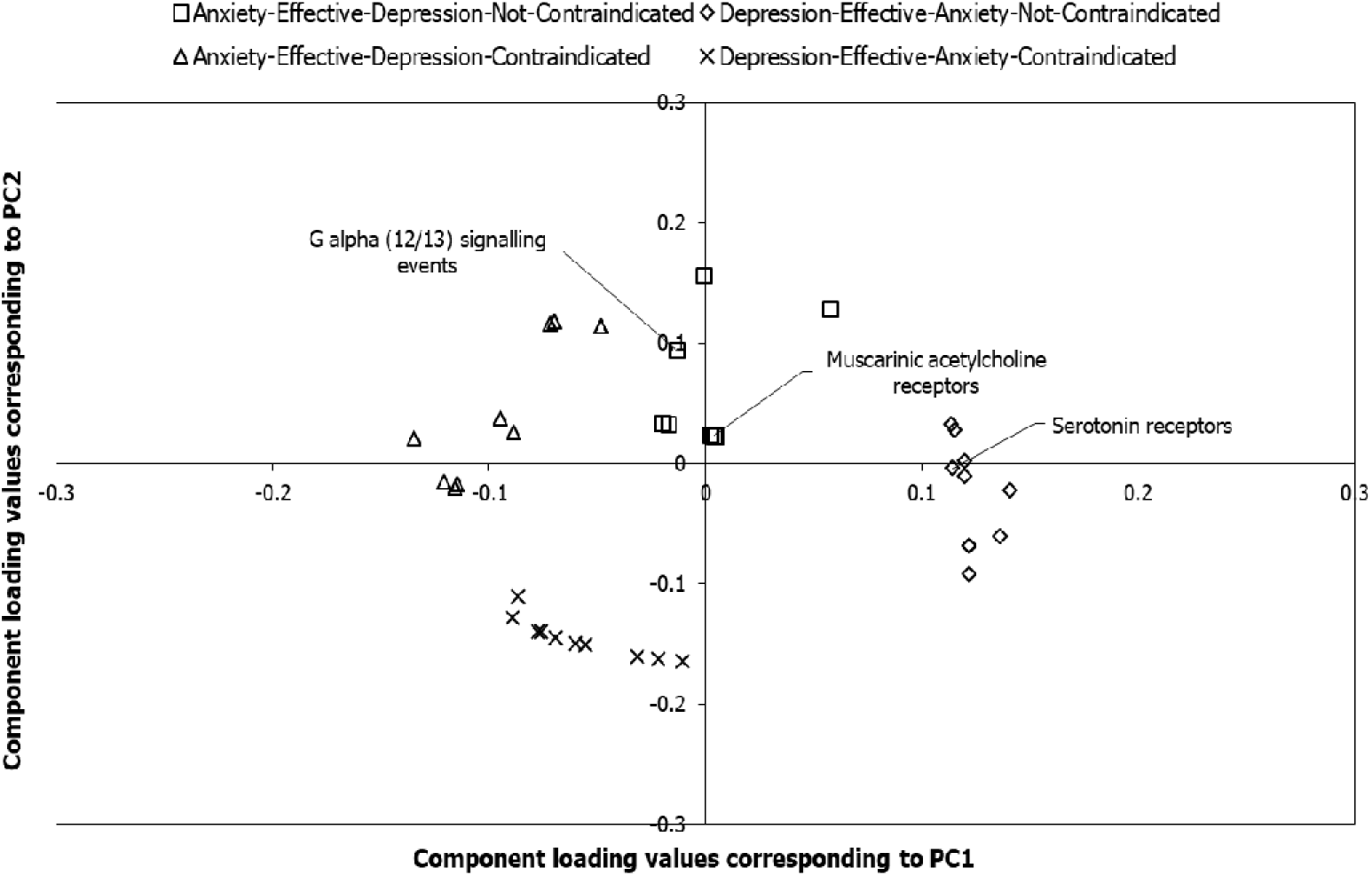
Pathways associated with the target networks of anxiety and depression drugs. The component loading values shown in the figure correspond to component scores of 4 drug target networks (DTNs) of anxiety and depression along PC1 and PC2, which explain 41.6% and 33.1% of the total variance respectively. The top-10 pathways that appeared to be highly related to each of the 4 DTNs, which were obtained after computing the Euclidean distance between the component loading values and the component scores, are shown as square-shaped data points for the DTN of drugs effective in anxiety and not contraindicated in depression, diamond-shaped data points for the DTN of drugs effective in depression and not contraindicated in anxiety, triangle-shaped data points for the DTN of drugs effective in anxiety and contraindicated in depression and cross-mark-shaped data points for the DTN of drugs effective in depression and contraindicated in anxiety. ‘G α(12/13) signaling events’ and ‘muscarinic acetylcholine receptors’ shown here are among the top-10 pathways associated with anti-anxiety drugs that are not contraindicated in depression. The drug maprotiline shown in **Fig. 4** corroborates this by showing antagonistic activity on adrenergic and muscarinic acetylcholine receptors. Similarly, serotonin receptors are associated with anti-depressants that are not contraindicated in anxiety; flupentixol and mirtazapine corroborate this by showing antagnostic activity on serotonin receptors.

**Figure 4:**
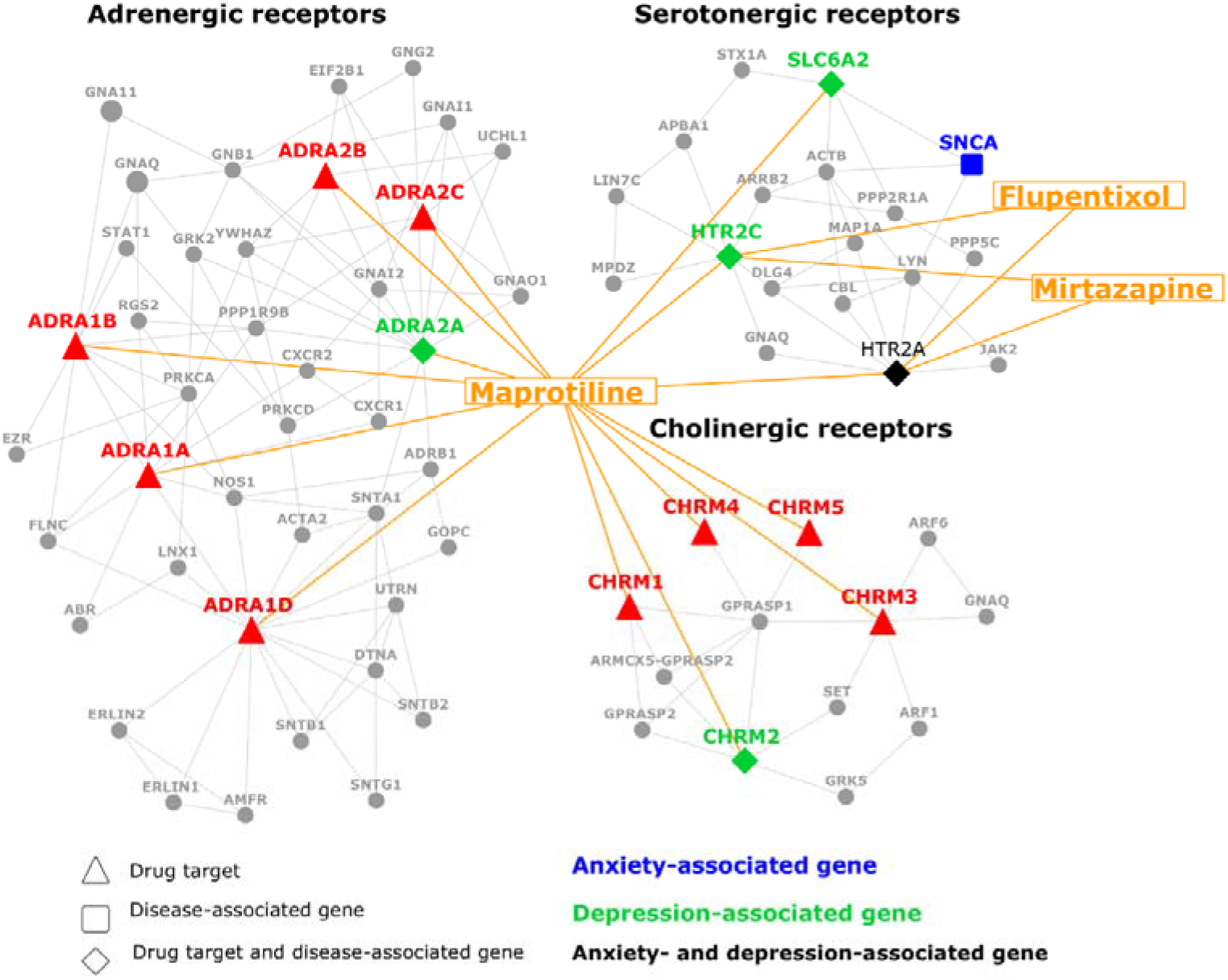
Network diagram showing the relationship between the targets of maprotiline, flupentixol and mirtazapine, and genes associated with anxiety and depression. The different families of receptors and transporter proteins targeted by maprotiline, flupentixol and mirtazapine and their interactions with the proteins encoded by anxiety (disease A) and/or depression (disease B) associated genes have been shown. Note that maprotiline (an anti-anxiety (disease A) drug not contraindicated in depression (disease B)) targets a higher number of proteins associated uniquely with depression in the adrenergic, serotonergic and cholinergic systems, which is in line with our observation that disease A drugs that are not contraindicated in disease B are closely associated with proteins uniquely found in the disease B network (i.e. depression in this specific example). Serotonin receptors were found to be associated in our analysis with depression drugs not contraindicated in anxiety; antagonistic activity on serotonin receptors is shown by two such drugs shown in the diagram (flupentixol and mirtazapine).

The pathway ‘dopamine receptors’ was found to be close to PD drugs contraindicated in SCZ (**Fig. 5**), indicating that the enhancement in dopamine levels brought about by PD drugs may in fact induce SCZ, which has been linked to a hyperdopaminergic state [57]. The dopamine agonists belonging to this group of PD drugs have been shown to induce psychosis, namely, levodopa (acting on DRD1, DRD2, DRD3, DRD4 and DRD5) and ropinirole (DRD2, DRD3 and DRD4) (**Fig. 6**) [73, 74]. It is notable that levodopa and ropinirole target a higher number of dopamine receptors associated uniquely with Parkinson’s disease (DRD1 and DRD2) (**Fig. 6**). It targets only one dopamine receptor (DRD3) uniquely associated with schizophrenia (**Fig. 6**). These observations are in line with our findings with the overlap of DTNs with subnetworks, i.e. DTNs of disease A drugs that are contraindicated in disease B (e.g. levodopa and ropinirole) are closely associated with proteins uniquely found in the disease A network (i.e. Parkinson’s disease in this specific example).

**Figure 5:**
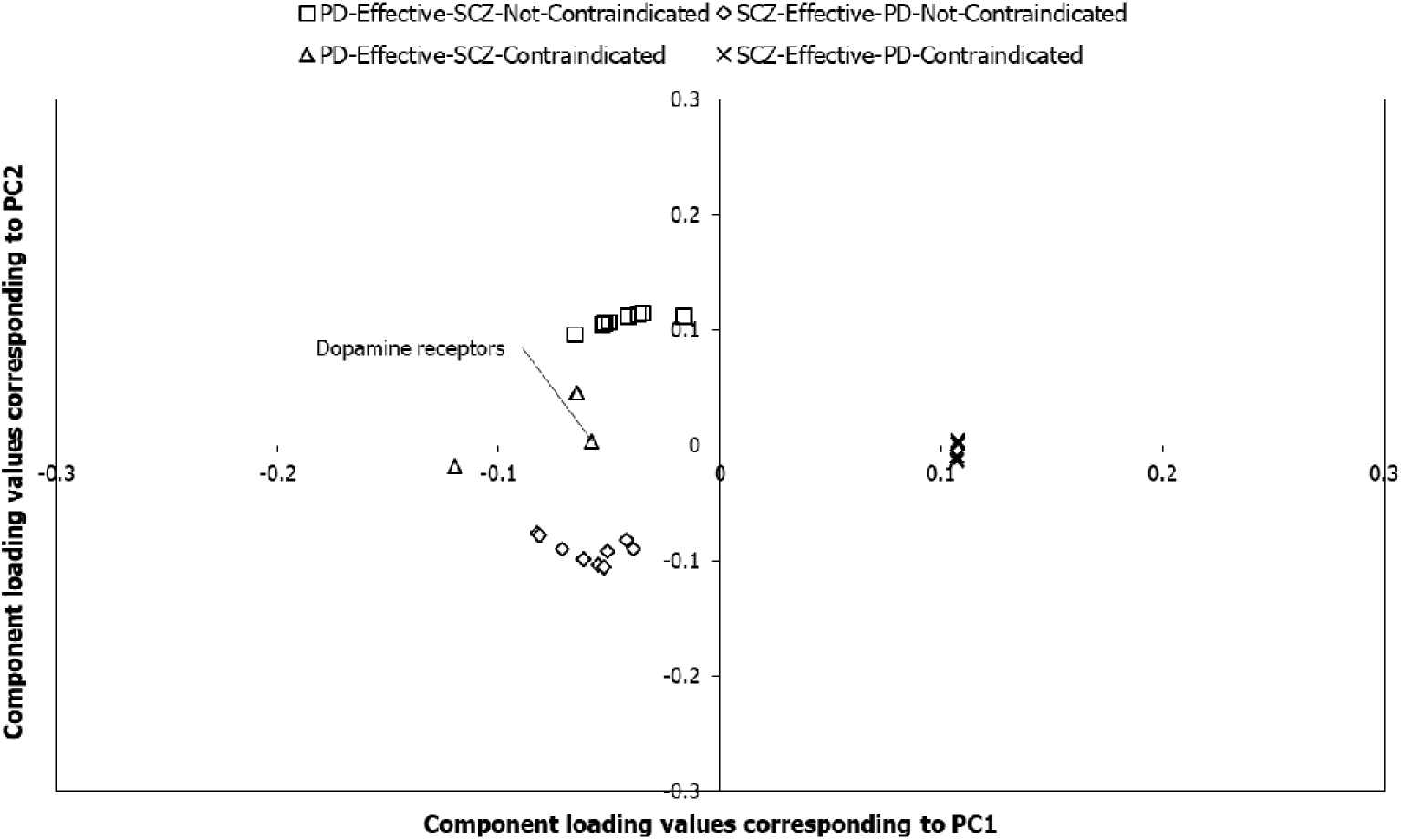
Pathways associated with the target networks of Parkinson’s disease and schizophrenia drugs. The component loading values shown in the figure correspond to component scores of 4 drug target networks (DTNs) of Parkinson’s disease (PD) and schizophrenia (SCZ) along PC1 and PC2, which explain 47.3% and 38.2% of the total variance respectively. The top-10 pathways that appeared to be highly related to each of the 4 DTNs, which were obtained after computing the Euclidean distance between the component loading values and the component scores, are shown as square-shaped data points for the DTN of drugs effective in PD and not contraindicated in SCZ, diamond-shaped data points for the DTN of drugs effective in SCZ and not contraindicated in PD, triangle-shaped data points for the DTN of drugs effective in PD and contraindicated in SCZ and cross-mark-shaped data points for the DTN of drugs effective in SCZ and contraindicated in PD. Dopamine receptors are among the top-10 pathways associated with PD drugs contraindicated in SCZ. Corroborating this, the drugs levodopa and ropinirole shown in **Fig. 6** stimulate dopaminergic receptors to alleviate Parkinsonian symptoms, but at the risk of inducing a hyperdopaminergic state conducive to the SCZ development.

**Figure 6:**
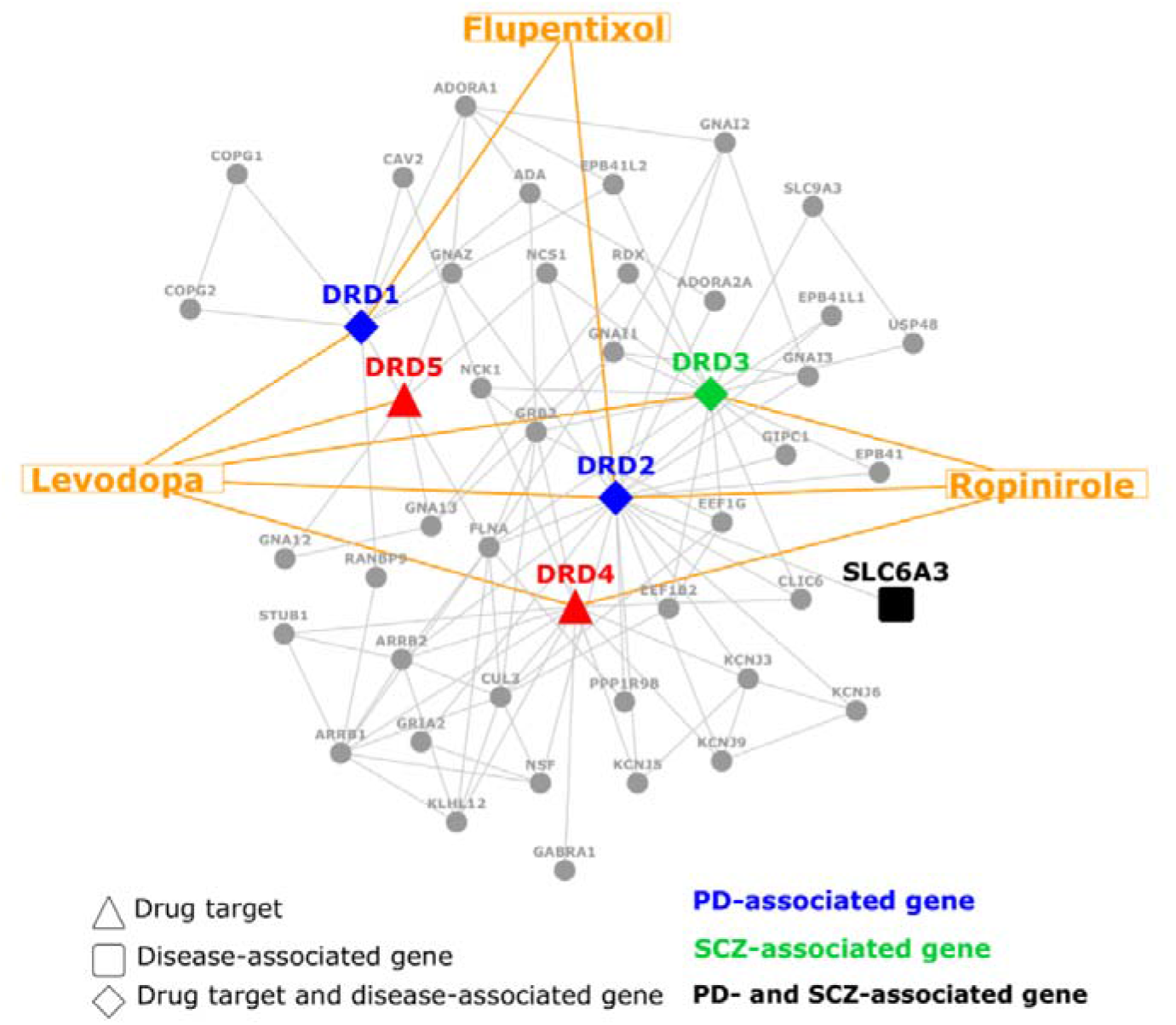
Network diagram showing the relationship between the targets of levodopa and ropinirole and genes associated with Parkinson’s disease and schizophrenia. The specific dopamine receptors targeted by levodopa, ropinirole and flupentixol and their interactions with the proteins encoded by Parkinson’s disease and/or schizophrenia associated genes have been shown. Note that levodopa and ropinirole are used in the treatment of Parkinson’s disease (disease A), but contraindicated in schizophrenia (disease B), and flupentixol is used in the treatment of schizophrenia, but contraindicated in Parkinson’s disease. Note that levodopa and ropinirole target a higher number of dopamine receptors associated uniquely with Parkinson’s disease, which supports our finding that disease A drugs that are contraindicated in disease B are closely associated with proteins uniquely found in the disease A network (i.e. Parkinson’s disease in this specific example).

### 3.5 Tissues and drug target networks

Using RNA-sequencing data of 53 postnatal human tissues obtained from GTEx [36] (version 8), we attempted to identify whether the four DTN classes showed any tissue-specific patterns. Genes with high/medium expression (transcripts per million (TPM) ≥ 9) in these 53 tissues, which were not housekeeping genes (as per the Human Protein Atlas [37]), were considered. For DTNs of each disease pair, we computed the distribution of genes expressed in a specific tissue among the DTN genes and compared it with the background distribution of genes expressed in this tissue among all the genes that were assayed for expression in any of the 53 tissues. We generated a data matrix of DTNs (columns) versus tissues (rows) containing the negative of log-transformed p-values and performed PCA with this matrix as the input. We calculated the Euclidean distance between the PC scores of each of the DTNs and the component loading values of all the tissues. For each of the disease pairs, we retrieved the top-10 tissues that were nearest to the four DTNs (**Additional Files 11-16: Figures S7-S13**). Following this, we employed the tissue-specific enrichment analysis database (TSEA-DB)[38] to retrieve the top-3 tissues that may be preferentially affiliated with the diseases in each of the pairs. TSEA-DB is a reference database for information on disease-associated tissues, specifically, the tissues in GTEx that show significant enrichment of genes harbouring diseases-associated variants compiled from the GWAS catalog [38]. We checked whether the top-3 tissues associated with each of the diseases in a disease pair (according to TSEA-DB) appeared among the list of tissues identified to be nearest to each of the 4 DTNs pertaining to this disease pair in our analysis. Out of the 11 tissues identified to be closer to the target networks of drugs used for a specific disease and not contraindicated in a comorbid condition, 6 were found to be associated with the comorbid condition as per TSEA-DB, whereas 3 were associated with the specific disease for which the drugs were used and 2 were associated with the disease as well as the comorbid condition. Conversely, out of the 9 tissues identified to be closer to the target networks of drugs used for a specific disease and contraindicated in a comorbid condition, 5 were found to be associated with the specific disease, whereas 3 were associated with the comorbid condition in which the drugs were contraindicated and one was associated with the disease as well as the comorbid condition.

These percentages obtained with a low number of tissues suggest cautious interpretation.

Nevertheless, these results seem to corroborate our previous findings with disease subnetworks and biological pathways. Specifically, the networks of disease A drugs that are not contraindicated for disease B seemed to be nearest to tissues preferentially affiliated with disease B. This could indicate that these tissues could be equally important to the pathophysiology of disease A and its therapeutic alleviation (as they might be to these same aspects of disease B), despite showing a high enrichment for genes harbouring disease B-associated variants. For example, the adrenal gland was detected as a tissue highly specific to depression by TSEA-DB. In our analysis, this tissue appeared to be nearest to the DTN of anxiety drugs that were not contraindicated in depression (**Fig. 7**), indicating that targeting of the adrenal gland may be vital to treat anxiety without aggravating comorbid depressive symptoms. The adrenal gland is an organ in the endocrine system that secretes the cortisol hormone, following the activation of the hypothalamic-pituitary-adrenal (HPA) axis by psychological stressors [75]. Several studies support the role of the adrenal gland as a focal point for depression. The adrenal gland exhibits a 70% increase in its volume in depressed individuals before successful anti-depressant treatment as well as in comparison with their matched controls [76, 77]. The cortisol hormone secreted by the adrenal gland, upon stress-induced activation of the HPA axis, has been linked to depressive symptoms in humans and monkeys. Increased cortisol levels have been positively correlated with depressive behaviour in rhesus macaques [78]. Enhanced cortisol secretion has been observed in depressive individuals [79], and has been proposed to (a) increase susceptibility to depression [80] and (b) be correlated with the stress experienced by depressed individuals [81]. Hyperactivation of the HPA axis has been noted in generalized anxiety disorder [82]. Treatment with selective serotonin reuptake inhibitors (SSRIs) has been shown to reduce HPA hyperactivity in both depressed patients and patients with generalized anxiety disorder [83–85]. Therefore, it is possible that anti-anxiety drugs that do not aggravate depressive symptoms target the adrenal gland, which produces the cortisol hormone, an effector or ‘endpoint’ of the HPA axis that seems to be regulated in a similar manner in depression as well as anxiety. We performed comparative transcriptome analysis of disease-associated, tissue-associated and drug-induced gene expression profiles using the BaseSpace Correlation Engine to analyse this hypothesis. BaseSpace Correlation Engine software suite is a data analysis platform that is used to study the effect of diseases and drugs on publicly available gene expression data [86].

**Figure 7:**
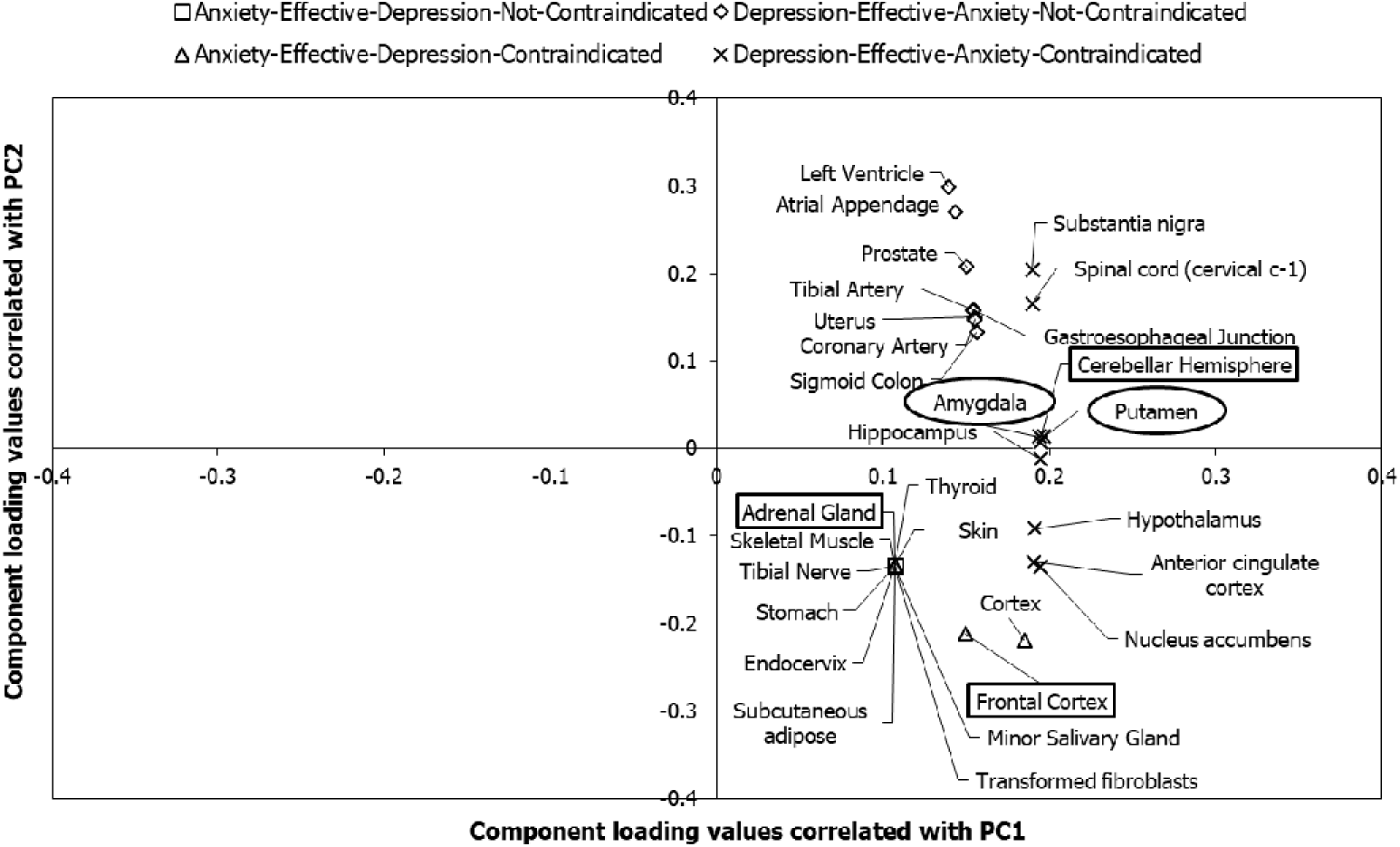
Tissues associated with the target networks of anxiety and depression drugs. The component loading values shown in the figure correspond to component scores of 4 DTNs of anxiety and depression along PC1 and PC2, which explain 90.7% and 6.5% of the total variance respectively. The tissues that were exclusively associated with each of the 4 DTNs among the top-ten tissues that were identified to be highly related to the DTNs, after computing the Euclidean distance between the component loading values and the component scores, are shown as square-shaped data points for the DTN of drugs effective in anxiety and not contraindicated in depression, diamond-shaped data points for the DTN of drugs effective in depression and not contraindicated in anxiety, triangle-shaped data points for the DTN of drugs effective in anxiety and contraindicated in depression and cross-mark-shaped data points for the DTN of drugs effective in depression and contraindicated in anxiety. The tissues shown in circular and rectangular boxes were also identified to be highly specific to anxiety and depression respectively by TSEA-DB (due to a significant enrichment of anxiety/depression-associated variants). Note that adrenal cortex, which was identified to be associated with anti-anxiety (disease A) drugs that are not contraindicated in depression (disease B), is a tissue enriched with depression (i.e. disease B) associated variants. This corroborates our finding that disease A drugs that are not contraindicated in disease B are affiliated with disease B-specific tissues.

As mentioned in the previous section, maprotiline was found among our list of anti-anxiety drugs that are not contraindicated in depression; clinical data supports its utility in the treatment of anxiety symptoms associated with depression [66]. The differential gene expression (DGE) profile induced by maprotiline (12.8 μM) in PC3 cells (Broad Connectivity Map (CMAP 2.0) [39]) was negatively correlated with the profile identified in the blood samples of patients with major depressive disorder patients (MDD) with generalized anxiety disorder (GAD) versus MDD patients without GAD (GSE98793 [40]) (**Fig. 8a**). This negative correlation of maprotiline with MDD/GAD could illustrate the fact that drugs administered to treat diseases often revert the expression of perturbed disease-associated genes to their normal levels [87, 88]. Secondly, the MDD/GAD profile was negatively correlated with the expression profile of adrenal gland cortex (**Fig. 8a**), indicating that this tissue could be critical to disease alleviation. Maprotiline-induced DGE profile was positively correlated with the profile of adrenal gland (**Fig. 8a**), indicating that maprotiline-mediated MDD/GAD alleviation may be dependent on adrenal gland, i.e. the reversal of MDD/GAD-associated expression profile induced by maprotiline could occur in the adrenal cortex. We then asked whether the genes differentially expressed in each of these profiles converged on a common set of biological processes. Specifically, we identified the top-10 Gene Ontology (GO) biological processes enriched among the genes differentially expressed in (i) MDD/GAD versus maprotiline (in different directions), (ii) MDD/GAD versus adrenal cortex (in different directions) and (iii) maprotiline versus adrenal cortex (in the same direction). We then used the web-based tool called NaviGO [89] to group these 30 enriched biological processes into functionally cohesive networks based on semantic similarity measures of GO terms. Two such functional networks not only had top-scoring edges between the GO terms, but also contained GO terms enriched among all the three differential expression profiles (**Fig. 8b, c**). One network contained four GO terms associated with protein folding (**Fig. 8b**), and another network contained eleven GO terms representing cell cycle events (**Fig. 8c**). Interestingly, ‘protein folding’, ‘cyclin D associated events in G1’ and ‘G1 phase’ were independently retrieved among our top-10 Reactome pathways found to be nearest (in terms of Euclidean distance) to the DTN of antidepressants that are not contraindicated in anxiety. Together, these results suggest that adrenal cortex may be preferentially targeted by drugs such as maprotiline that produce beneficial effects in anxiety as well as in depression, and that their actions may converge on protein folding and cell cycle processes. Note that maprotiline has been discontinued from usage [69] and is only being cited here as a demonstrative example.

**Figure 8:**
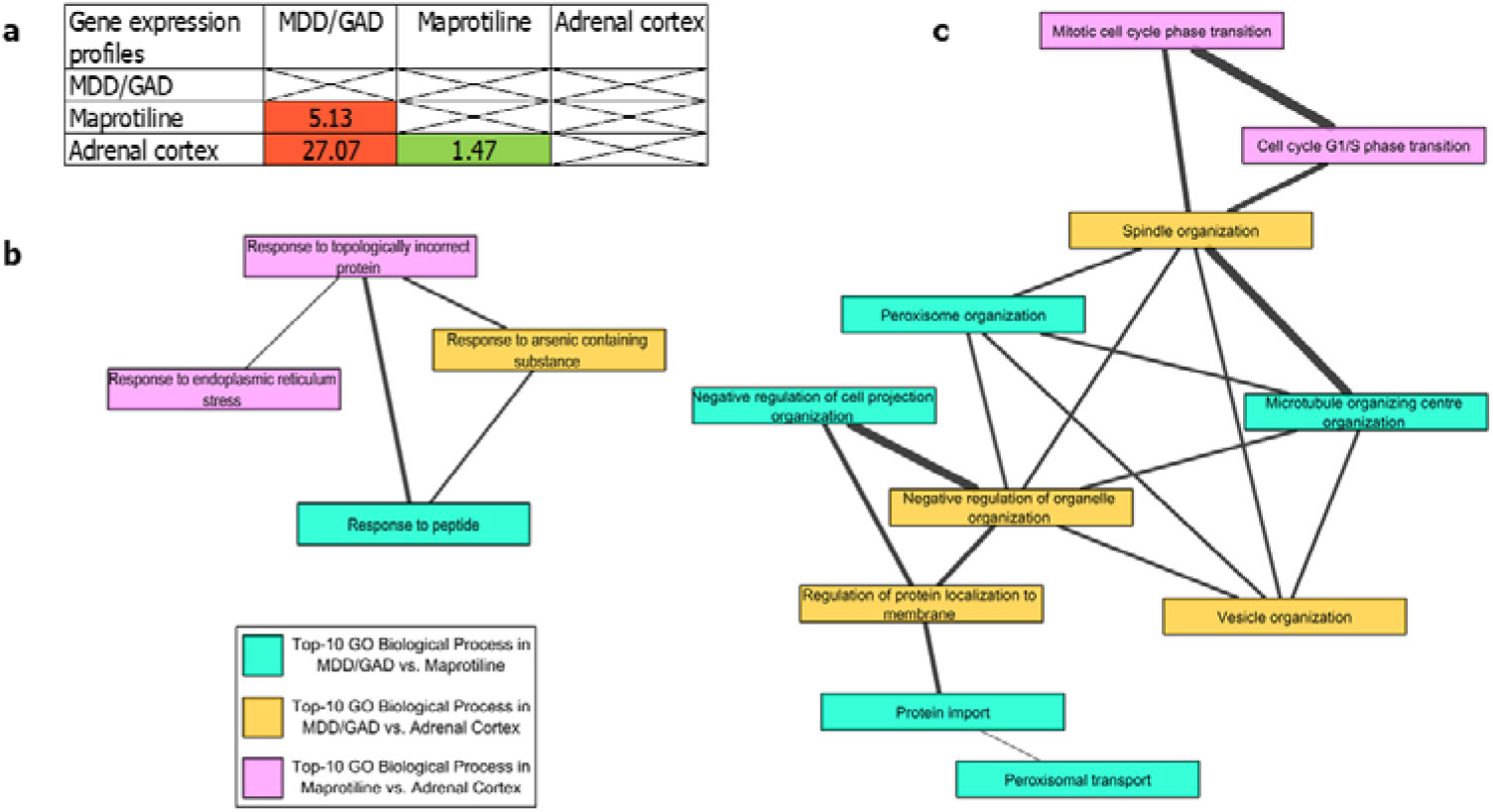
Relationship between MDD/GAD, maprotiline and adrenal cortex at transcriptomic and biological process levels. (a) Correlation of differential gene expression profiles associated with a comorbid condition (major depressive disorder and generalized anxiety disorder), a drug (maprotiline) and a tissue (adrenal cortex). −log_10_(p-values) indicating the overlap of the expression profiles have been shown; red and green colors indicate negative and positive correlations between the profiles respectively. Significant overlap was found among the genes that are upregulated in patients with both major depressive disorder (MDD) and generalized anxiety disorder (GAD) and downregulated on treating PC3 cells with maprotiline (p-value = 7.4E-06), among the genes that are upregulated in MDD/GAD patients and downregulated in adrenal cortex (p-value = 8.4E-28), and among the genes that are downregulated on treating PC3 cells with maprotiline and downregulated in adrenal cortex (p-value = 0.034). (b, c) The functional networks of the Gene Ontology biological processes related to (b) protein folding and (c) cell cycle events that were enriched in the three expression profiles. The GO terms associated with each of the expression profiles have been shown using different node colors. The thickness of the edges corresponds to the Resnik semantic similarity score for GO terms (greater the thickness of the edges, greater is the similarity between the linked GO terms).

On the other hand, the networks of disease A that are contraindicated for disease B seemed to be nearest to tissues preferentially affiliated with disease A. This could indicate that these disease A-specific tissues may play a role in producing beneficial effects in disease A, while producing deleterious effects in disease B. For example, spleen was detected as a tissue highly specific to rheumatoid arthritis. The primary functions of this lymphoid organ are blood filtration, recycling of iron from old blood cells and generation of adaptive immune responses against bacterial, fungal and viral infections [90]. However, spleen has also been shown to act as a reservoir of osteoclast precursor cells, which upon resorption into bones, differentiate into osteoclasts.[91] Splenomegaly (enlargement of the spleen) has been noted in 5-10% and 52% of rheumatoid arthritis patients in separate studies (based on physical examination and imaging studies respectively) [92–94]. Rheumatoid arthritis patients are also prone to developing spontaneous splenic ruptures [95]. In our analysis, spleen was identified to be nearest (in terms of Euclidean distance) to the DTN of rheumatoid arthritis drugs that were contraindicated in osteoporosis (**Fig. 9**). This seemed to indicate that spleen mediated opposite effects in rheumatoid arthritis and osteoporosis. Anecdotal evidence seemed to support this conjecture. While splenectomy seemed to improve rheumatoid arthritis in a patient [96], it seemed to inhibit (a) attenuation of osteoporosis in a rat model [97] and (b) fracture healing in patients [98].

**Figure 9:**
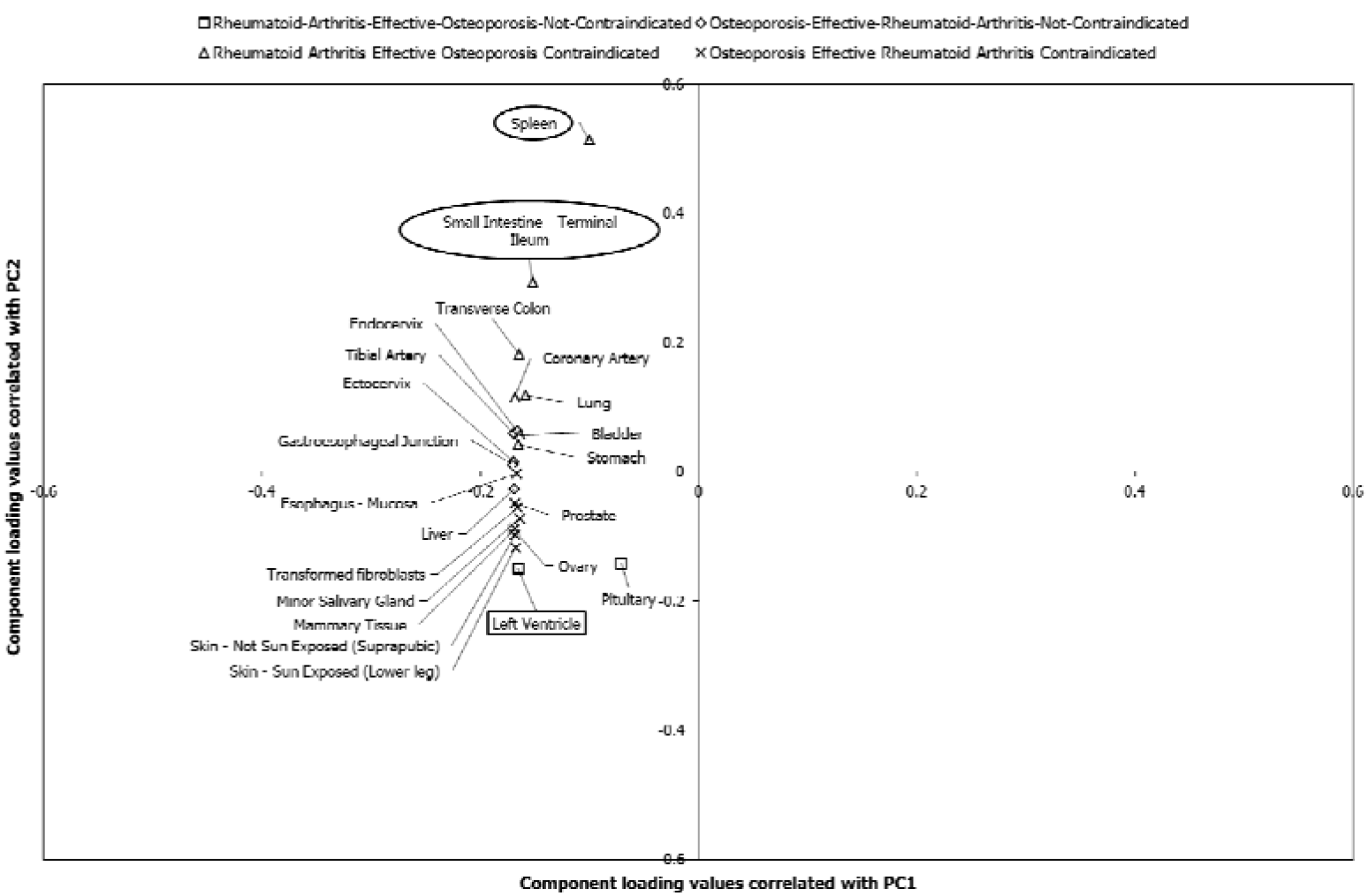
Tissues associated with the target networks of rheumatoid arthritis and osteoporosis drugs. Component loading values of 39 tissues associated with the drug target networks (DTNs) of rheumatoid arthritis (RA) and osteoporosis to PC1 and PC2 have been plotted along the X and Y axes respectively. PCA was performed with the p-values of enrichment of the tissues significantly associated (p-value < 0.05) with the DTNs of RA and osteoporosis. These values were transformed to −log_10_P values, which were then assembled into a data matrix containing tissues as rows and DTNs as columns. Unit variance scaling was applied across this matrix. Single value decomposition (SVD) with imputation was used to extract the principal components (PCs). The component loading values shown in the figure correspond to component scores of 4 DTNs along PC1 and PC2 that explain 89% and 6% of the total variance respectively. The tissues that were exclusively associated with each of the 4 DTNs among the top-ten tissues that were identified to be highly related to the DTNs, after computing the Euclidean distance between the component loading values and the component scores, are shown as square-shaped data points for the DTN of drugs effective in RA and not contraindicated in osteoporosis, diamond-shaped data points for the DTN of drugs effective in osteoporosis and not contraindicated in RA, triangle-shaped data points for the DTN of drugs effective in RA and contraindicated in osteoporosis and cross-mark-shaped data points for the DTN of drugs effective in osteoporosis and contraindicated in RA. The tissues shown in circular and rectangular boxes were also identified to be highly specific to RA and osteoporosis respectively by TSEA-DB (due to a significant enrichment of RA/osteoporosis-associated variants). Note that spleen, which was identified to be associated with rheumatoid arthritis (disease A) drugs that are contraindicated in osteoporosis (disease B), is a tissue enriched with rheumatoid arthritis (i.e. disease A) associated variants. This corroborates our finding that disease A drugs that are contraindicated in disease B are affiliated with disease A-specific

**Table 5** summarizes the general conclusions of our study. We discovered that the DTNs of disease A drugs that are not contraindicated for a disease B may be nearest (in terms of Euclidean distance) to (a) proteins that are either uniquely found in the PPI network of disease B or shared between the PPI networks of disease A and disease B, (b) biological pathways that are associated with B or are commonly active in both the diseases, and are regulated in the same direction in both the diseases and (c) tissues showing a high enrichment of disease-B associated variants and thereby preferential affiliation with the etiology of disease B, while also being important to the pathophysiology and treatment of disease A (**Table 5**). On the other hand, disease A drugs that are contraindicated for a disease B may be nearest to (a) proteins that are either uniquely found in the PPI network of disease A or are shared between the PPI networks of disease A and disease B, (b) biological pathways that are associated with disease A or are commonly active in both the diseases, and are regulated in an opposing manner in both the diseases and (c) tissues showing a high enrichment of disease A-associated variants and thereby preferential affiliation with the etiology of disease A, and mediating opposing effects in disease A and disease B (**Table 5**).

**Table 5:**
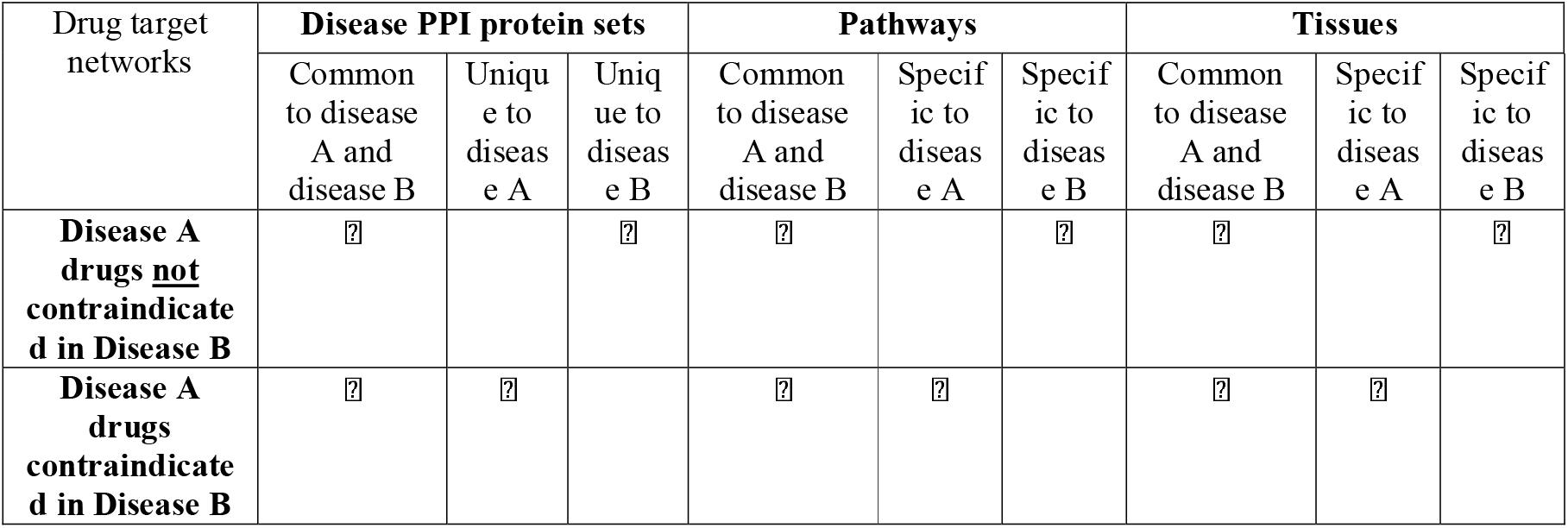
Disease network, pathway and tissue-level characterization of drugs that are contraindicated/not contraindicated in comorbid conditions. A ⍰ has been used to indicate the close affiliation of a specific category of drug target network with specific disease protein sets, disease-associated pathways and tissues.

## 3. Discussion

Despite the increased prevalence of adverse drug reactions in comorbidities, knowledge on the mechanistic basis of drug contraindications in such conditions is limited. In our study, we attempted to characterize the biological profiles of the target networks of drugs used in specific diseases that are either contraindicated or not contraindicated in a comorbid disease. We sought to provide an integrated interactome, pathway and tissue level view of the drug target networks.

The first key finding in our study was that the relative risk of comorbidity between diseases was proportional to their network similarity measures (**Fig. 2**). The four network similarity measures along with the relative risk were low in the case of our three negative control pairs, namely, Multiple sclerosis – Peroxisomal disorders, Schizophrenia – Rheumatoid arthritis, and Asthma – Schizophrenia. This confirmed that these were indeed non-comorbid pairs. The network similarity measures and relative risk were higher in the case of Anxiety – Depression, Asthma – Hypertension, Chronic obstructive pulmonary disorder – Heart failure, Type 2 diabetes – Obesity, Rheumatoid arthritis – Osteoporosis, and Parkinson’s disease – Schizophrenia, confirming that they were comorbidities. However, these measures do not follow the same trend in the case of the comorbid pairs. The higher relative risks of Rheumatoid arthritis – Osteoporosis and Parkinson’s disease – Schizophrenia (compared with the other comorbid pairs) were not accompanied by a corresponding increase in the network similarity measures. Several factors may explain these variations in our analysis. Firstly, it has been shown that relative risk overestimates the comorbid associations between rare diseases and underestimates the associations between highly prevalent diseases [43]. The number of cases in the HuDiNe database for Rheumatoid arthritis – Osteoporosis and Parkinson’s disease – Schizophrenia are 24629 and 5439 respectively, which can be classified as rare occurrences when compared with the other comorbid pairs. **Additional File 17: Figure S14** shows the relationship of the relative risks of the nine pairs of diseases with the individual prevalence of the diseases and the prevalence of the disease pairs as comorbidities. Secondly, the human interactome is a progressively developing network with ~85% remaining to be discovered. Therefore, the inherent incompleteness of the human PPI network, sampling biases introduced as a result of the selective discovery of PPIs, and the tendency of such incomplete networks to exhibit small overlaps [60] could have led to the underestimation of the network overlaps. Our second key finding was that druggable proteins were highly enriched among the proteins shared between the networks of two comorbid diseases (**Table 2**). Based on these results, we speculated that drug action on targets shared between the two diseases may give rise to contraindications in comorbidities. Interestingly, this hypothesis was only partially supported in our study.

The major finding in this respect was that the target network of the drugs used in the treatment of a specific disease A and contraindicated in a comorbid disease B showed preferential affiliation to proteins shared between the PPI networks of both the diseases or proteins uniquely found in the PPI network of the disease A, pathways shared by the two diseases or pathways associated with the disease A and tissues specifically associated with disease A (**Table 5**). As explained before, this was contrary to our hypothesis that these target networks would be preferentially affiliated with common mechanisms underlying the two diseases. This hypothesis was based on the assumption that adverse events stem from drugs inducing opposite pharmacological effects in comorbid diseases by targeting effectors that are shared between the two diseases. However, our findings indicate that mechanisms underlying the pathology of disease A may contribute to contraindications in the comorbid disease B. Although further studies are required to examine the basis of this finding, it seems to indicate that the possibility of contraindications may be high when disease A drugs are highly specific to disease A in terms of the targeted PPI network, pathway and tissue. Instead, rational drug development should take into account the causative and correlational influences of the other comorbid conditions (disease B) that co-exist with disease A.

The target network of the drugs used in the treatment of a specific disease A and not contraindicated in a comorbid disease B showed preferential affiliation to proteins shared between the PPI networks of both the diseases or proteins uniquely found in the PPI network of the comorbid disease B, pathways shared between the two diseases or pathways associated with the comorbid disease B and tissues specifically associated with the comorbid disease B (**Table 5**). This was contrary to our expectation that these target networks would be preferentially affiliated with biological modalities pertaining to disease A. This conjecture was based on the assumption that for a drug to be specifically active against a specific disease A without aggravating a comorbid disease B, it had to reverse the phenotypes specifically associated with disease A. In this model, phenotypes of disease B were considered as ‘off-targets’ in line with the principles of conventional pharmacology, in which unintended effects of the drugs were attributed to interaction with pathways that may not be consequential to the pathology of disease A (i.e. pathways relevant to disease B) [13]. Our findings on the contrary indicate that the mechanisms underlying the pathology of the comorbid disease B may contribute to the therapeutic alleviation of disease A. Although further investigations may be necessary to dissect the basis for this observation, it is possible that an etiological association between the two diseases may cause their emergence or development to be interdependent. Specifically, future studies should concentrate on 3 etiological models of comorbidity [99], namely, the direct causation model, the associated risk factors model and the heterogeneity model. Disease B could be directly responsible for causing disease A in the ‘disease causation model’. The comorbidity of disease A and disease B may arise from the correlation of the risk factors of disease B with the risk factors of disease A in the ‘associated risk factors model’. On the other hand, comorbidity in the ‘heterogeneity model’ may arise not from the correlation of the risk factors associated with disease A and disease B, but from the capacity of the risk factors of disease A to cause disease B and vice versa. On applying the disease causation model to our findings, one may speculate that drugs targeting the proteins uniquely found in the disease B PPI network, and the pathways and tissues associated with disease B may alleviate disease A without aggravating disease B. The associated risk factors and heterogeneity models in this scenario would imply that the risk factors of disease B would influence the development of disease A directly, or through correlation with the risk factors of disease A. This model can be illustrated for genetic risk factors of disease B with the capacity to influence disease A. For example, the alterations in such genes would have led to pathway perturbations in specific tissues, which if counteracted by the drugs, may lead to alleviation of disease A.

Despite disease A drugs contraindicated in disease B and disease A drugs not contraindicated in disease B showing preferential affiliation with disease A and disease B respectively, it was clear, at least in the case of the drug target and disease network analysis, that both these categories also showed affiliation with proteins shared between the two diseases (**Table 5**). This is in line with the speculation that both beneficial and adverse outcomes of drug treatment may arise from shared effectors and pathways, and that it may be difficult to delineate the separate mechanisms underlying the two outcomes [13]. Future analysis should focus on biological variables with the potential to differentially affect the functions of such shared proteins, specifically their cellular, pathway and tissue landscapes.

Our current approach has some limitations. Firstly, our study is based on 6 pairs of diseases that were selected based on literature survey. Ideally, future studies must be expanded to include all the known pairs of comorbid disorders. Secondly, our analysis did not take the overlaps among the drug target networks into account; this would have allowed us to identify the network configurations of disease A – disease B – disease A drug not contraindicated in disease B – disease B drug not contraindicated in disease A. Secondly, although we were able to support our findings by citing evidence based on the known clinical activity of specific drugs, further investigations with the six comorbid disease pairs are essential to confirm the validity of our findings. These should focus on large-scale analysis of patient treatment data collected from observational studies and functional assays in animal models of human comorbidities.

In summary, our findings suggest that studies driven by biological modalities that influence comorbidities, such as disease PPI networks, pathways and tissue-specificity, are essential for rational drug development and minimization of adverse events. The results from our study have therapeutic applications and may directly benefit future assessments of drug contraindications in individuals with comorbidities.

## 4. Conclusions

We observed that the target networks of disease A drugs that were not contraindicated in disease B were mostly affiliated with the disease B network, and pathways and tissues associated with disease B. On the other hand, the target networks of disease A drugs that were contraindicated in disease B were affiliated with the disease A network, and pathways and tissues associated with disease A. This could indicate that etiological associations between the two diseases could play an active role in their therapeutic alleviation. In summary, our findings suggest that the enrichment patterns of drug target networks in pathways, tissues and the PPI networks of comorbid diseases will help identify drugs with/without contraindications in comorbidities.

## Supporting information

Combined Supplementary File

## 3. Acknowledgments

## Funding

This work was partially supported by INSA Senior Scientist grant of Prof. N. Balakrishnan.

## Author contributions

NBK conceived and designed the research. KBK designed and performed the analyses. NBK and MKG supervised the interactome-based analyses, and SKB and SJ provided scientific inputs on the biological aspects of the study. KBK prepared the manuscript and NBK, SKB, MKG and SJ edited the manuscript through extensive mutual consultations.

## Competing interests

None

## Data availability

The condition concept names from the TWOSIDES database that were used to categorize the drugs associated with each of the comorbid disease pairs, the drug lists, the proteins targeted by the drugs, and the genes associated with the comorbid disease pairs have been made available as **Table S1**, **Table S2**, **Table S3** and **Table S4**, respectively.

## Notes

### Competing Interest Statement

The authors have declared no competing interest.

