## Supplementary material for "Drug Contraindications in Comorbid Diseases: a Protein Interactome Perspective": Combined Supplementary File

**Table S1.** Condition concept names in the TWOSIDES database used to compile the 4 drug groups for each of the disease pairs

| **Anxiety-Depression** | |
| --- | --- |
| Anxiety: condition_concept_name | Depression: condition_concept_name |
| Anxiety | Depression |
| Anxiety disorder | Major depression |
| Generalised anxiety disorder | Depressive symptom |
| Adjustment disorder with anxiety | Depression suicidal |
| Adjustment disorder with mixed anxiety and depressed mood | Depression postoperative |
|  | Postpartum depression |
|  | Depressive delusion |
|  | Agitated depression |

| **Type 2 diabetes-Obesity** | |
| --- | --- |
| Type 2 diabetes: condition_concept_name | Obesity: condition_concept_name |
| Type 2 diabetes mellitus | Obesity |
| Diabetes mellitus inadequate control | Central obesity |
| Diabetes mellitus non-insulin-dependent |  |
| Insulin-requiring type 2 diabetes mellitus |  |
| Insulin resistant diabetes |  |

| **Rheumatoid arthritis-Osteoporosis** | |
| --- | --- |
| Rheumatoid arthritis: condition_concept_name | Osteoporosis: condition_concept_name |
| Rheumatoid arthritis | Osteoporosis |

| **Parkinson's disease-Schizophrenia** | |
| --- | --- |
| Parkinson's disease: condition_concept_name | Schizophrenia: condition_concept_name |
| Parkinson's disease | Schizophrenia |
| Parkinsonism | Schizophrenia, paranoid type |
| Parkinsonian rest tremor | Schizophreniform disorder |
| Parkinsonian gait | Schizophrenia, disorganised type |
|  | Schizophrenia, undifferentiated type |

| **Asthma-Hypertension** | |
| --- | --- |
| Asthma: condition_concept_name | Hypertension: condition_concept_name |
| Asthma | Hypertension |
| Cardiac asthma | Pulmonary arterial hypertension |
| Status asthmaticus | Pulmonary hypertension |
| Asthmatic crisis | Essential hypertension |
| Analgesic asthma syndrome | Portal hypertension |
| Asthma exercise induced | Benign intracranial hypertension |
|  | Malignant hypertension |
|  | Labile hypertension |
|  | Ocular hypertension |
|  | Systolic hypertension |
|  | Procedural hypertension |
|  | Renal hypertension |
|  | Renovascular hypertension |
|  | Accelerated hypertension |
|  | Orthostatic hypertension |
|  | Gestational hypertension |
|  | Diastolic hypertension |
|  | Secondary hypertension |
|  | Congenital pulmonary hypertension |
|  | Pregnancy induced hypertension |

| **Chronic obstructive pulmonary disease-Heart failure** | |
| --- | --- |
| Chronic obstructive pulmonary disease: condition_concept_name | Heart failure: condition_concept_name |
| Chronic obstructive pulmonary disease | Cardiac failure |
| Chronic obstructive airways disease | Cardiac failure congestive |
| Chronic obstructive airways disease exacerbated | Cardiac failure acute |
|  | Cardiac failure chronic |
|  | Cardiac failure high output |

**Table S2.** Drugs belonging to the 4 groups in each disease pair compiled based on drug activity in the two diseases. The drugs were compiled from Drug Bank and categorized into 4 groups based on their associations with adverse events documented in the TWOSIDES database.

| **Anxiety-Depression** | | | |
| --- | --- | --- | --- |
| Anxiety drugs not contraindicated in depression | Depression drugs not contraindicated in anxiety | Anxiety drugs contraindicated in depression | Depression drugs contraindicated in anxiety |
| Acetylsalicylic acid | Clomipramine | Hydroxyzine | Buspirone |
| Bromopride | Desipramine | Diazepam | nefazodone |
| Delorazepam | Ephedrine | Oxazepam | Citalopram |
| Etifoxine | Flupentixol | Prazepam | Amitriptyline |
| Glycyrrhiza glabra | Fluphenazine | Lorazepam | Amphetamine |
| Isopropamide | Ginkgo biloba | Oxymorphone |  |
| Ketazolam | Isocarboxazid | Trazodone |  |
| Maprotiline | Lithium cation | Phenobarbital |  |
| Methotrimeprazine | Mianserin | clotiazepam |  |
| Mexazolam | Mirtazapine | Flunitrazepam |  |
| Otilonium | Pindolol | Trimebutine |  |
| Propantheline | Pipradrol |  |  |
|  | Protriptyline |  |  |
|  | Trimipramine |  |  |
|  | Tryptophan |  |  |
|  | Valpromide |  |  |

| **Type 2 diabetes-Obesity** | | | |
| --- | --- | --- | --- |
| Type 2 diabetes drugs not contraindicated in BMI >30 kg/m2 | BMI >30 kg/m2 drugs not contraindicated in type 2 diabetes | Type 2 diabetes drugs contraindicated in BMI >30 kg/m2 | BMI >30 kg/m2 drugs contraindicated in type 2 diabetes |
| Acarbose | Benzphetamine | glimepiride | Bupropion |
| Albiglutide | Lecithin, soybean | exenatide | Fluoxetine |
| Alogliptin | Lorcaserin | rosiglitazone | Phentermine |
| Bromocriptine | Naltrexone | Metformin | orlistat |
| Canagliflozin | Pipradrol | Simvastatin | Amphetamine |
| Chlorpropamide |  | atorvastatin | sibutramine |
| Colesevelam |  | insulin detemir |  |
| Dapagliflozin |  | Glyburide |  |
| Dulaglutide |  | sitagliptin |  |
| Empagliflozin |  | pioglitazone |  |
| Ertugliflozin |  | topiramate |  |
| Gemigliptin |  | Insulin Lispro |  |
| Gliclazide |  | Glucagon |  |
| Gliquidone |  | Pramlintide |  |
| Insulin degludec |  | repaglinide |  |
| Lipoic acid |  | Insulin Glargine |  |
| Lixisenatide |  | nateglinide |  |
| Luseogliflozin |  | Linagliptin |  |
| Miglitol |  | Glipizide |  |
| Mitiglinide |  |  |  |
| Saxagliptin |  |  |  |
| Semaglutide |  |  |  |
| Teneligliptin |  |  |  |
| Tolazamide |  |  |  |
| Tolbutamide |  |  |  |
| Trelagliptin |  |  |  |
| Vildagliptin |  |  |  |
| Voglibose |  |  |  |

| **Rheumatoid arthritis-Osteoporosis** | | | |
| --- | --- | --- | --- |
| Rheumatoid arthritis drugs not contraindicated in osteoporosis | Osteoporosis drugs not contraindicated in rheumatoid arthritis | Rheumatoid arthritis drugs contraindicated in osteoporosis | Osteoporosis drugs contraindicated in rheumatoid arthritis |
| Aceclofenac | Alendronic acid | Ibuprofen | Lidocaine |
| Acetylsalicylic acid | Boron | Diclofenac | Estradiol |
| Auranofin | Calcium Phosphate | Hydrocortisone | Calcium |
| Azapropazone | Calcium lactate gluconate | Dexamethasone | Calcium Carbonate |
| Betamethasone phosphate | Conjugated estrogens | Acetaminophen | Teriparatide |
| Bupivacaine | Cyproterone acetate | Naproxen | Calcium Citrate |
| Chloroquine | Genistein | Prednisone | zoledronic acid |
| Choline magnesium trisalicylate | L-Glutamine | infliximab | Calcium Gluconate |
| Corticotropin | Medroxyprogesterone acetate | Methylprednisolone | Ibandronate |
| Cortisone acetate | Menatetrenone | Sulfasalazine | Cholecalciferol |
| Dexketoprofen | Norethisterone | prednisolone | salmon calcitonin |
| Diflunisal | Romosozumab | meloxicam | Raloxifene |
| Etofenamate |  | Ranitidine | risedronic acid |
| Etoricoxib |  | Tolmetin |  |
| Fenbufen |  | valdecoxib |  |
| Fenoprofen |  | Etanercept |  |
| Flurbiprofen |  | Omeprazole |  |
| Hydrocortisone acetate |  | Tacrolimus |  |
| Ketoprofen |  | oxaprozin |  |
| Lornoxicam |  | nabumetone |  |
| Meclofenamic acid |  | Triamcinolone |  |
| Methylprednisolone hemisuccinate | denosumab |  |  |
| Piroxicam cinnamate |  | Betamethasone |  |
| Proglumetacin |  | celecoxib |  |
| Sodium aurothiomalate |  | Piroxicam |  |
| Sulindac |  | leflunomide |  |
| Tenoxicam |  | Hydroxychloroquine |  |
| Tetracosactide |  | Etodolac |  |
| Thiocolchicoside |  | Azathioprine |  |
| Tiaprofenic acid |  |  |  |

| **Chronic obstructive pulmonary disease-Heart failure** | | | |
| --- | --- | --- | --- |
| Chronic obstructive pulmonary disease drugs not contraindicated in heart failure | Heart failure drugs not contraindicated in chronic obstructive pulmonary disease | Chronic obstructive pulmonary disease drugs contraindicated in heart failure | Heart failure drugs contraindicated in chronic obstructive pulmonary disease |
| Aclidinium | Enalaprilat | Arformoterol | Atenolol |
| Beclomethasone dipropionate |  | Budesonide | Benazepril |
| Carmoterol |  | Cefpodoxime | Bisoprolol |
| Ceftibuten |  | Ciclesonide | Bumetanide |
| Clenbuterol |  | Formoterol | Captopril |
| Doxapram |  | Ipratropium | Fosinopril |
| Doxofylline |  | Roflumilast | Furosemide |
| Erdosteine |  | Salmeterol | Hydralazine |
| Fluticasone furoate |  | Terbutaline | Isosorbide dinitrate |
| Fluticasone propionate |  | Theophylline | Losartan |
| Glycopyrronium |  | Tiotropium | Ramipril |
| Indacaterol |  | Tulobuterol | Spironolactone |
| Lactose |  |  | Telmisartan |
| Levosalbutamol |  |  | Trandolapril |
| Olodaterol |  |  |  |
| Oxtriphylline |  |  |  |
| Revefenacin |  |  |  |
| Rufloxacin |  |  |  |
| Salbutamol |  |  |  |
| Umeclidinium |  |  |  |
| Vilanterol |  |  |  |

| **Parkinson's disease-Schizophrenia** | | | |
| --- | --- | --- | --- |
| Parkinson's disease drugs not contraindicated in schizophrenia | Schizophrenia drugs not contraindicated in Parkinson's disease | Parkinson's disease drugs contraindicated in schizophrenia | Schizophrenia drugs contraindicated in Parkinson's disease |
| Amantadine | Amisulpride | Levodopa | Amitriptyline |
| Apomorphine | Aripiprazole lauroxil | ropinirole | aripiprazole |
| Benserazide | Asenapine | Biperiden | Chlorpromazine |
| Bornaprine | Brexpiprazole | Selegiline | Clozapine |
| Bromocriptine | Cariprazine | entacapone | Haloperidol |
| Carbidopa | Chlorprothixene |  | lurasidone |
| Droxidopa | Fluspirilene |  | Mirtazapine |
| Istradefylline | Iloperidone |  | olanzapine |
| Melevodopa | Loxapine |  | Prochlorperazine |
| Opicapone | Lumateperone |  | Promazine |
| Pergolide | Methotrimeprazine |  | Risperidone |
| Piribedil | Molindone |  | Sulpiride |
| Pramipexole | Paliperidone |  | Trazodone |
| Profenamine | Perphenazine |  | Trifluoperazine |
| Rasagiline | Pipotiazine |  | ziprasidone |
| Rotigotine | Sertindole |  |  |
| Safinamide | Thioproperazine |  |  |
| Tolcapone | Thioridazine |  |  |
|  | Thiothixene |  |  |
|  | Zuclopenthixol |  |  |

| **Asthma-Hypertension** | | | |
| --- | --- | --- | --- |
| Asthma drugs not contraindicated in hypertension | Hypertension drugs not contraindicated in asthma | Asthma drugs contraindicated in hypertension | Hypertension drugs contraindicated in asthma |
| Beclomethasone dipropionate | Acebutolol | fluticasone | Metoprolol |
| Carmoterol | Ammonium chloride | montelukast | Indapamide |
| Cortisone acetate | Azilsartan medoxomil | Hydrocortisone | Hydrochlorothiazide |
| Cromoglicic acid | Bendroflumethiazide | Budesonide | Losartan |
| Etafedrine | Benidipine | Dexamethasone | rosuvastatin |
| Fluticasone furoate | Candesartan cilexetil | flunisolide | Amlodipine |
| Fluticasone propionate | Canrenoic acid | salmeterol | Triamterene |
| Indacaterol | Chlorothiazide | Methylprednisolone | Ramipril |
| Lactose | Cilazapril | Terbutaline | quinapril |
| Methoxyphenamine | Cilnidipine | Epinephrine | Captopril |
| Mometasone furoate | Clevidipine | zafirlukast | Felodipine |
| Nedocromil | Clopamide | Prednisone | topiramate |
| Orciprenaline | Delapril | Aminophylline | Lisinopril |
| Oxtriphylline | Enalaprilat | Ipratropium | Propranolol |
| Pranlukast | Eprosartan | formoterol | candesartan |
| Racepinephrine | Guanfacine | Triamcinolone | olmesartan |
| Reproterol | Hydroflumethiazide | Guaifenesin | nebivolol |
| Salbutamol | Lacidipine | Mometasone | Furosemide |
| Vilanterol | Lercanidipine | Chlorpheniramine | irbesartan |
| Zileuton | Levamlodipine | desloratadine | Enalapril |
|  | Methyclothiazide | arformoterol | Fosinopril |
|  | Moexipril | tiotropium | telmisartan |
|  | Moxonidine | Theophylline | Spironolactone |
|  | Nitrendipine | ciclesonide | atorvastatin |
|  | Pindolol | sodium citrate | Perindopril |
|  | Piretanide |  | Clonidine |
|  | Polythiazide |  | aliskiren |
|  | Prazosin |  | urapidil |
|  | Reserpine |  | Isosorbide Dinitrate |
|  | Rilmenidine |  | Bisoprolol |
|  | Torasemide |  | valsartan |
|  | Trandolapril |  | Nadolol |
|  | Xipamide |  | Amiloride |
|  | Zofenopril |  | Labetalol |
|  |  |  | Diltiazem |
|  |  |  | Terazosin |
|  |  |  | benazepril |
|  |  |  | Atenolol |
|  |  |  | Doxazosin |
|  |  |  | Verapamil |
|  |  |  | Nifedipine |
|  |  |  | Isradipine |
|  |  |  | Methyldopa |
|  |  |  | carvedilol |
|  |  |  | Betaxolol |
|  |  |  | eplerenone |
|  |  |  | Celiprolol |
|  |  |  | Nicardipine |
|  |  |  | Nisoldipine |

| **Schizophrenia-Asthma** | | | |
| --- | --- | --- | --- |
| Schizophrenia not contraindicated in asthma | Schizophrenia contraindicated in asthma | Asthma drugs not contraindicated in schizophrenia | Asthma drugs contraindicated in schizophrenia |
| Aripiprazole lauroxil | Amitriptyline | Aminophylline | Chlorpheniramine |
| Asenapine | Aripiprazole | Arformoterol | Dexamethasone |
| Brexpiprazole | Chlorpromazine | Beclomethasone dipropionate | Fluticasone |
| Cariprazine | Clozapine | Budesonide | Guaifenesin |
| Chlorprothixene | Haloperidol | Carmoterol | Hydrocortisone |
| Fluspirilene | Lurasidone | Ciclesonide | Methylprednisolone |
| Iloperidone | Olanzapine | Cortisone acetate | Montelukast |
| Loxapine | Paliperidone | Cromoglicic acid | Prednisone |
| Lumateperone | Perphenazine | Desloratadine | Tiotropium |
| Methotrimeprazine | Prochlorperazine | Epinephrine | Triamcinolone |
| Molindone | Quetiapine | Etafedrine | Zafirlukast |
| Promazine | Risperidone | Flunisolide |  |
| Samidorphan | Thioridazine | Fluticasone furoate |  |
| Sertindole | Thiothixene | Fluticasone propionate |  |
| Sulpiride | Trazodone | Formoterol |  |
| Thioproperazine | Trifluoperazine | Indacaterol |  |
| Zuclopenthixol | Ziprasidone | Ipratropium |  |
|  |  | Lactose |  |
|  |  | Methoxyphenamine |  |
|  |  | Mometasone |  |
|  |  | Mometasone furoate |  |
|  |  | Nedocromil |  |
|  |  | Orciprenaline |  |
|  |  | Oxtriphylline |  |
|  |  | Pranlukast |  |
|  |  | Racepinephrine |  |
|  |  | Reproterol |  |
|  |  | Salbutamol |  |
|  |  | Salmeterol |  |
|  |  | Sodium citrate |  |
|  |  | Terbutaline |  |
|  |  | Theophylline |  |
|  |  | Vilanterol |  |
|  |  | Zileuton |  |

| **Schizophrenia-Rheumatoid arthritis** | | | |
| --- | --- | --- | --- |
| Schizophrenia not contraindicated in rheumatoid arthritis | Schizophrenia contraindicated in rheumatoid arthritis | Rheumatoid arthritis drugs not contraindicated in schizophrenia | Rheumatoid arthritis drugs contraindicated in schizophrenia |
| Aripiprazole lauroxil | Amitriptyline | Aceclofenac | Acetaminophen |
| Asenapine | Aripiprazole | Acetylsalicylic acid | Betamethasone |
| Brexpiprazole | Chlorpromazine | Auranofin | Celecoxib |
| Cariprazine | Clozapine | Azapropazone | Dexamethasone |
| Chlorprothixene | Haloperidol | Azathioprine | Diclofenac |
| Fluspirilene | Olanzapine | Betamethasone phosphate | Etanercept |
| Iloperidone | Prochlorperazine | Bupivacaine | Etodolac |
| Loxapine | Quetiapine | Chloroquine | Hydrocortisone |
| Lumateperone | Risperidone | Choline magnesium trisalicylate | Ibuprofen |
| Methotrimeprazine | Thioridazine | Corticotropin | Meloxicam |
| Molindone | Thiothixene | Cortisone acetate | Methylprednisolone |
| Promazine | Trazodone | Denosumab | Nabumetone |
| Samidorphan | Ziprasidone | Dexketoprofen | Naproxen |
| Sertindole |  | Diflunisal | Omeprazole |
| Sulpiride |  | Etofenamate | Oxaprozin |
| Thioproperazine |  | Etoricoxib | Piroxicam |
| Zuclopenthixol |  | Fenbufen | Prednisone |
| Lurasidone |  | Fenoprofen | Ranitidine |
| Paliperidone |  | Flurbiprofen | Triamcinolone |
| Perphenazine |  | Hydrocortisone acetate |  |
| Trifluoperazine |  | Hydrocortisone succinate |  |
|  |  | Hydroxychloroquine |  |
|  |  | Ketoprofen |  |
|  |  | Leflunomide |  |
|  |  | Lornoxicam |  |
|  |  | Meclofenamic acid |  |
|  |  | Methylprednisolone hemisuccinate | |
|  |  | Niflumic acid |  |
|  |  | Piroxicam cinnamate |  |
|  |  | Prednisolone |  |
|  |  | Proglumetacin |  |
|  |  | Sodium aurothiomalate |  |
|  |  | Sulfasalazine |  |
|  |  | Sulindac |  |
|  |  | Tacrolimus |  |
|  |  | Tenoxicam |  |
|  |  | Tetracosactide |  |
|  |  | Thiocolchicoside |  |
|  |  | Tiaprofenic acid |  |
|  |  | Tolmetin |  |
|  |  | Valdecoxib |  |

| **Multiple sclerosis-Peroxisomal disorders** | |
| --- | --- |
| Multiple sclerosis drugs | Peroxisomal disorder drugs |
| Amantadine | Cholic acid |
| Azathioprine |  |
| Baclofen |  |
| Cannabidiol |  |
| Dalfampridine |  |
| Dantrolene |  |
| Dimethyl fumarate |  |
| Glatiramer |  |
| Methotrexate |  |
| Modafinil |  |
| Ozanimod |  |
| Peginterferon beta-1a |  |
| Teriflunomide |  |

**Table S3.** Proteins targeted by the drugs in each of the 4 groups for each of the disease pairs. The proteins targets of each drug group were compiled by querying the Drug Bank database through DGIdb (drug gene interaction database).

| **Anxiety-Depression** | | | |
| --- | --- | --- | --- |
| Anxiety drugs not contraindicated in depression | Depression drugs not contraindicated in anxiety | Anxiety drugs contraindicated in depression | Depression drugs contraindicated in anxiety |
| DRD2 | HTR2C | HRH1 | DRD2 |
| GABRA1 | HTR2B | KCNH2 | HTR1A |
| CHRM4 | HTR2A | GABRA1 | DRD3 |
| CHRM3 | SLC6A4 | OPRM1 | ADRA1A |
| ADRA2A | GSTP1 | OPRD1 | DRD4 |
| HTR7 | SLC6A2 | HTR1A | ADRA2A |
| SLC6A2 | SMPD1 | SLC6A4 | HTR2A |
| ADRA1A | ADRA1A | ADRA2A | SLC6A4 |
| HRH1 | CHRM1 | ADRA1A | SLC6A3 |
| CHRM5 | DRD2 | HTR2A | HTR2C |
| HTR2A | HRH1 | HTR2C | SLC6A2 |
| CHRM1 | HTR1A | GRIA2 | KCNH2 |
| CHRM2 | ADRA2A | CHRNA4 | ADRA1B |
| HTR2C | ADRB2 | CHRNA7 | HRH1 |
| DRD4 | CHRM5 | GRIN1 | HTR1D |
| DRD1 | CHRM2 | GRIK2 | HTR6 |
| ADRA2B | CHRM3 | NR1I2 | HTR7 |
| ADRA2C | CHRM4 | CHRM4 | HRH2 |
| DRD5 | ADRB1 | CHRM2 | CHRM1 |
| ADRA1D | DRD1 | CHRM1 | OPRD1 |
| DRD3 | CALM1 | CHRM3 | SIGMAR1 |
| ADRA1B | AR | KCNMA1 | HTR1B |
|  | MAOA | CACNA1G | KCNA1 |
|  | MAOB | CACNA1C | OPRK1 |
|  | OPRK1 |  | OPRM1 |
|  | SLC6A3 |  | NTRK1 |
|  | ADRA2B |  | HRH4 |
|  | HTR7 |  | ADRA1D |
|  | DRD3 |  | KCNQ2 |
|  | HTR6 |  | NTRK2 |
|  | HTR1F |  | KCNQ3 |
|  | HRH4 |  | MAOA |
|  | ADRA2C |  | CARTPT |
|  | ADRB3 |  | ADRB1 |
|  | HTR1B |  | SLC18A2 |
|  | HTR1D |  | TAAR1 |
|  | HTR3A |  | MAOB |
|  | ADRA1B |  |  |
|  | WARS |  |  |
|  | WARS2 |  |  |

| **Parkinson's disease-Schizophrenia** | | | |
| --- | --- | --- | --- |
| Parkinson's disease drugs not contraindicated in schizophrenia | Schizophrenia drugs not contraindicated in Parkinson's disease | Parkinson's disease drugs contraindicated in schizophrenia | Schizophrenia drugs contraindicated in Parkinson's disease |
| DRD2 | HTR2A | DRD3 | HTR1D |
| GRIN3A | HTR7 | DRD2 | HTR6 |
| CHRNA4 | DRD2 | DRD5 | HTR7 |
| CHRNA7 | DRD3 | DRD1 | HRH2 |
| CHRNA3 | DRD4 | DRD4 | SLC6A4 |
| DRD5 | HTR2B | ADRA1A | CHRM1 |
| HTR1A | HTR2C | CHRM1 | OPRD1 |
| ADRA2A | ADRA2C | MAOA | ADRA1B |
| HTR1B | HRH1 | MAOB | SLC6A2 |
| HTR2B | HTR1B | COMT | SIGMAR1 |
| HTR2C | HTR6 |  | HTR1B |
| ADRA2B | HTR1A |  | ADRA1A |
| DRD1 | ADRB2 |  | KCNA1 |
| HTR2A | HRH2 |  | HTR1A |
| ADRA2C | ADRA2B |  | HRH1 |
| HTR1D | ADRA2A |  | OPRK1 |
| DRD4 | ADRB1 |  | OPRM1 |
| DRD3 | ADRA1A |  | NTRK1 |
| DDC | DRD1 |  | HRH4 |
| ADRA1D | HTR5A |  | ADRA1D |
| HTR7 | ADRA1B |  | HTR2C |
| ADRA1A | CHRM2 |  | HTR2A |
| ADRA1B | CHRM1 |  | ADRA2A |
| ADRB2 | CHRM4 |  | KCNQ2 |
| PAH | CHRM3 |  | NTRK2 |
| ADRB1 | CHRM5 |  | KCNH2 |
| ADRB3 | CACNG1 |  | KCNQ3 |
| ADORA2A | SLC6A2 |  | CHRM3 |
| ADORA1 | HTR1E |  | HTR3A |
| COMT | HTR3A |  | HTR1E |
| CHRM2 | DRD5 |  | DRD2 |
| CHRM1 | SLC6A3 |  | DRD4 |
| MAOB | HTR1D |  | CHRM2 |
| BCL2 | SLC6A4 |  | ADRA2B |
|  | HRH4 |  | DRD3 |
|  | GRIN2B |  | ADRA2C |
|  | ADRA1D |  | DRD5 |
|  | CALM1 |  | CHRM4 |
|  | KCNH2 |  | CHRM5 |
|  |  |  | DRD1 |
|  |  |  | SLC6A3 |
|  |  |  | ADRB1 |
|  |  |  | GRIN1 |
|  |  |  | HTR5A |
|  |  |  | HRH3 |
|  |  |  | HTR2B |
|  |  |  | ADRB2 |
|  |  |  | ORM1 |
|  |  |  | CALM1 |
|  |  |  | SMPD1 |
|  |  |  | CALY |
|  |  |  | GSTP1 |
|  |  |  | GRIN2B |
|  |  |  | MCHR1 |
|  |  |  | SLC18A2 |
|  |  |  | GABRA1 |
|  |  |  | CA2 |
|  |  |  | CA3 |
|  |  |  | S100A4 |
|  |  |  | TNNC1 |

| **Asthma-Hypertension** | | | |
| --- | --- | --- | --- |
| Asthma drugs not contraindicated in hypertension | Hypertension drugs not contraindicated in asthma | Asthma drugs contraindicated in hypertension | Hypertension drugs contraindicated in asthma |
| NR3C1 | ADRB1 | NR3C1 | ADRB2 |
| ADRB2 | ADRB2 | PGR | ADRB1 |
| GLT6D1 | CA4 | NR3C2 | SLC12A3 |
| LGALS3 | CA1 | PLA2G4A | KCNMA1 |
| GLTP | CA2 | CYSLTR1 | AGTR1 |
| GLB1 | SLC12A3 | ALOX5 | HMGCR |
| CYSLTR2 | KCNMA1 | ANXA1 | ITGAL |
| CYSLTR1 | CACNA1C | NR0B1 | CA1 |
| FPR1 | CACNA1G | NOS2 | CACNB1 |
| PTGDR | CACNA1B | NR1I2 | CACNA1B |
| HSP90AA1 | ACE | ADRB2 | CACNA1C |
| HDAC2 | CACNA1F | ADRB1 | CACNA2D3 |
| ADORA2A | CACNA1S | ADRB3 | CACNA1I |
| PDE3A | CACNA1D | ADRA1B | SMPD1 |
| ADORA1 | BDKRB1 | ADRA1A | SCNN1G |
| PDE4A | AGTR1 | ADRA1D | SCNN1A |
| IL5 | ADRA2B | ADRA2B | SCNN1B |
| MUC2 | ADRA2A | ADRA2A | SCNN1D |
| RNASE3 | SLC12A1 | TNF | ACE |
| TNF | CA12 | ADORA1 | BDKRB1 |
| NFKB1 | CA9 | PDE3A | MMP9 |
| ADRA1A | ATP1A1 | ADORA3 | MMP2 |
| ADRB1 | CA7 | HDAC2 | LTA4H |
| ADRA2A | CACNG1 | CHRM1 | NR3C2 |
| ADRB3 | NOS2 | CHRM2 | CACNB2 |
| ADRA1B | NOS3 | CHRM3 | CACNA1D |
| ADRA2B | NISCH | GRIN1 | CACNA2D1 |
| ADRA1D | CACNA1H | SLC6A3 | PDE1B |
| ADRA2C | CACNA2D2 | SLC6A4 | CACNA1S |
| ALOX5 | CACNB2 | HRH1 | CACNA2D2 |
|  | CACNA2D1 | SLC6A2 | PDE1A |
|  | HTR1A | CHRM5 | CACNA1H |
|  | ADRB3 | CHRM4 | CALM1 |
|  | HTR1B | ADORA2B | TNNC1 |
|  | ADRA1B | PDE4A | TNNC2 |
|  | KCNH2 | PDE4B | SCN1A |
|  | ADRA1A | ADORA2A | GABRA1 |
|  | ADRA1D | PDE5A | CA4 |
|  | SLC18A1 | CPNE1 | GRIK1 |
|  | SLC18A2 | TTLL3 | CA2 |
|  | BIRC5 | RIC3 | CACNA1E |
|  | SLC12A2 | PARP1 | CA3 |
|  |  | NOMO1 | REN |
|  |  | HM13 | ADRB3 |
|  |  |  | HTR1B |
|  |  |  | HTR1A |
|  |  |  | SLC12A1 |
|  |  |  | GPR35 |
|  |  |  | JUN |
|  |  |  | PPARG |
|  |  |  | AR |
|  |  |  | CYP11B2 |
|  |  |  | NR3C1 |
|  |  |  | PGR |
|  |  |  | SHBG |
|  |  |  | CYP17A1 |
|  |  |  | SRD5A1 |
|  |  |  | CACNG1 |
|  |  |  | NR1I2 |
|  |  |  | AHR |
|  |  |  | DPP4 |
|  |  |  | HDAC2 |
|  |  |  | NR1I3 |
|  |  |  | SFRP4 |
|  |  |  | ADRA2A |
|  |  |  | ADRA1A |
|  |  |  | ADRA2B |
|  |  |  | ADRA1D |
|  |  |  | ADRA1B |
|  |  |  | ADRA2C |
|  |  |  | ASIC1 |
|  |  |  | ASIC2 |
|  |  |  | SLC9A1 |
|  |  |  | AOC1 |
|  |  |  | PLAU |
|  |  |  | KCNH7 |
|  |  |  | KCNH6 |
|  |  |  | KCNH2 |
|  |  |  | TGFB1 |
|  |  |  | KCNJ11 |
|  |  |  | CACNA1G |
|  |  |  | CACNA1A |
|  |  |  | SLC6A4 |
|  |  |  | KCND3 |
|  |  |  | APLP1 |
|  |  |  | APP |
|  |  |  | CYB5A |
|  |  |  | HDAC1 |
|  |  |  | ASPA |
|  |  |  | ESR1 |
|  |  |  | HDAC4 |
|  |  |  | APLP2 |
|  |  |  | DAND5 |
|  |  |  | HDAC8 |
|  |  |  | TP73 |
|  |  |  | DDC |
|  |  |  | DCD |
|  |  |  | MPG |
|  |  |  | HBB |
|  |  |  | APOA2 |
|  |  |  | INS |
|  |  |  | AGT |
|  |  |  | CPN1 |
|  |  |  | C4BPB |
|  |  |  | ORM2 |
|  |  |  | IL3 |
|  |  |  | C1R |
|  |  |  | CFI |
|  |  |  | MT2A |
|  |  |  | CP |
|  |  |  | IGFALS |
|  |  |  | KNG1 |
|  |  |  | IDH3G |
|  |  |  | IGLV3-21 |
|  |  |  | MDM2 |
|  |  |  | CFB |
|  |  |  | KRT9 |
|  |  |  | ITIH1 |
|  |  |  | S100A9 |
|  |  |  | PARP1 |
|  |  |  | MT3 |
|  |  |  | FCN3 |
|  |  |  | C3 |
|  |  |  | PZP |
|  |  |  | NME1 |
|  |  |  | ENO1 |
|  |  |  | KRT6A |
|  |  |  | PGLYRP2 |
|  |  |  | PDCD6 |
|  |  |  | A1BG |
|  |  |  | PRDX1 |
|  |  |  | TUFM |
|  |  |  | VTN |
|  |  |  | ITIH4 |
|  |  |  | F13B |
|  |  |  | S100A2 |
|  |  |  | C1S |
|  |  |  | C4BPA |
|  |  |  | SERPINA4 |
|  |  |  | HPR |
|  |  |  | ITIH3 |
|  |  |  | C1QB |
|  |  |  | KRT5 |
|  |  |  | C8G |
|  |  |  | C8A |
|  |  |  | TTR |
|  |  |  | FN1 |
|  |  |  | GSN |
|  |  |  | IGHA1 |
|  |  |  | GAPDHS |
|  |  |  | SEMG1 |
|  |  |  | S100A8 |
|  |  |  | PON1 |
|  |  |  | APOA1 |
|  |  |  | TP53 |
|  |  |  | IGHM |
|  |  |  | CCS |
|  |  |  | HBA1 |
|  |  |  | DSP |
|  |  |  | SERPINA6 |
|  |  |  | GLRA1 |
|  |  |  | S100A7 |
|  |  |  | SERPINA1 |
|  |  |  | EEF1A1 |
|  |  |  | ITIH2 |
|  |  |  | IGKV1-17 |
|  |  |  | SOD1 |
|  |  |  | APOBR |
|  |  |  | C1QC |
|  |  |  | SERPIND1 |
|  |  |  | SIVA1 |
|  |  |  | F12 |
|  |  |  | APOA4 |
|  |  |  | P4HB |
|  |  |  | KRT14 |
|  |  |  | F2 |
|  |  |  | APCS |
|  |  |  | ARG1 |
|  |  |  | CFH |
|  |  |  | IDH3A |
|  |  |  | HRNR |
|  |  |  | TAB1 |
|  |  |  | JUP |
|  |  |  | CLU |
|  |  |  | FGA |
|  |  |  | APOE |
|  |  |  | PSPH |
|  |  |  | SELENOP |
|  |  |  | SERPINA3 |
|  |  |  | ALDOA |
|  |  |  | MT1A |
|  |  |  | KRT16 |
|  |  |  | APOL1 |
|  |  |  | MGMT |
|  |  |  | PDIA3 |
|  |  |  | KRT1 |
|  |  |  | BRCC3 |
|  |  |  | AHSG |
|  |  |  | KRT10 |
|  |  |  | TPI1 |
|  |  |  | A2M |
|  |  |  | C8B |
|  |  |  | JCHAIN |
|  |  |  | C4B |
|  |  |  | C5 |
|  |  |  | TF |
|  |  |  | IGKV3-20 |
|  |  |  | UTRN |
|  |  |  | CPN2 |
|  |  |  | KLKB1 |
|  |  |  | KRT2 |
|  |  |  | HIF1A |
|  |  |  | NDUFC2 |
|  |  |  | VCAM1 |
|  |  |  | SELE |
|  |  |  | KCNJ4 |
|  |  |  | KCNJ2 |
|  |  |  | NPPB |
|  |  |  | GJA1 |
|  |  |  | VEGFA |
|  |  |  | CHRM1 |
|  |  |  | CHRM2 |
|  |  |  | CHRM4 |
|  |  |  | CHRM3 |
|  |  |  | CHRM5 |

| **Type 2 diabetes-Obesity** | | | |
| --- | --- | --- | --- |
| Type 2 diabetes drugs not contraindicated in BMI >30 kg/m2 | BMI >30 kg/m2 drugs not contraindicated in type 2 diabetes | Type 2 diabetes drugs contraindicated in BMI >30 kg/m2 | BMI >30 kg/m2 drugs contraindicated in type 2 diabetes |
| MGAM | SLC18A2 | ABCC8 | SLC6A2 |
| SI | ADRA2A | KCNJ1 | CHRNA3 |
| AMY2A | ADRA1A | KCNJ11 | SLC6A3 |
| GAA | SLC6A3 | GLP1R | HTR3A |
| GLP1R | HTR2C | PPARG | SLC6A4 |
| DPP4 | OPRD1 | ACSL4 | CHRNA2 |
| DRD3 | OPRM1 | RXRB | KCNH2 |
| DRD5 | OPRK1 | PPARA | HTR2C |
| ADRA1D | SIGMAR1 | RXRG | CHRNB4 |
| ADRA2B | DRD1 | RXRA | CKS1B |
| DRD1 |  | PPARD | MAOB |
| HTR2C |  | PRKAB1 | MAOA |
| HTR1D |  | ETFDH | NPY |
| HTR2A |  | GPD1 | PNLIP |
| HTR7 |  | HMGCR | LIPF |
| ADRA1A |  | ITGAL | FASN |
| HTR2B |  | HDAC2 | CARTPT |
| ADRA2C |  | AHR | ADRB1 |
| DRD4 |  | DPP4 | SLC18A2 |
| ADRA1B |  | NR1I3 | TAAR1 |
| HTR1B |  | ABCC9 | DRD2 |
| HTR1A |  | ABCB11 | ADRA1A |
| ADRA2A |  | ABCA1 |  |
| DRD2 |  | CFTR |  |
| SLC5A2 |  | TRPM4 |  |
| ABCC8 |  | CPT1A |  |
| VEGFA |  | MAOB |  |
| KCNJ8 |  | SCN1A |  |
| SLC5A6 |  | GABRA1 |  |
| LIPT1 |  | CA4 |  |
| LIAS |  | GRIK1 |  |
| GANC |  | CA2 |  |
| GANAB |  | CA1 |  |
| PPARG |  | CACNA1C |  |
| KCNJ1 |  | CACNA1E |  |
|  |  | CA3 |  |
|  |  | GCGR |  |
|  |  | GLP2R |  |
|  |  | RAMP1 |  |
|  |  | CALCR |  |
|  |  | RAMP3 |  |
|  |  | RAMP2 |  |

| **Rheumatoid arthritis-Osteoporosis** | | | |
| --- | --- | --- | --- |
| Rheumatoid arthritis drugs not contraindicated in osteoporosis | Osteoporosis drugs not contraindicated in rheumatoid arthritis | Rheumatoid arthritis drugs contraindicated in osteoporosis | Osteoporosis drugs contraindicated in rheumatoid arthritis |
| PTGS1 | ESRRB | BCL2 | SCN5A |
| PTGS2 | ESR2 | PTGS2 | SCN9A |
| PRDX5 | ESRRA | THBD | SCN10A |
| IKBKB | SHBG | CFTR | EGFR |
| ANXA1 | PTK2B | PPARG | SCN4A |
| NR3C1 | AKT1 | PTGS1 | ORM1 |
| SCN10A | NCOA2 | FABP2 | ORM2 |
| PTGER1 | NCOA1 | GP1BA | ESR1 |
| TLR9 | CYP1B1 | PPARA | ESR2 |
| HMGB1 | GPER1 | S100A7 | MT-ATP6 |
| ACE2 | NR1I2 | NR3C1 | GPER1 |
| GSTM1 | ESR1 | ANXA1 | NR1I2 |
| TNF | TOP2A | NR0B1 | BECN1 |
| GSTA2 | CTPS1 | NOS2 | CHRNA4 |
| MC2R | PPAT | NR1I2 | TNNC2 |
| CRH | GLUL | TRPV1 | CACNA1C |
| PPARA | PGR | PTGES3 | SPTBN1 |
| PPARG | AR | TNF | ATP2C1 |
| CXCR1 | NR3C1 | ALOX5 | TNNC1 |
| ALOX5 | SOST | PLA2G1B | CP |
| KCNQ2 |  | TBXAS1 | CALM1 |
| KCNQ3 |  | SLC7A11 | ALPP |
| PPARD |  | IKBKB | S100A8 |
| AKR1B1 |  | ACAT1 | S100A9 |
| PTGDR2 |  | CHUK | S100B |
| MAPK3 |  | HRH2 | S100A13 |
| AKR1B10 |  | ACHE | BMP4 |
| GABRA1 |  | CA2 | PDCD6 |
| GLRA1 |  | CA3 | MGP |
| TNFSF11 |  | FCGR1A | COMP |
|  |  | FCGR3A | CAST |
|  |  | FCGR2B | S100A2 |
|  |  | C1QA | PCDH19 |
|  |  | FCGR2C | AOC1 |
|  |  | FCGR2A | PTH1R |
|  |  | LTA | CALR3 |
|  |  | FCGR3B | CALM2 |
|  |  | ATP4A | CALR |
|  |  | AHR | RGN |
|  |  | FKBP1A | CHP1 |
|  |  | TNFSF11 | NUCB2 |
|  |  | PDPK1 | PEF1 |
|  |  | CDH11 | CASQ1 |
|  |  | DHODH | SRI |
|  |  | PTK2B | GCA |
|  |  | TLR7 | CANX |
|  |  | TLR9 | CALM3 |
|  |  | ACE2 | S100A6 |
|  |  | RXRA | CASR |
|  |  | RAC1 | NRXN1 |
|  |  |  | S100A16 |
|  |  |  | CASQ2 |
|  |  |  | SPARC |
|  |  |  | NUCB1 |
|  |  |  | CIB2 |
|  |  |  | CIB1 |
|  |  |  | CADPS |
|  |  |  | CAPS |
|  |  |  | FBN3 |
|  |  |  | CADPS2 |
|  |  |  | TPT1 |
|  |  |  | NCS1 |
|  |  |  | CALB2 |
|  |  |  | FBN2 |
|  |  |  | PCAP |
|  |  |  | FDPS |
|  |  |  | GGPS1 |
|  |  |  | VDR |
|  |  |  | SERPINB9 |
|  |  |  | TFF1 |

| **Chronic obstructive pulmonary disease-Heart failure** | | | |
| --- | --- | --- | --- |
| Chronic obstructive pulmonary disease drugs not contraindicated in heart failure | Heart failure drugs not contraindicated in chronic obstructive pulmonary disease | Chronic obstructive pulmonary disease drugs contraindicated in heart failure | Heart failure drugs contraindicated in chronic obstructive pulmonary disease |
| CHRM3 | BDKRB1 | ADRB2 | ADRB1 |
| CHRM5 | ACE | NR3C1 | ADRB2 |
| CHRM2 |  | ANXA1 | ACE |
| CHRM1 |  | ADRB1 | SLC12A1 |
| CHRM4 |  | ADRB3 | SLC12A2 |
| NGF |  | CHRM1 | SLC12A4 |
| ADRB2 |  | CHRM2 | SLC12A5 |
| ADRB1 |  | CHRM3 | CFTR |
| ADRB3 |  | PDE4A | MMP9 |
| TNF |  | PDE4D | MMP2 |
| KCNK3 |  | PDE4B | LTA4H |
| KCNK9 |  | PDE4C | BDKRB1 |
| ADORA2A |  | PDE3A | GPR35 |
| PDE2A |  | ADORA2B | CA2 |
| GLTP |  | HDAC2 | AOC3 |
| GLB1 |  | ADORA2A | P4HA1 |
| LGALS3 |  | ADORA1 | HIF1A |
| GLT6D1 |  | PDE5A | AGTR1 |
| HDAC2 |  | CPNE1 | AR |
| PDE3A |  | TTLL3 | NR3C2 |
| ADORA1 |  | RIC3 | CYP11B2 |
| PDE4A |  | PARP1 | NR3C1 |
|  |  | NOMO1 | PGR |
|  |  | HM13 | SHBG |
|  |  | CHRM5 | CYP17A1 |
|  |  | CHRM4 | SRD5A1 |
|  |  |  | CACNG1 |
|  |  |  | NR1I2 |
|  |  |  | PPARG |

| **Schizophrenia-Asthma** | | | |
| --- | --- | --- | --- |
| Schizophrenia not contraindicated in asthma | Schizophrenia contraindicated in asthma | Asthma drugs not contraindicated in schizophrenia | Asthma drugs contraindicated in schizophrenia |
| DRD2 | ERBB4 | IL10 | APOE |
| DRD1 | CCKBR | ADRB1 | SI |
| ADRA1A | DRD2 | CYP2D6 | PYGL |
| CHRM1 | MPHOSPH8 | CYP3A5 | CCL17 |
| DRD3 | ANKK1 | NR3C1 | CDK2 |
| EHMT2 | CYP3A5 | NFE2L2 | NFE2L2 |
| XBP1 | SLC1A1 | ADORA2B | NR5A1 |
| ATAD5 | CAMK2B | CYP3A4 | FGF2 |
| HTR2B | MTOR | PDE4B | CYP3A4 |
| ADRA1B | MCL1 | GLCCI1 | ACTC1 |
| PPARD | MTHFR | UGT1A1 | TDO2 |
| HRH1 | CYP3A4 | ADRB2 | AGTR1 |
| HTR1A | DRD1 | POLK | TYMS |
| ADRA1D | CYP2D6 | CHRM1 | DOK5 |
| HRH2 | GIPR | DRD1 | MAPK1 |
| ADRA2B | BDNF | CYP2C19 | ANXA1 |
| AR | HTR1E | HYAL2 | GSTP1 |
| HTR2A | GRIA1 | APEX1 | HIF1A |
| DRD4 | HTR6 | ANXA1 | CYSLTR1 |
| IDH1 | UGT1A5 | IL1B | PZP |
| CYP2D6 | DRD3 | PDE4A | HAS3 |
| KCNH2 | CYP2C19 | ALOX5 | PTPRC |
| HTR7 | GRIN2B | HRH1 | ABCB1 |
| HTR6 | ABCB1 | ADRA1B | CHRM3 |
| NRG3 | CYP1A1 | T | NRG1 |
| CHRM4 | RYR1 | PPARD | BAZ2B |
| NPSR1 | HTR1A | TSHB | IGFBP1 |
| CHRM3 | CYP1A2 | SERPINA6 | IFNG |
| CBX1 | TP53 | PIK3CG | TGFBR3 |
| GMNN | ADRA2C | ADRA2B | ADRB2 |
| CYP2C19 | UGT1A6 | ADRA1D | GGT1 |
| CHRM2 | ADRA2B | MPO | ERBB2 |
| CYP3A4 |  | KIAA0391 | SLC2A1 |
| HTR2C | DRD4 | CYP2C9 | TGFB1 |
| HTR1B | GRM7 | CYP2E1 | CTNNB1 |
| MPHOSPH8 | TJP1 | BGLAP | SFTPA1 |
| THPO | ADRA1A | AR | CRISPLD2 |
| HIF1A | MAPK1 | TNFRSF11B | NPPC |
| ADRA2C | USP1 | IL2RA | USP1 |
| CYP1A2 | TNF | CRHR1 | TG |
| CHRM5 | CHRM5 | PDE3A | CYP2C19 |
| KDM4A | MC4R | CYSLTR1 | CDK4 |
| HTR5A | RGS4 | CA10 | NR3C1 |
| ADRA2A | MSANTD1 | TPM3 | CYP3A5 |
| CERKL | EPM2A |  | MT1F |
| NUBPL | HTR2A | IDH1 | UBR1 |
| DRD5 | AKT1 | TNFRSF8 | CYP2D6 |
| SMAD3 | PMCH | FEN1 | BIRC5 |
| HTT | HTR7 | SLC2A4 | ADA |
| CYP2C9 | CYP2J2 | CHRM2 | LTA4H |
| CELF4 | CHRM2 | HSPB1 | TSC22D3 |
|  | SH2B1 | CSF2 | RARA |
|  | HTR1D | MAPT | PTH1R |
|  | ADRA1B | MMP12 | CYP2C9 |
|  | HTR2C | MAPKAPK2 | IRS2 |
|  | HRH1 | COL22A1 | GSTM1 |
|  | GRIK4 | ADRA2C | TJP1 |
|  | ATXN2 | CLOCK | NR1I2 |
|  | JUND | PLA2G4A | AR |
|  | HTR3A | MTOR | CYP1A2 |
|  | CBX1 | MMP2 | VDR |
|  | HRH4 | HSD17B10 | CDH17 |
|  | GHRL | ADORA1 | APOB |
|  | SLC6A4 | CYP1A2 | TPMT |
|  | GDNF | NR1I2 | S100A10 |
|  | CNR1 | ADRA2A | G6PD |
|  | ATP1A2 | NOS1 | CHRM2 |
|  | GMNN | ADRA1A | HSD17B10 |
|  | HLA-DRB5 | XDH | LTC4S |
|  | HTR2B | EHMT2 | CA10 |
|  | HLA-B | CHRM3 | ARNT |
|  | SPOPL | VDR | UGT1A1 |
|  | SRC | CCL5 | ALOX5 |
|  | IDH1 | RECQL | JUNB |
|  | CHRM3 | EDN1 | SERPINA6 |
|  | TAAR6 | HIF1A | HCG22 |
|  | GAA | G6PD | SERPINE1 |
|  | GRIN2A | DUSP1 | SOAT1 |
|  | DRD5 | PDE5A | ABCC1 |
|  | IL2 | GUCA2A | HYAL2 |
|  | CACNA1C | PRKAR2B | THRSP |
|  | FMO1 | CCL11 | BDNF |
|  | FKBP5 | PIK3CB | NFKB1 |
|  | GLP1R | CYP3A7 | MC2R |
|  | DLG4 | PTH | ABCC9 |
|  | CALY | ADORA2A | IL2RA |
|  | NPY | IFNG | GATA3 |
|  | EIF2AK4 | NOS2 | LINC00251 |
|  | SLC6A3 | CD44 | HSPA8 |
|  | CYP2C9 | MAPK10 | ITGAM |
|  | LIPA | ITGAM | LOH19CR1 |
|  | FASN | KDM4E | TYMSOS |
|  | ADRB2 | HSPA4 | HRH1 |
|  | FAAH | CRHR2 | KRT19 |
|  | LDLR | GHRL | BCHE |
|  | RABEP1 | FCER1A | NTRK1 |
|  | TNFRSF11A | CAST | ITGB2 |
|  | LEP | CD163 | CXCL10 |
|  | EHMT2 |  | SMAD2 |
|  | HSD17B10 |  | GMNN |
|  | ITIH3 |  | HTR7 |
|  | UGT1A9 |  | RELA |
|  | KDM4A |  | CTLA4 |
|  | ADRA2A |  | CDK6 |
|  | IL1RN |  | VCAM1 |
|  | AR |  | CD1A |
|  | NR4A1 |  | DPEP1 |
|  | NR1I2 |  |  |
|  | IL1A |  | BMP7 |
|  | TAC1 |  | TEAD1 |
|  | HSPA4 |  | FABP1 |
|  | MAPK14 |  | GLCCI1 |
|  | GRIN1 |  | CXCL12 |
|  | THPO |  | KAT2A |
|  | KCNH2 |  | IL5 |
|  | PIK3CG |  | CHRM1 |
|  | HRH3 |  | SDS |
|  | SLC6A2 |  | SLCO2B1 |
|  | AKAP13 |  | F9 |
|  | CYP1B1 |  | MYC |
|  | COMT |  | CALCA |
|  | ARID5B |  | TP53 |
|  | GSTM3 |  | GLS |
|  | TPMT |  | CRHR1 |
|  | HTR1B |  | MBP |
|  | BAZ2B |  | CD86 |
|  | ADRB3 |  | CARTPT |
|  | CYP2E1 |  | RPS19 |
|  | DTNBP1 |  | CYP2C8 |
|  | GCG |  | NOTCH1 |
|  | NTS |  | CHAT |
|  | SIGMAR1 |  | RB1 |
|  | CHAT |  | UGT1A3 |
|  | PLA2G1B |  | CHKA |
|  | HLA-C |  | MMP1 |
|  | HTR3E |  | MLLT3 |
|  | CYP3A43 |  | LIF |
|  | PPARD |  | SAG |
|  | CHRM4 |  | SMAD3 |
|  | ZIC4 |  | RET |
|  | PPA2 |  | BCL2 |
|  | PLEKHA6 |  | VWF |
|  | RAPGEF4 |  | CD80 |
|  | KCNJ3 |  | GSTA1 |
|  | HSPG2 |  | GALNT17 |
|  | ANPEP |  | FOLR1 |
|  | PPARG |  | LPL |
|  | ADRB1 |  | SLC2A4 |
|  | APOB |  | DROSHA |
|  | ABCB5 |  | POLB |
|  | UGT1A1 |  |  |
|  | SV2C |  |  |
|  | CAMK2G |  |  |
|  | IFNG |  |  |
|  | FMO3 |  |  |
|  | UGT1A8 |  |  |
|  | CCK |  |  |
|  | CHRM1 |  |  |
|  | PRL |  |  |
|  | MAOB |  |  |
|  | UGT1A3 |  |  |
|  | UGT1A4 |  |  |
|  | NKX6-1 |  |  |
|  | HTT |  |  |
|  | AHR |  |  |
|  | NTRK2 |  |  |
|  | DPP6 |  |  |
|  | CCL2 |  |  |
|  | NT5E |  |  |
|  | NTF3 |  |  |
|  | GNB3 |  |  |
|  | GSTM1 |  |  |
|  | UBE2N |  |  |
|  | ADA |  |  |
|  | ABCG2 |  |  |
|  | TOR1A |  |  |
|  | GRID2 |  |  |
|  | RTKN2 |  |  |
|  | PDE4D |  |  |
|  | HOMER1 |  |  |
|  | HLA-DPB1 |  |  |
|  | CHRNA7 |  |  |
|  | RGS2 |  |  |
|  | GSTT1 |  |  |
|  | TBC1D1 |  |  |
|  | UGT1A7 |  |  |
|  | POLI |  |  |
|  | GFER |  |  |
|  | SLC6A5 |  |  |
|  | GRM3 |  |  |
|  | HLA-DRB3 |  |  |
|  | MIR582 |  |  |
|  | CSF2 |  |  |
|  | UGT1A10 |  |  |
|  | RAD52 |  |  |
|  | C3 |  |  |
|  | SLC2A4 |  |  |
|  | KAT2A |  |  |

| **Schizophrenia-Rheumatoid arthritis** | | | |
| --- | --- | --- | --- |
| Schizophrenia not contraindicated in rheumatoid arthritis | Schizophrenia contraindicated in rheumatoid arthritis | Rheumatoid arthritis drugs not contraindicated in schizophrenia | Rheumatoid arthritis drugs contraindicated in schizophrenia |
| DRD2 | CCKBR | FMO3 | APOE |
| DRD1 | DRD2 | BAX | CYP2D6 |
| ADRA1A | MPHOSPH8 | TNF | CASP3 |
| ERBB4 | ANKK1 | PTGS2 | SI |
| CHRM1 | CYP3A5 | LOH19CR1 | PTGS2 |
| CAMK2B | SLC1A1 | TSHB | ALOX12 |
| MTOR | MCL1 | MTHFR | PYGL |
| DRD3 | MTHFR | KCNA1 | CCL17 |
| EHMT2 | CYP3A4 | DHODH | NOS1 |
| XBP1 | DRD1 | FCGR3B | CDK2 |
| ATAD5 | CYP2D6 | CYP1A2 | NR1I2 |
| CYP3A4 | GIPR | IL10 | NFE2L2 |
| HTR2B | BDNF | PPAT | CYP3A4 |
| ADRA1B | HTR1E | HSD17B10 | CACNA2D1 |
| PPARD | GRIA1 | CD83 | STAT3 |
| TP53 | HTR6 | AR | ATP2A1 |
| HRH1 | UGT1A5 | ANXA1 | HRH2 |
| HTR1A | DRD3 | HMGB1 | NR5A1 |
| ADRA1D | CYP2C19 | TYMSOS | CYP2C9 |
| MAPK1 | GRIN2B | AKR1B1 | FGF2 |
| USP1 | ABCB1 | IL18 | ACTC1 |
| RGS4 | CYP1A1 | CYP3A4 | TDO2 |
| HRH2 | RYR1 | PTGS1 | AGTR1 |
| ADRA2B | HTR1A | ACAT1 | TYMS |
| AR | CYP1A2 | G6PD | CA12 |
| HTR2A | ADRA2C | AOX1 | FAAH |
| DRD4 | UGT1A6 | CAPN10 | ATP4A |
| IDH1 | ADRA2B | ALOX5 | PTGS1 |
| CYP1A2 | DRD4 | GLRB | DOK5 |
| ATXN2 | GRM7 | HIF1A | PTGER4 |
| CYP2D6 | TJP1 | CYP2C9 | DDIT3 |
| KCNH2 | ADRA1A | APC | ANXA1 |
| HTR7 | TNF | TNFRSF10A | GSTP1 |
| HTR6 | CHRM5 | ABCB1 | MMP9 |
| NRG3 | MC4R | BAZ2B | CACNA1D |
| CHRM4 | MSANTD1 | NFE2L2 | PSORS1C1 |
| NPSR1 | EPM2A | TLR9 | HIF1A |
| EPM2A | HTR2A | TNFSF11 | PZP |
| MC4R | AKT1 | KDM4A | ATP2A3 |
| CYP2C19 | PMCH | MAPT | AR |
| AKT1 | HTR7 | TCF7L2 | ATP5E |
| CHRM3 | CYP2J2 | CYP2D6 | HAS3 |
| CBX1 | CHRM2 | TARDBP | PTPRC |
| GMNN | SH2B1 | BGLAP | APC |
| GLP1R | HTR1D | PLA2G6 | ABCB1 |
| CALY | ADRA1B | HTT | NRG1 |
| CHRM2 | HTR2C | BRCA1 | IGFBP1 |
| LIPA | HRH1 | TYMS | IFNG |
| HTR2C | GRIK4 | HBB | CYP2J2 |
| HTR1B | JUND | POLI | CXCL8 |
| MPHOSPH8 | HTR3A | HPGD | TRPV1 |
| THPO | CBX1 | FKBP1A | TGFBR3 |
| HIF1A | HRH4 | FTO | CACNA1G |
| ADRA2C | GHRL | POLK | CBX1 |
| CHRM5 | SLC6A4 | LIF | GGT1 |
| KDM4A | GDNF | EHMT2 | ERBB2 |
| APOB | CNR1 | ABCG2 | CYP1A2 |
| HTR5A | RGS4 | PPP3CA | SMN2 |
| ADRA2A | ATP1A2 | SCN10A | AHR |
| CERKL | GMNN | CYP3A5 | CACNB1 |
| NUBPL | TP53 | ABCC4 | SLC2A1 |
| UBE2N | HLA-DRB5 | TG | PLA2G2A |
| DRD5 | HTR2B | ACKR1 | TGFB1 |
| SMAD3 | HLA-B | DHFR | CTNNB1 |
| RABEP1 | SPOPL | PLA2G1B | SULT1A1 |
| HTT | SRC | RXRA | SFTPA1 |
| CSF2 | IDH1 | UGDH | CRISPLD2 |
| RAD52 | CHRM3 | ASS1 | NR3C1 |
| CYP2C9 | TAAR6 | TAC1 | SULT1A3 |
| SH2B1 | GAA | PLG | BAZ2B |
| CELF4 | GRIN2A | GABRG2 | NPPC |
|  | DRD5 | IL1A | CCND1 |
|  | IL2 | TSPYL1 | CYP7B1 |
|  | CACNA1C | TP53 | TG |
|  | FMO1 | TLR7 | CYP2C19 |
|  | FKBP5 | PIK3CA | CDK4 |
|  | DLG4 | SPP1 | CDKN1A |
|  | NPY | SST | MT1F |
|  | EIF2AK4 | RB1 | IL23R |
|  | SLC6A3 | APP | NOS2 |
|  | CYP2C9 | NTRK2 | HEXB |
|  | FASN | CLCNKA | UBR1 |
|  | ADRB2 | NR1I2 | G6PD |
|  | FAAH | TAT | BIRC5 |
|  | LDLR | NUDT15 | CYP2E1 |
|  | RABEP1 | ATAD5 | ADA |
|  | TNFRSF11A | RAB9A | PLA2G1B |
|  | LEP | CD34 | TSC22D3 |
|  | EHMT2 | NR3C1 | RARA |
|  | HSD17B10 | PPARA | PTH1R |
|  | ITIH3 | PMP22 | CYP2A6 |
|  | UGT1A9 | TPMT | CSF2 |
|  | KDM4A | NOTCH1 | IRS2 |
|  | ADRA2A | PNPLA3 | MPO |
|  | ATXN2 | PPARD | KDM4A |
|  | IL1RN | MMP1 | GSTM1 |
|  | AR | CRTC2 | PRL |
|  | NR4A1 | CYP2C8 | TJP1 |
|  | NR1I2 | GABRB2 | KDM4E |
|  | IL1A | NT5E | CA9 |
|  | TAC1 | BIRC5 | HSPA8 |
|  | HSPA4 | CXCL8 | CDH17 |
|  | MAPK14 | UGT1A8 | CACNA2D3 |
|  | GRIN1 | IKBKB | BCAR1 |
|  | THPO | IL3 | CYP3A5 |
|  | PIK3CG | CYP2C19 | APOB |
|  | HRH3 | CYP3A7 | TPMT |
|  | SLC6A2 | HLA-DQA1 | PTH |
|  | AKAP13 | TYMP | PIK3CA |
|  | CYP1B1 | ITPA | TNFRSF11A |
|  | COMT | UGT1A4 | RORC |
|  | ARID5B | CCL17 | S100A10 |
|  | GSTM3 | UGT2B7 | UGT1A9 |
|  | MTOR | MC2R | HPGD |
|  | TPMT | SUMO4 | PTGER1 |
|  | HTR1B | TLR3 | PDK1 |
|  | BAZ2B | ESR1 | KCNQ4 |
|  | ADRB3 | PTH | PLAT |
|  | CYP2E1 | IDH1 | HSD17B10 |
|  | DTNBP1 | GPR35 | TARDBP |
|  | GCG | GLRA1 | ARNT |
|  | NTS | DDRGK1 | CACNA2D4 |
|  | SIGMAR1 | SMN1 | UGT1A1 |
|  | CHAT | ABCC2 | JUNB |
|  | PLA2G1B | HLA-B | UGT2B4 |
|  | HLA-C | BDNF | SERPINA6 |
|  | HTR3E | GSTM1 | HCG22 |
|  | CYP3A43 | NOD2 | ATP2A2 |
|  | PPARD | PPARG | PLCG1 |
|  | CHRM4 | ALPP | SERPINE1 |
|  | ZIC4 | NRG1 | HTR4 |
|  | PPA2 | HSPA8 | SOAT1 |
|  | PLEKHA6 | LCAT | PLK1 |
|  | RAPGEF4 | MYOD1 | HYAL2 |
|  | KCNJ3 | HSD11B1 | THRSP |
|  | HSPG2 | CYP2J2 | BDNF |
|  | GLP1R | NFKB2 | NFKB1 |
|  | ANPEP | FOXP3 | SLCO1C1 |
|  | PPARG | KDM4E | CXCR1 |
|  | ADRB1 | CTLA4 | MC2R |
|  | ABCB5 | NAT1 | CACNA1B |
|  | UGT1A1 | SMN2 | BRD2 |
|  | SV2C | GPT | CACNB2 |
|  | CAMK2G | MPO | OPRM1 |
|  | IFNG | ZSCAN25 | PPARG |
|  | KCNH2 | FCGR3A | IL2RA |
|  | FMO3 | FGF6 | VIP |
|  | UGT1A8 | GABRA1 | EDN1 |
|  | CCK | POR | GATA3 |
|  | CHRM1 | MYC | LINC00251 |
|  | PRL | BCHE | IGFBP3 |
|  | MAOB | HLA-DRB1 | HSPB1 |
|  | UGT1A3 | KCNQ1 | PLA2G4A |
|  | UGT1A4 | TLR4 | UGT1A6 |
|  | NKX6-1 | KCNJ11 | HLA-DRB1 |
|  | HTT | DSE | ITGAM |
|  | AHR |  | CASP9 |
|  | NTRK2 |  | UGT2B17 |
|  | DPP6 |  | HSPA4 |
|  | CCL2 |  | LOH19CR1 |
|  | NT5E |  | UGT1A3 |
|  | NTF3 |  | TYMSOS |
|  | GNB3 |  | TNF |
|  | GSTM1 |  | PTGES |
|  | ADA |  | VHL |
|  | ABCG2 |  | KRT19 |
|  | TOR1A |  | BCHE |
|  | GRID2 |  | NTRK1 |
|  | RTKN2 |  | F8 |
|  | PDE4D |  | VDR |
|  | HOMER1 |  | AMACR |
|  | HLA-DPB1 |  | KCNQ2 |
|  | CHRNA7 |  | CACNA1F |
|  | RGS2 |  | NOS3 |
|  | GSTT1 |  | IL12B |
|  | TBC1D1 |  | CYP2C8 |
|  | UGT1A7 |  | HLA-DQB1 |
|  | POLI |  | ITGB2 |
|  | GFER |  | GHRL |
|  | SLC6A5 |  | FCGR2A |
|  | GRM3 |  | UGT1A4 |
|  | HLA-DRB3 |  | CXCL10 |
|  | MIR582 |  | ANPEP |
|  | UGT1A10 |  | CACNA1C |
|  | C3 |  | ACHE |
|  | SLC2A4 |  | TNFRSF1B |
|  | KAT2A |  | SMAD2 |
|  |  |  | GDF15 |
|  |  |  | CACNA1I |
|  |  |  | HDC |
|  |  |  | GRP |
|  |  |  | CACNB3 |
|  |  |  | HTR7 |
|  |  |  | HMOX2 |
|  |  |  | RELA |
|  |  |  | MCL1 |
|  |  |  | CTLA4 |
|  |  |  | MAPT |
|  |  |  | ALB |
|  |  |  | CDK6 |
|  |  |  | VCAM1 |
|  |  |  | CD1A |
|  |  |  | SULT2A1 |
|  |  |  | TRAF1 |
|  |  |  | KCNQ1 |
|  |  |  | DPEP1 |
|  |  |  | BMP7 |
|  |  |  | TEAD1 |
|  |  |  | FABP1 |
|  |  |  | GLCCI1 |
|  |  |  | CXCL12 |
|  |  |  | CALCA |
|  |  |  | TAC1 |
|  |  |  | STAT4 |
|  |  |  | KLRD1 |
|  |  |  | TNFRSF1A |
|  |  |  | SULT1A4 |
|  |  |  | IL5 |
|  |  |  | CDKN1B |
|  |  |  | SDS |
|  |  |  | IGF2 |
|  |  |  | UGT2B7 |
|  |  |  | F9 |
|  |  |  | VEGFA |
|  |  |  | COL18A1 |
|  |  |  | MYC |
|  |  |  | MT-CO2 |
|  |  |  | SAA1 |
|  |  |  | TP53 |
|  |  |  | CACNA1S |
|  |  |  | GLS |
|  |  |  | CRHR1 |
|  |  |  | UGT1A |
|  |  |  | CYP19A1 |
|  |  |  | MBP |
|  |  |  | FCGR3A |
|  |  |  | CD86 |
|  |  |  | CARTPT |
|  |  |  | RPS19 |
|  |  |  | UGT2B15 |
|  |  |  | NPIPB8 |
|  |  |  | CACNA2D2 |
|  |  |  | CXCR2 |
|  |  |  | KCNQ5 |
|  |  |  | BGLAP |
|  |  |  | NOTCH1 |
|  |  |  | CHAT |
|  |  |  | EHMT2 |
|  |  |  | RB1 |
|  |  |  | PDPK1 |
|  |  |  | KCNQ3 |
|  |  |  | ABL1 |
|  |  |  | SULT1E1 |
|  |  |  | CHKA |
|  |  |  | CACNA1E |
|  |  |  | MMP1 |
|  |  |  | PLAU |
|  |  |  | LIF |
|  |  |  | SAG |
|  |  |  | UGT1A10 |
|  |  |  | SMAD3 |
|  |  |  | CACNA1H |
|  |  |  | RET |
|  |  |  | IL6 |
|  |  |  | CD84 |
|  |  |  | CACNB4 |
|  |  |  | HLA-E |
|  |  |  | BCL2 |
|  |  |  | VWF |
|  |  |  | CD80 |
|  |  |  | RAPGEF4 |
|  |  |  | GSTA1 |
|  |  |  | FOLR1 |
|  |  |  | LPL |
|  |  |  | SLC2A4 |
|  |  |  | CACNA1A |
|  |  |  | IL1B |
|  |  |  | DROSHA |
|  |  |  | SMN1 |

| **Multiple sclerosis-Peroxisomal disorders** | |
| --- | --- |
| Multiple sclerosis drugs | Peroxisomal disorder drugs |
| BAX | PLA2G1B |
| TSHB | DHCR7 |
| CYP2C9 | AMACR |
| CYP2D6 | ABCB11 |
| ADORA2A-AS1 | NR1H4 |
| KCNA1 | CYP7A1 |
| PPAT | HSD3B7 |
| ATIC | CYP27A1 |
| DHFR | CES1 |
| TYMS | FECH |
| TYMSOS | AKR1D1 |
| CCND1 |  |
| AOX1 |  |
| RYR2 |  |
| POLH |  |
| MAPT |  |
| CALCA |  |
| VDR |  |
| CYP3A4 |  |
| PYGL |  |
| NFKB1 |  |
| PTPRM |  |
| FOS |  |
| CYP3A5 |  |
| TPM3 |  |
| GABBR1 |  |
| PACSIN2 |  |
| AFP |  |
| KLRD1 |  |
| NFE2L2 |  |
| ABCB1 |  |
| GGH |  |
| UGT1A7 |  |
| ENOSF1 |  |
| ABCC3 |  |
| UGT1A8 |  |
| HBB |  |
| CYP1A2 |  |
| HMGCR |  |
| SHMT1 |  |
| FOXP3 |  |
| ABCC2 |  |
| AMPD1 |  |
| DOK5 |  |
| FTO |  |
| CDKN1B |  |
| UGT1A6 |  |
| UGT1A4 |  |
| HSD17B10 |  |
| UGT1A3 |  |
| ABCC4 |  |
| FOLH1 |  |
| SLC22A9 |  |
| SLC19A1 |  |
| TP53 |  |
| ITPA |  |
| BMP7 |  |
| GSTM1 |  |
| ABCC1 |  |
| NALCN |  |
| S100A12 |  |
| NUDT15 |  |
| TNFAIP3 |  |
| SLC16A7 |  |
| ATAD5 |  |
| RB1 |  |
| NOTCH1 |  |
| ADORA2A |  |
| HGF |  |
| GSTP1 |  |
| FPGS |  |
| TPMT |  |
| SULT2A1 |  |
| SLCO1B3 |  |
| NOS1 |  |
| GCG |  |
| HLA-C |  |
| GABBR2 |  |
| KIR2DS4 |  |
| SLC22A11 |  |
| BDNF |  |
| SLCO1A2 |  |
| RYR1 |  |
| FOLR1 |  |
| ATP5E |  |
| TAT |  |
| GLS |  |
| NCOA3 |  |
| TLR4 |  |
| CYP2C19 |  |
| DHODH |  |
| PPARG |  |
| SLCO1B1 |  |
| KEAP1 |  |
| HLA-DQA1 |  |
| IDH1 |  |
| IKZF1 |  |
| HLA-E |  |
| FGFR4 |  |
| GATA3 |  |
| DROSHA |  |
| NTRK1 |  |
| PTH |  |
| SLC6A3 |  |
| MPHOSPH8 |  |
| MTRR |  |
| UGT1A9 |  |
| DDRGK1 |  |
| ADA |  |
| HLA-G |  |
| LINC00251 |  |
| SOD2 |  |
| EHMT2 |  |
| UGT1A1 |  |
| NOS3 |  |
| ARID5B |  |
| SLC46A1 |  |
| COMT |  |
| SLAMF1 |  |
| POLB |  |
| SLC22A1 |  |
| APEX1 |  |
| TAC1 |  |
| SPECC1L |  |
| ALOX5 |  |
| ICAM3 |  |
| UGT1A5 |  |
| POLI |  |
| FLT3 |  |
| SLC22A8 |  |
| UGT1A10 |  |
| NR1I2 |  |
| HLA-DRB1 |  |
| S1PR1 |  |
| ABCG2 |  |

**Table S4.** Genes associated with each of the diseases compiled from the DisGeNET database

| **Anxiety** | **Depression** |
| --- | --- |
| SLC6A4 | CRH |
| COMT | CRHR1 |
| DRD2 | DRD2 |
| FMR1 | NR3C1 |
| BDNF | HTR2A |
| LINC02210-CRHR1 | IL1B |
| SNCA | IL6 |
| HTR2A | NPY |
| MECP2 | NTRK2 |
| GTF2I | BDNF |
| DAO | S100A10 |
| GABRG2 | S100B |
| GRN | SLC6A2 |
| COX2 | SLC6A4 |
| PINK1 | TPH1 |
| SOD1 | GSK3B |
| TP53 | NGF |
| PDE10A | CHRM2 |
| PARK7 | ADRA2A |
| LRRK2 | DBH |
| CYP17A1 | DRD1 |
| C9orf72 | CRHBP |
| ERBB4 | CRHR2 |
| TARDBP | IL6R |
| NIPBL | PPP1R1B |
| GNAS | HDAC5 |
| GNRHR | PDE4D |
| LAMA1 | TH |
| HSPG2 | TPH2 |
| HCN1 | CNR1 |
| MEN1 | COMT |
| ND6 | CREB1 |
| TLR9 | CYP2D6 |
| AHI1 | DRD4 |
| PSEN1 | ESR1 |
| TACR3 | FKBP5 |
| XK | GAD1 |
| EHMT1 | DISC1 |
| KISS1R | GRIN2B |
| SGCE | HCRT |
| TMEM161B-AS1 | HTR1A |
| RNF103-CHMP3 | HTR2C |
| STIMATE-MUSTN1 | IFNG |
| BORCS7-ASMT | IGF1 |
| USH1C | IDO1 |
| PQBP1 | AR |
| OPTN | KCNK2 |
| LINC01876 | LEP |
| STUB1 | MAOA |
| BAIAP2 | NR3C2 |
| CIB2 | MTHFR |
| DEAF1 | OPRK1 |
| LINC02503 | OXTR |
| STAG2 | P2RX7 |
| RAI1 | GAL |
| PPARGC1A | ABCB1 |
| SLC38A3 | POMC |
| TLK2 | PTGS2 |
| ADCY5 | SLC6A3 |
| KPTN | TAC1 |
| NISCH | TNF |
| POLG2 | VEGFA |
| PRRT2 | ARTN |
| STX1B | IL1A |
| TWF2 | IL18 |
| CLCN4 | REN |
| CLN3 | CLOCK |
| TPH2 | GRIA1 |
| USH1G | CXCL8 |
| CNR1 | CYP2C19 |
| PROKR2 | DRD3 |
| CPOX | PCLO |
| SGCZ | COX2 |
| CRH | CACNA1C |
| CRHR1 | BICC1 |
| CRP | GFAP |
| NAGS | HP |
| DENND1B | HTR1B |
| DCTN1 | ARNTL |
| DNMT3A | PDE4A |
| DSCAM | SLC18A2 |
| DUSP6 | TNFRSF1A |
| TOR1A | CNR2 |
| ARID2 | AKT1 |
| ELN | HTR3A |
| ARHGAP27 | MAOB |
| TSNARE1 | NOS2 |
| EPHA4 | NOS3 |
| ESR1 | PDE4B |
| ACSL4 | MAPK3 |
| BPTF | BRCA1 |
| JMJD1C | FTO |
| FGF8 | PER2 |
| FGFR1 | CHRNA4 |
| FKBP5 | GNB3 |
| ARSG | TNFRSF1B |
| FAN1 | TRH |
| NT5C2 | VGF |
| UNC13A | HOMER1 |
| IQSEC2 | CARTPT |

| **Parkinson's disease** | **Schizophrenia** |
| --- | --- |
| SNCA | AKT1 |
| PARK7 | MAGI2 |
| DDC | CHRNA7 |
| DRD2 | GRIN2B |
| ATP13A2 | HTR2A |
| MAOB | NOS1 |
| PRKN | RELN |
| PINK1 | SRR |
| SLC18A2 | CSMD1 |
| TH | TCF4 |
| DRD1 | SHANK3 |
| IGF1R | NRXN1 |
| LRRK2 | RTN4R |
| GAK | SP4 |
| GBA | SETD1A |
| MAPT | PPP3R1 |
| BST1 | SYNGAP1 |
| HLA-DRA | MDK |
| PARK16 | NRG3 |
| DNM1L | CNR1 |
| PPARGC1A | DISC1 |
| CP | GRM5 |
| CYP2D6 | GSK3B |
| GDNF | ZDHHC8 |
| GFAP | APOE |
| TMEM230 | NR4A2 |
| GSTM1 | SLC6A3 |
| HFE | DTNBP1 |
| HMOX1 | PPP1R1B |
| HSPA9 | KMO |
| IL6 | CPLX2 |
| MAOA | BACE1 |
| MTHFR | MBP |
| NOS1 | TAAR1 |
| ABCB1 | MAP6 |
| VPS35 | PTGS2 |
| BDNF | PLCB1 |
| SLC6A3 | LRRTM1 |
| SOD1 | MAP2K7 |
| SOD2 | NLGN2 |
| TNF | SLC6A1 |
| GSTP1 | YWHAH |
| DDIT4 | MTOR |
| CYP2E1 | AVPR1A |
| MAP3K5 | ZIC2 |
| NGF | GNAS |
| NQO1 | BECN1 |
| AIF1 | CHI3L1 |
| GSTA4 | COMT |
| IGF2 | DRD3 |
| TRPM2 | BRD1 |
| HLA-DRB5 | DAOA |
| IGF2R | DISC2 |
| INS | ANK3 |
| ENO2 | GRIA1 |
| FBP1 | GRM3 |
| FCER2 | NRG1 |
| GPX1 | HLA-DRB1 |
| HGF | MTHFR |
| HSPA1A | NCAM1 |
| INSR | NOTCH4 |
| MIR181C | NRGN |
| MAG | PDE4B |
| MTA1 | ABCB1 |
| BAG5 | PRODH |
| TCL1B | SLC1A1 |
| ADARB2 | TBX1 |
| COL19A1 | CACNA1C |
| SLC2A14 | ZNF804A |
| EDN1 | FOXP2 |
| FGB | FEZ1 |
| CNTNAP2 | NOS1AP |
| GSK3B | SRGAP3 |
| HBG1 | SYN2 |
| HSPA4 | MED12 |
| HSPA8 | CHRNA5 |
| IL1B | HLA-DQB1 |
| DRAXIN | VRK2 |
| KCNJ4 | HLA-A |
| MAP2 | NDE1 |
| CEACAM6 | DLG2 |
| NR4A2 | GABBR1 |
| PITX3 | MIR137HG |
| NCAPG2 | ARVCF |
| SLC30A10 | TCF7L2 |
| NECTIN2 | FYN |
| RPL6 | CNNM2 |
| RPL23A | YWHAE |
| RPS8 | TSPAN18 |
| TALDO1 | TSNARE1 |
| TFAM | MPC2 |
| RPL14 | ITIH3 |
| OPTN | NTRK3 |
| SPR | PBRM1 |
| TWNK | MAD1L1 |
| ATG7 | DCC |
| GRK5 | ESR2 |
| HTR1A | NKAPL |
| PENK | HLA-B |
| GDF5 | MSRA |

| **Asthma** | **Hypertension** |
| --- | --- |
| TBX21 | PTGIS |
| IL5 | NOS3 |
| IL13 | AGT |
| CCL11 | AGTR1 |
| TGFB1 | GNB3 |
| ICAM1 | ADD1 |
| IL4 | NOS2 |
| IL6 | RGS5 |
| MMP9 | ATP1B1 |
| NOS2 | ECE1 |
| CCL2 | CALY |
| CCL5 | CYP3A5 |
| SCGB3A2 | GATA5 |
| ADRB2 | UMOD |
| ALOX5 | SLC12A3 |
| HLA-DQB1 | MTHFR |
| HLA-DRB1 | ADRB1 |
| HLA-G | ACE2 |
| GSDMB | WNK4 |
| TNF | ACE |
| TSLP | HSD11B2 |
| IL33 | EDN1 |
| IL1RL1 | REN |
| ORMDL3 | CYP11B2 |
| HLA-DQA1 | PGR-AS1 |
| IL6R | ADRB2 |
| RAD50 | GRK4 |
| PDE4D | TNF |
| IKZF3 | KNG1 |
| HNMT | LEP |
| CDHR3 | ATM |
| KIF3A | IL6 |
| PYHIN1 | SLC33A1 |
| MUC7 | NR3C2 |
| PLA2G7 | CHGA |
| DNAH5 | ALDH2 |
| TNIP1 | ADIPOQ |
| WDR36 | RNU1-4 |
| PTGDR2 | INSR |
| CCR3 | NEDD4L |
| PARP1 | CLCNKB |
| EDN1 | APOE |
| GATA3 | STK39 |
| GSTM1 | LPL |
| GSTP1 | TGFB1 |
| HLA-DPB1 | KLK1 |
| TNC | ALB |
| IL1B | ADRB3 |
| IL1RN | SYNE1 |
| IL2RA | CYP4A11 |
| IL4R | APLN |
| ARG1 | CRP |
| NPSR1 | ATP2B1 |
| RNASE3 | VEGFA |
| STAT6 | AGTR2 |
| VEGFA | SELE |
| CAT | CYP2J2 |
| CD14 | BDKRB2 |
| ARG2 | ERAP1 |
| AREG | NPR3 |
| NQO1 | SERPINE1 |
| MMP1 | LEPR |
| ALDH2 | CYP2C9 |
| CTNNA3 | SPP1 |
| SOD1 | PPARG |
| BCL2 | CCHCR1 |
| RUNX3 | GSTK1 |
| HMOX1 | SLCO6A1 |
| PLAU | ADM |
| TRPA1 | SELENBP1 |
| IFNL3 | MFN2 |
| NPY | GCGR |
| PTEN | HCCAT5 |
| MMP10 | HMOX1 |
| ADCY2 | WNK1 |
| ADCYAP1R1 | SHBG |
| MYB | HSPA4 |
| BGLAP | TRPC3 |
| SPRR2B | EDNRA |
| TIMP3 | SLC7A1 |
| CCL26 | TSC1 |
| DNMT1 | TNFRSF1B |
| EGFR | VDR |
| NR3C1 | TH |
| HSD11B2 | HLA-DRB1 |
| IL10 | HPGDS |
| IL17A | HP |
| KRT19 | LPA |
| MUC5AC | CYBA |
| SERPINE1 | EMILIN1 |
| IL22 | TESC |
| PDE4B | KCNJ11 |
| SERPINA1 | RGS2 |
| PPP2CA | UTS2 |
| PTGS2 | NPR1 |
| DPP10 | CYP2D6 |
| CCL17 | POMC |
| SCGB1A1 | AGER |
| CXCL14 | DRD1 |
| AGER | GSTT1 |

| **Type 2 diabetes** | **Obesity** |
| --- | --- |
| HNF4A | MC4R |
| PAX4 | PPARG |
| PPP1R3A | POMC |
| AKT2 | CPE |
| GCK | AGRP |
| IRS1 | SH2B1 |
| KCNJ11 | LEP |
| HNF1A | LEPR |
| NEUROD1 | PCSK1 |
| SLC2A4 | ENPP1 |
| HNF1B | SIM1 |
| ADCY5 | UCP3 |
| CAPN10 | NR0B2 |
| HMGA1 | BBS4 |
| INS | MKKS |
| INSR | BBS1 |
| PDX1 | CNR1 |
| ENPP1 | CRP |
| PPARG | ESR1 |
| CDKAL1 | SIRT3 |
| SLC2A2 | SIRT1 |
| ABCC8 | HSD11B1 |
| TCF7L2 | IL6 |
| TGFB1 | IRS1 |
| WFS1 | LBP |
| IRS2 | LPL |
| LIPC | CCL2 |
| BCL2 | ADIPOQ |
| MAPK8IP1 | ICAM1 |
| PPARGC1A | CYP2E1 |
| EDN1 | F2 |
| EDNRA | PMCH |
| SIRT1 | ACSL1 |
| HMOX1 | FOXO3 |
| HP | GPX3 |
| ICAM1 | HK2 |
| LEP | NPY5R |
| LEPR | CD40 |
| NOS3 | HADH |
| TNF | ADRB2 |
| UCP2 | ADRB3 |
| ADIPOQ | GLUL |
| TNFRSF1A | APOE |
| NOS2 | INS |
| KL | NPY1R |
| CPT1A | GHRL |
| SNAP25 | INPP5E |
| EDNRB | TF |
| ATP2A2 | FTO |
| IGF2BP2 | NEIL1 |
| SLC30A8 | CARTPT |
| GLIS3 | ADCY3 |
| JAZF1 | GNAS |
| GCGR | NPC1 |
| GCKR | PRKAR1A |
| GLP1R | AHR |
| KCNQ1 | SDCCAG8 |
| MTNR1B | MYT1L |
| RETN | NTRK2 |
| PTPN1 | SDC3 |
| FTO | KCNMA1 |
| THADA | RAI1 |
| HMG20A | MRAP2 |
| NOTCH2 | BBS9 |
| PROX1 | BBS2 |
| GPD2 | VPS13B |
| UBE2E2 | AFF4 |
| KCNK16 | HOXB5 |
| AP3S2 | PACS1 |
| MAEA | BBS10 |
| GRB14 | PHF6 |
| PEPD | NAMPT |
| CMIP | TTC8 |
| ITGA1 | TMEM18 |
| PAM | BBS5 |
| PLEKHA1 | GNPDA2 |
| ZFAND3 | BBS12 |
| DGKD | DPYD |
| CCND2 | RMST |
| PSMD6 | AKT1 |
| KSR2 | FAAH |
| NFATC2 | FASN |
| ZNF257 | NEGR1 |
| KLF14 | FGF21 |
| CYBA | GCG |
| ETS1 | GH1 |
| FBN1 | GNB3 |
| CNKSR2 | HCRT |
| JADE2 | HTR2C |
| AUTS2 | IGF2 |
| EPC2 | AQP7 |
| FGF21 | LDLR |
| GCG | MC3R |
| GNB3 | MMP9 |
| GP2 | SERPINE1 |
| GSTM1 | ANGPTL4 |
| HHEX | PLIN1 |
| IAPP | PPARA |
| IDE | PPARD |
| IGF2 | MKS1 |

| **Rheumatoid arthritis** | **Osteoporosis** |
| --- | --- |
| PTPN22 | LRP5 |
| SLC22A4 | COL1A1 |
| TNF | VDR |
| CRP | WNT1 |
| IL6ST | COL1A2 |
| CCR6 | PTH |
| FCGR2A | TGFB1 |
| PADI4 | TNFSF11 |
| HLA-DPB1 | CYP19A1 |
| HLA-DRB1 | TNFRSF11B |
| IL2RA | CALCR |
| IL6R | PDLIM4 |
| IL10 | GORAB |
| IRF5 | ANTXR2 |
| CIITA | ESR1 |
| STAT4 | ESR2 |
| TNFAIP3 | IGF1 |
| TRAF1 | IL6 |
| TRAF6 | AR |
| CD28 | KL |
| CD40 | REN |
| PTPN2 | SOD2 |
| TYK2 | GPC6 |
| RUNX1 | POMC |
| PTPRC | CA2 |
| AGER | GAPDH |
| IL2RB | GPX1 |
| NFKBIL1 | GSN |
| AFF3 | ANXA2 |
| REL | LTF |
| BLK | P4HB |
| CDK6 | ZDHHC13 |
| CD244 | TPM4 |
| MMEL1 | HDAC5 |
| ANKRD55 | TUBA1B |
| GATA3 | CAP1 |
| NFKBIE | CCT2 |
| PLD4 | PRDX3 |
| DNASE1L3 | ADCY5 |
| KIF5A | PARK7 |
| ARID5B | MGLL |
| RASGRP1 | CYP24A1 |
| CSF2 | ENO1 |
| CTLA4 | FGA |
| ACAN | FGB |
| RCAN1 | SIRT1 |
| AHR | DAAM2 |
| SPRED2 | PGLS |
| ENO1 | GPD2 |
| ANXA3 | RAB7B |
| HLA-DQA2 | IDH2 |
| IFNG | IL6R |
| CCN1 | LEP |
| IL1B | ATIC |
| IL1RN | PNP |
| IL6 | OXCT1 |
| CXCL8 | PKM |
| IL18 | PLEK |
| MIF | PSMA2 |
| MMP2 | PSMA5 |
| MPO | RSU1 |
| MTHFR | BGLAP |
| NCF2 | CLEC11A |
| TNFRSF11B | TLN1 |
| FOXP3 | ACTG1 |
| IL23A | TPI1 |
| ABCB1 | U2AF1 |
| PON1 | VCL |
| PTGS2 | PIR |
| CCL21 | WDR1 |
| SLC11A1 | GH1 |
| STAT1 | IL1B |
| TLR2 | ACE |
| VEGFA | AGER |
| CXCR4 | IRS2 |
| CAT | IRAK3 |
| ADIPOQ | MAPK14 |
| CCN2 | GHR |
| TNFSF14 | IGF2 |
| DHFR | CIITA |
| TNFRSF14 | MYC |
| TAGAP | PCNA |
| IRAK1 | BAX |
| COL2A1 | BCL2 |
| ALOX5 | DSPP |
| GC | DKK1 |
| HOXD13 | ID4 |
| NCF1 | IFNGR1 |
| TXNDC5 | IRS1 |
| GRK2 | PLS3 |
| FPGS | PTGER4 |
| IGFBP3 | BMP2 |
| FASLG | RUNX2 |
| ATIC | TNFRSF11A |
| PRKCQ | NOTCH2 |
| PTGS1 | PPARG |
| CCL8 | CYP17A1 |
| SOD2 | LRP6 |
| GGH | SOX6 |
| B3GNT2 | ALB |

| **Chronic obstructive pulmonary disease** | **Heart failure** |
| --- | --- |
| MMP1 | PPARGC1A |
| HMOX1 | ADRB1 |
| SFTPD | GRK2 |
| HDAC2 | ACE |
| MMP9 | AGT |
| TGFB1 | AGTR1 |
| FAM13A | EDN1 |
| SERPINA1 | HIF1A |
| DSP | IL6 |
| ELN | NOS3 |
| EPHX1 | NPPA |
| MTCL1 | NPPB |
| IL6 | ATP2A2 |
| CXCL8 | TNF |
| EEFSEC | CCL2 |
| TNF | HMOX1 |
| NOS3 | SOD2 |
| CYP1A1 | MSTN |
| SOD3 | ADRB3 |
| FOXO3 | HTR2B |
| NOS2 | RAC1 |
| CXCL1 | CRP |
| CYP1A2 | CCN2 |
| HTR2A | EDNRA |
| TRPV4 | ALB |
| RAPGEF3 | GCG |
| TP53 | NRG1 |
| CXCL2 | IL1B |
| MMP14 | NR3C2 |
| KLF5 | PIK3CG |
| TNNT2 | AVP |
| CD8A | PTH |
| IL17A | REN |
| MIR218-2 | VEGFA |
| MUC5AC | ADIPOQ |
| NFE2L2 | GDF15 |
| TIMP1 | SIRT1 |
| VEGFA | PPARG |
| EGFR | PPP1R1A |
| CASP3 | ADRA2C |
| SERPINE1 | CYBB |
| MUC1 | NFE2L2 |
| PLAU | PTGS2 |
| SLPI | NOX4 |
| CCN2 | SLC9A1 |
| MPO | TNFRSF1A |
| STAT4 | UCN2 |
| DDIT3 | NOS2 |
| SMAD4 | SERPINE1 |
| BNIP3 | PON1 |
| CASP12 | PRL |
| CFTR | EDNRB |
| CHRNA3 | VWF |
| CHRNA5 | PEBP1 |
| NQO1 | RETN |
| AGER | TLR2 |
| ERBB3 | CS |
| FAIM2 | CSF2 |
| FAS | EPHX2 |
| FASLG | GSK3B |
| IREB2 | IFNG |
| PLAUR | CXCL8 |
| SMPD3 | NPR1 |
| CASP8 | OLR1 |
| CHRNB4 | POMC |
| CTLA4 | AVPR2 |
| MMP3 | RBP4 |
| HSPA4 | XDH |
| CHRM3 | CSF3 |
| SCGB1A1 | CFD |
| PLB1 | GATM |
| PRTN3 | ACACA |
| THSD4 | ACADS |
| CYBA | SOD3 |
| HSPA1A | TNNT2 |
| HTR4 | UCP1 |
| NFKB1 | NRIP1 |
| RNF150 | CAT |
| TNS1 | HAND2 |
| CDH13 | ROCK2 |
| HYKK | FBLN5 |
| CYBB | DSTN |
| DNAAF4 | CIDEA |
| PSORS1C1 | ADM |
| DLG2 | ADRB2 |
| DNAH5 | FASN |
| DOCK1 | BAMBI |
| PDZD2 | NOX1 |
| NPNT | GPX4 |
| HLA-DPA1 | CXCL2 |
| HLA-DPB1 | APCS |
| HSPA1B | HSPB1 |
| HSPA1L | APOC1 |
| INPP5D | IGF1 |
| KCNK1 | INS |
| ARHGEF38 | ITGB1 |
| INTS12 | LGALS3 |
| CSMD1 | MME |
| WIPF1 | MMP9 |
| FTO | ACLY |

| **Schizophrenia** | **Asthma** |
| --- | --- |
| AKT1 | TBX21 |
| MAGI2 | IL5 |
| CHRNA7 | IL13 |
| GRIN2B | CCL11 |
| HTR2A | TGFB1 |
| NOS1 | ICAM1 |
| RELN | IL4 |
| SRR | IL6 |
| CSMD1 | MMP9 |
| TCF4 | NOS2 |
| SHANK3 | CCL2 |
| NRXN1 | CCL5 |
| RTN4R | SCGB3A2 |
| SP4 | ADRB2 |
| SETD1A | ALOX5 |
| PPP3R1 | HLA-DQB1 |
| SYNGAP1 | HLA-DRB1 |
| MDK | HLA-G |
| NRG3 | GSDMB |
| CNR1 | TNF |
| DISC1 | TSLP |
| GRM5 | IL33 |
| GSK3B | IL1RL1 |
| ZDHHC8 | ORMDL3 |
| APOE | HLA-DQA1 |
| NR4A2 | IL6R |
| SLC6A3 | RAD50 |
| DTNBP1 | PDE4D |
| PPP1R1B | IKZF3 |
| KMO | HNMT |
| CPLX2 | CDHR3 |
| BACE1 | KIF3A |
| MBP | PYHIN1 |
| TAAR1 | MUC7 |
| MAP6 | PLA2G7 |
| PTGS2 | DNAH5 |
| PLCB1 | TNIP1 |
| LRRTM1 | WDR36 |
| MAP2K7 | PTGDR2 |
| NLGN2 | CCR3 |
| SLC6A1 | PARP1 |
| YWHAH | EDN1 |
| MTOR | GATA3 |
| AVPR1A | GSTM1 |
| ZIC2 | GSTP1 |
| GNAS | HLA-DPB1 |
| BECN1 | TNC |
| CHI3L1 | IL1B |
| COMT | IL1RN |
| DRD3 | IL2RA |
| BRD1 | IL4R |
| DAOA | ARG1 |
| DISC2 | NPSR1 |
| ANK3 | RNASE3 |
| GRIA1 | STAT6 |
| GRM3 | VEGFA |
| NRG1 | CAT |
| HLA-DRB1 | CD14 |
| MTHFR | ARG2 |
| NCAM1 | AREG |
| NOTCH4 | NQO1 |
| NRGN | MMP1 |
| PDE4B | ALDH2 |
| ABCB1 | CTNNA3 |
| PRODH | SOD1 |
| SLC1A1 | BCL2 |
| TBX1 | RUNX3 |
| CACNA1C | HMOX1 |
| ZNF804A | PLAU |
| FOXP2 | TRPA1 |
| FEZ1 | IFNL3 |
| NOS1AP | NPY |
| SRGAP3 | PTEN |
| SYN2 | MMP10 |
| MED12 | ADCY2 |
| CHRNA5 | ADCYAP1R1 |
| HLA-DQB1 | MYB |
| VRK2 | BGLAP |
| HLA-A | SPRR2B |
| NDE1 | TIMP3 |
| DLG2 | CCL26 |
| GABBR1 | DNMT1 |
| MIR137HG | EGFR |
| ARVCF | NR3C1 |
| TCF7L2 | HSD11B2 |
| FYN | IL10 |
| CNNM2 | IL17A |
| YWHAE | KRT19 |
| TSPAN18 | MUC5AC |
| TSNARE1 | SERPINE1 |
| MPC2 | IL22 |
| ITIH3 | PDE4B |
| NTRK3 | SERPINA1 |
| PBRM1 | PPP2CA |
| MAD1L1 | PTGS2 |
| DCC | DPP10 |
| ESR2 | CCL17 |
| NKAPL | SCGB1A1 |
| HLA-B | CXCL14 |
| MSRA | AGER |

| **Schizophrenia** | **Rheumatoid arthritis** |
| --- | --- |
| AKT1 | PTPN22 |
| MAGI2 | SLC22A4 |
| CHRNA7 | TNF |
| GRIN2B | CRP |
| HTR2A | IL6ST |
| NOS1 | CCR6 |
| RELN | FCGR2A |
| SRR | PADI4 |
| CSMD1 | HLA-DPB1 |
| TCF4 | HLA-DRB1 |
| SHANK3 | IL2RA |
| NRXN1 | IL6R |
| RTN4R | IL10 |
| SP4 | IRF5 |
| SETD1A | CIITA |
| PPP3R1 | STAT4 |
| SYNGAP1 | TNFAIP3 |
| MDK | TRAF1 |
| NRG3 | TRAF6 |
| CNR1 | CD28 |
| DISC1 | CD40 |
| GRM5 | PTPN2 |
| GSK3B | TYK2 |
| ZDHHC8 | RUNX1 |
| APOE | PTPRC |
| NR4A2 | AGER |
| SLC6A3 | IL2RB |
| DTNBP1 | NFKBIL1 |
| PPP1R1B | AFF3 |
| KMO | REL |
| CPLX2 | BLK |
| BACE1 | CDK6 |
| MBP | CD244 |
| TAAR1 | MMEL1 |
| MAP6 | ANKRD55 |
| PTGS2 | GATA3 |
| PLCB1 | NFKBIE |
| LRRTM1 | PLD4 |
| MAP2K7 | DNASE1L3 |
| NLGN2 | KIF5A |
| SLC6A1 | ARID5B |
| YWHAH | RASGRP1 |
| MTOR | CSF2 |
| AVPR1A | CTLA4 |
| ZIC2 | ACAN |
| GNAS | RCAN1 |
| BECN1 | AHR |
| CHI3L1 | SPRED2 |
| COMT | ENO1 |
| DRD3 | ANXA3 |
| BRD1 | HLA-DQA2 |
| DAOA | IFNG |
| DISC2 | CCN1 |
| ANK3 | IL1B |
| GRIA1 | IL1RN |
| GRM3 | IL6 |
| NRG1 | CXCL8 |
| HLA-DRB1 | IL18 |
| MTHFR | MIF |
| NCAM1 | MMP2 |
| NOTCH4 | MPO |
| NRGN | MTHFR |
| PDE4B | NCF2 |
| ABCB1 | TNFRSF11B |
| PRODH | FOXP3 |
| SLC1A1 | IL23A |
| TBX1 | ABCB1 |
| CACNA1C | PON1 |
| ZNF804A | PTGS2 |
| FOXP2 | CCL21 |
| FEZ1 | SLC11A1 |
| NOS1AP | STAT1 |
| SRGAP3 | TLR2 |
| SYN2 | VEGFA |
| MED12 | CXCR4 |
| CHRNA5 | CAT |
| HLA-DQB1 | ADIPOQ |
| VRK2 | CCN2 |
| HLA-A | TNFSF14 |
| NDE1 | DHFR |
| DLG2 | TNFRSF14 |
| GABBR1 | TAGAP |
| MIR137HG | IRAK1 |
| ARVCF | COL2A1 |
| TCF7L2 | ALOX5 |
| FYN | GC |
| CNNM2 | HOXD13 |
| YWHAE | NCF1 |
| TSPAN18 | TXNDC5 |
| TSNARE1 | GRK2 |
| MPC2 | FPGS |
| ITIH3 | IGFBP3 |
| NTRK3 | FASLG |
| PBRM1 | ATIC |
| MAD1L1 | PRKCQ |
| DCC | PTGS1 |
| ESR2 | CCL8 |
| NKAPL | SOD2 |
| HLA-B | GGH |
| MSRA | B3GNT2 |

| **Multiple sclerosis** | **Peroxisomal disorders** |
| --- | --- |
| CLEC16A | HSD17B4 |
| HLA-DRB1 | PIPOX |
| IL2RA | PEX6 |
| IL7R | PEX1 |
| TNFRSF1A | ABCD1 |
| CD40 | TRIM37 |
| CD58 | ACOX1 |
| TYK2 | PEX5 |
| KIF1B | PHEX |
| HLA-DRA | DNM1L |
| CBLB | PAF1 |
| STAT4 | PEX2 |
| TNFAIP3 | DHRS11 |
| TNFSF14 | PEX3 |
| CD6 | ACBD5 |
| NLRP3 | PEX12 |
| HLA-DPB1 | PEX10 |
| HLA-DQB1 | HSD17B7 |
| ICAM1 | LPA |
| IRF8 | HSD17B13 |
| IFNB1 | HADH |
| IFNG | AMACR |
| APOE | SIRT1 |
| IL1B | EPAS1 |
| IL1RN | ABCG2 |
| IL7 |  |
| IL10 |  |
| IL17A |  |
| P2RX7 |  |
| VDR |  |
| CNR1 |  |
| GC |  |
| CASP1 |  |
| SLC11A1 |  |
| CLDN11 |  |
| POMC |  |
| PDCD1 |  |
| NECTIN2 |  |
| SELE |  |
| VCAM1 |  |
| IL12A |  |
| RBPJ |  |
| MCAM |  |
| KCNJ10 |  |
| BCHE |  |
| TNF |  |
| PRF1 |  |
| CYP27B1 |  |
| LRCH1 |  |
| HLA-A |  |
| HLA-B |  |
| HLA-DQA1 |  |
| STAT3 |  |
| CD226 |  |
| IL23R |  |
| GPC5 |  |
| NR1H3 |  |
| MAPK1 |  |
| IFIH1 |  |
| EVI5 |  |
| ATG5 |  |
| ERG |  |
| KIF21B |  |
| TAP2 |  |
| CCR6 |  |
| PTPN22 |  |
| MGAT5 |  |
| FCRL3 |  |
| TAGAP |  |
| CCR3 |  |
| ITGAM |  |
| NOTCH4 |  |
| RGS1 |  |
| CXCR5 |  |
| CD86 |  |
| MERTK |  |
| SP140 |  |
| GALC |  |
| GFI1 |  |
| HSPA1L |  |
| TMEM39A |  |
| PTPN2 |  |
| RPS6KB1 |  |
| MMEL1 |  |
| ANKRD55 |  |
| TSFM |  |
| MPHOSPH9 |  |
| MALT1 |  |
| ADAD1 |  |
| ZBTB46 |  |
| CTSH |  |
| ZNF433 |  |
| ERBB3 |  |
| ICOSLG |  |
| ICOS |  |
| HLA-DMB |  |
| HLA-DQA2 |  |
| HLA-DRB4 |  |
| IRGM |  |
| IFNGR2 |  |

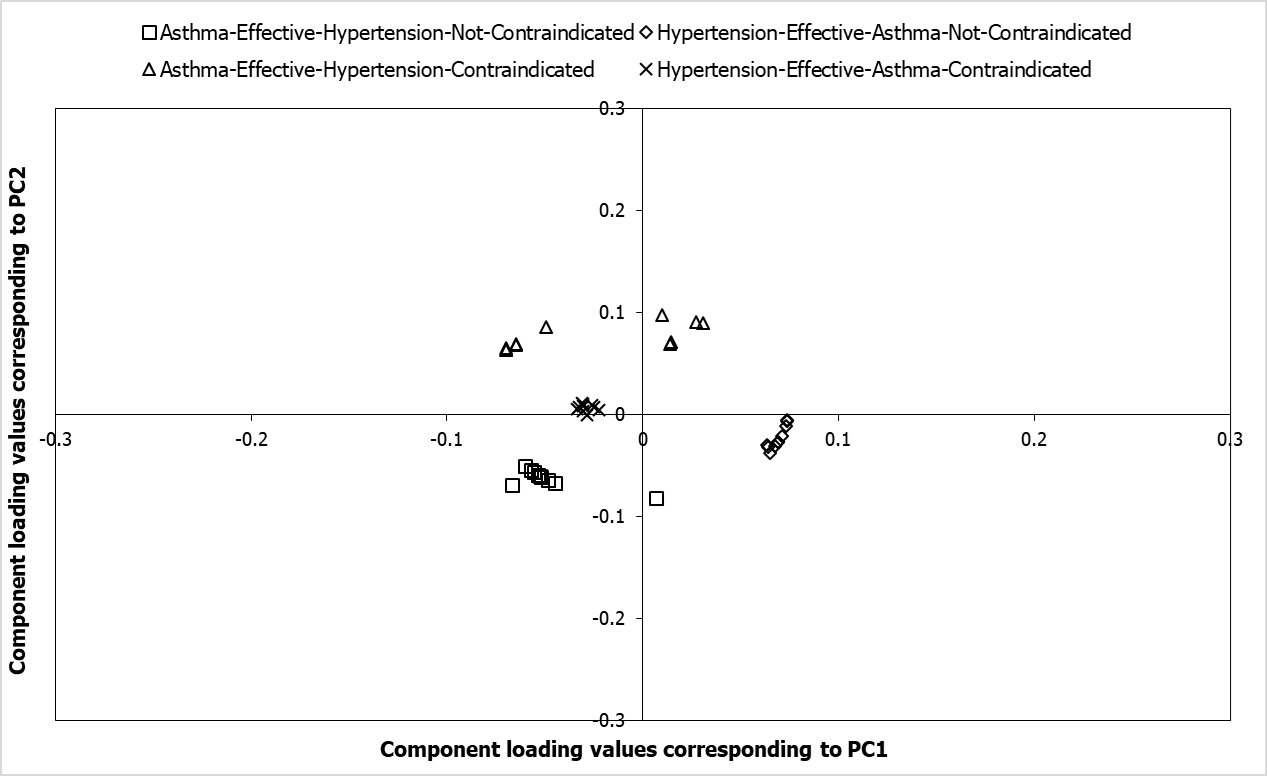
**Figure S1: Pathways associated with the target networks of asthma and hypertension drugs.** Component loading values of 378 pathways associated with the drug target networks (DTNs) of asthma and hypertension corresponding to PC1 and PC2 have been plotted along the X and Y axes respectively. PCA was performed with the p-values of enrichment of the pathways significantly associated (p-value < 0.05) with the DTNs of asthma and hypertension. These values were transformed to –log_10_P values, which were then assembled into a data matrix containing pathways as rows and DTNs as columns. Unit variance scaling was applied across this matrix. Single value decomposition (SVD) with imputation was used to extract the principal components (PCs). The component loading values shown in the figure correspond to component scores of 4 DTNs along PC1 and PC2 that explain 42.9% and 37.2% of the total variance respectively. The top-10 pathways that appeared to be highly related to each of the 4 DTNs, which were obtained after computing the Euclidean distance between the component loading values and the component scores, are shown as square-shaped data points for the DTN of drugs effective in asthma and not contraindicated in hypertension, diamond-shaped data points for the DTN of drugs effective in hypertension and not contraindicated in asthma, triangle-shaped data points for the DTN of drugs effective in asthma and contraindicated in hypertension and cross-mark-shaped data points for the DTN of drugs effective in hypertension and contraindicated in asthma.

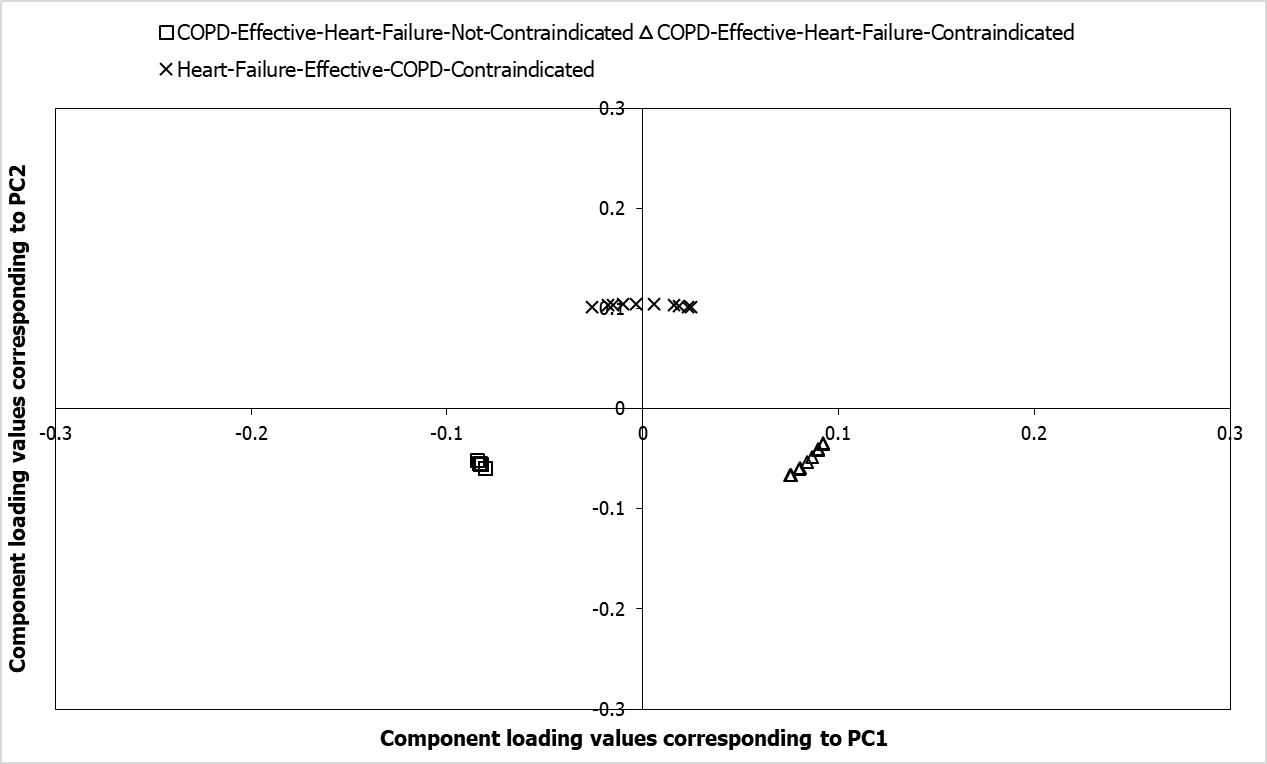
**Figure S2: Pathways associated with the target networks of chronic obstructive pulmonary disease and heart failure drugs.** Component loading values of 233 pathways associated with the drug target networks (DTNs) of chronic obstructive pulmonary disease (COPD) and heart failure corresponding to PC1 and PC2 have been plotted along the X and Y axes respectively. PCA was performed with the p-values of enrichment of the pathways significantly associated (p-value < 0.05) with the DTNs of COPD and heart failure. These values were transformed to –log_10_P values, which were then assembled into a data matrix containing pathways as rows and DTNs as columns. Unit variance scaling was applied across this matrix. Single value decomposition (SVD) with imputation was used to extract the principal components (PCs). The component loading values shown in the figure correspond to component scores of 4 DTNs along PC1 and PC2 that explain 53.6% and 46.4% of the total variance respectively. The top-10 pathways that appeared to be highly related to each of the 4 DTNs, which were obtained after computing the Euclidean distance between the component loading values and the component scores, are shown as square-shaped data points for the DTN of drugs effective in COPD and not contraindicated in heart failure, diamond-shaped data points for the DTN of drugs effective in heart failure and not contraindicated in COPD, triangle-shaped data points for the DTN of drugs effective in COPD and contraindicated in heart failure and cross-mark-shaped data points for the DTN of drugs effective in heart failure and contraindicated in COPD.

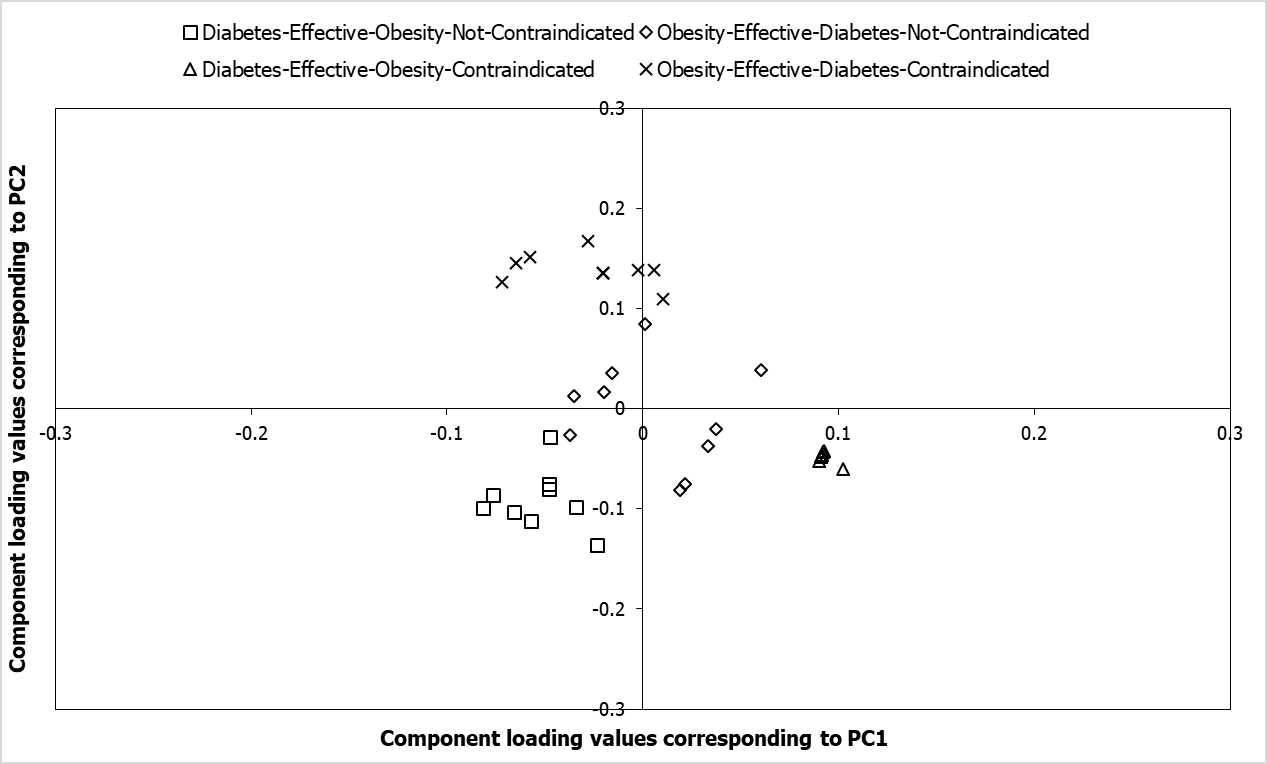

**Figure S3: Pathways associated with the target networks of type 2 diabetes and obesity drugs.** Component loading values of 270 pathways associated with the drug target networks (DTNs) of type 2 diabetes and obesity corresponding to PC1 and PC2 have been plotted along the X and Y axes respectively. PCA was performed with the p-values of enrichment of the pathways significantly associated (p-value < 0.05) with the DTNs of type 2 diabetes and obesity. These values were transformed to –log_10_P values, which were then assembled into a data matrix containing pathways as rows and DTNs as columns. Unit variance scaling was applied across this matrix. Single value decomposition (SVD) with imputation was used to extract the principal components (PCs). The component loading values shown in the figure correspond to component scores of 4 DTNs along PC1 and PC2 that explain 56.9% and 27.1% of the total variance respectively. The top-10 pathways that appeared to be highly related to each of the 4 DTNs, which were obtained after computing the Euclidean distance between the component loading values and the component scores, are shown as square-shaped data points for the DTN of drugs effective in type 2 diabetes and not contraindicated in obesity, diamond-shaped data points for the DTN of drugs effective in obesity and not contraindicated in type 2 diabetes, triangle-shaped data points for the DTN of drugs effective in type 2 diabetes and contraindicated in obesity and cross-mark-shaped data points for the DTN of drugs effective in obesity and contraindicated in type 2 diabetes.

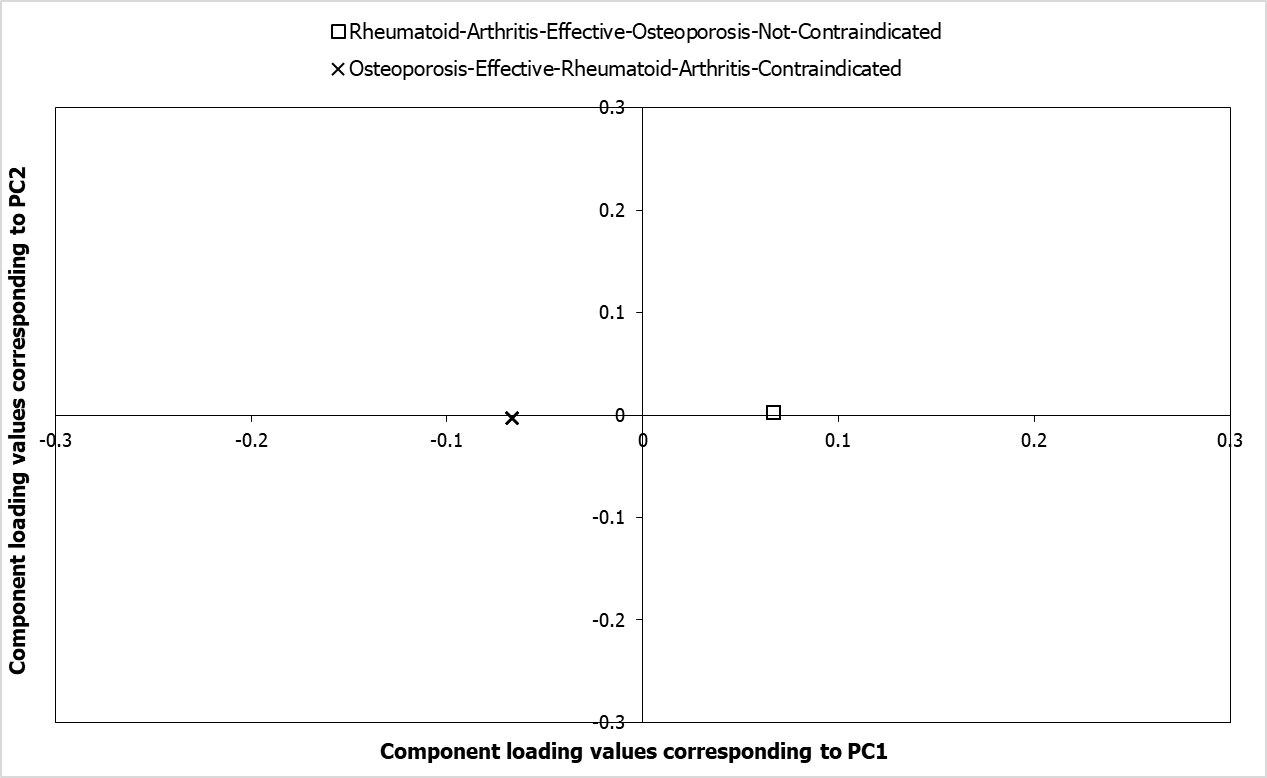

**Figure S4: Pathways associated with the target networks of rheumatoid arthritis and osteoporosis drugs.** Component loading values of 226 pathways associated with the drug target networks (DTNs) of rheumatoid arthritis (RA) and osteoporosis corresponding to PC1 and PC2 have been plotted along the X and Y axes respectively. PCA was performed with the p-values of enrichment of the pathways significantly associated (p-value < 0.05) with the DTNs of RA and osteoporosis. These values were transformed to –log_10_P values, which were then assembled into a data matrix containing pathways as rows and DTNs as columns. Unit variance scaling was applied across this matrix. Single value decomposition (SVD) with imputation was used to extract the principal components (PCs). The component loading values shown in the figure correspond to component scores of 4 DTNs along PC1 and PC2 that explain 100% and 0% of the total variance respectively. The top-10 pathways that appeared to be highly related to each of the 4 DTNs, which were obtained after computing the Euclidean distance between the component loading values and the component scores, are shown as square-shaped data points for the DTN of drugs effective in RA and not contraindicated in osteoporosis, diamond-shaped data points for the DTN of drugs effective in osteoporosis and not contraindicated in RA, triangle-shaped data points for the DTN of drugs effective in RA and contraindicated in osteoporosis and cross-mark-shaped data points for the DTN of drugs effective in osteoporosis and contraindicated in RA.

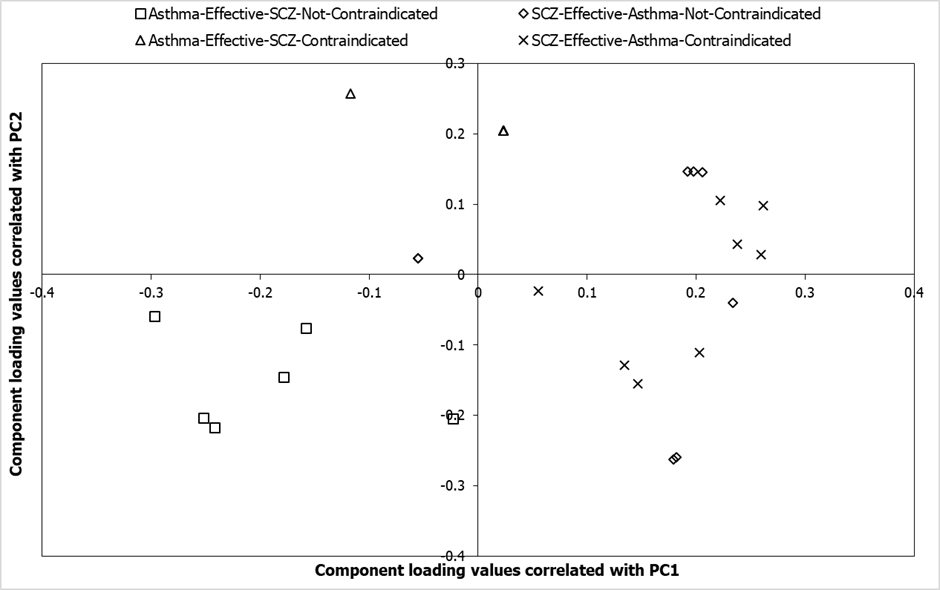
**Figure S5: Pathways associated with the target networks of asthma and schizophrenia drugs.** Component loading values of 35 pathways associated with the drug target networks (DTNs) of asthma and schizophrenia (SCZ) corresponding to PC1 and PC2 have been plotted along the X and Y axes respectively. PCA was performed with the p-values of enrichment of the pathways significantly associated (p-value < 0.05) with the DTNs of asthma and SCZ. These values were transformed to –log_10_P values, which were then assembled into a data matrix containing pathways as rows and DTNs as columns. Unit variance scaling was applied across this matrix. Single value decomposition (SVD) with imputation was used to extract the principal components (PCs). The component loading values shown in the figure correspond to component scores of 4 DTNs along PC1 and PC2 that explain 45.1% and 33.3% of the total variance respectively. The top-10 pathways that appeared to be highly related to each of the 4 DTNs, which were obtained after computing the Euclidean distance between the component loading values and the component scores, are shown as square-shaped data points for the DTN of drugs effective in asthma and not contraindicated in SCZ, diamond-shaped data points for the DTN of drugs effective in SCZ and not contraindicated in asthma, triangle-shaped data points for the DTN of drugs effective in asthma and contraindicated in SCZ and cross-mark-shaped data points for the DTN of drugs effective in SCZ and contraindicated in asthma.

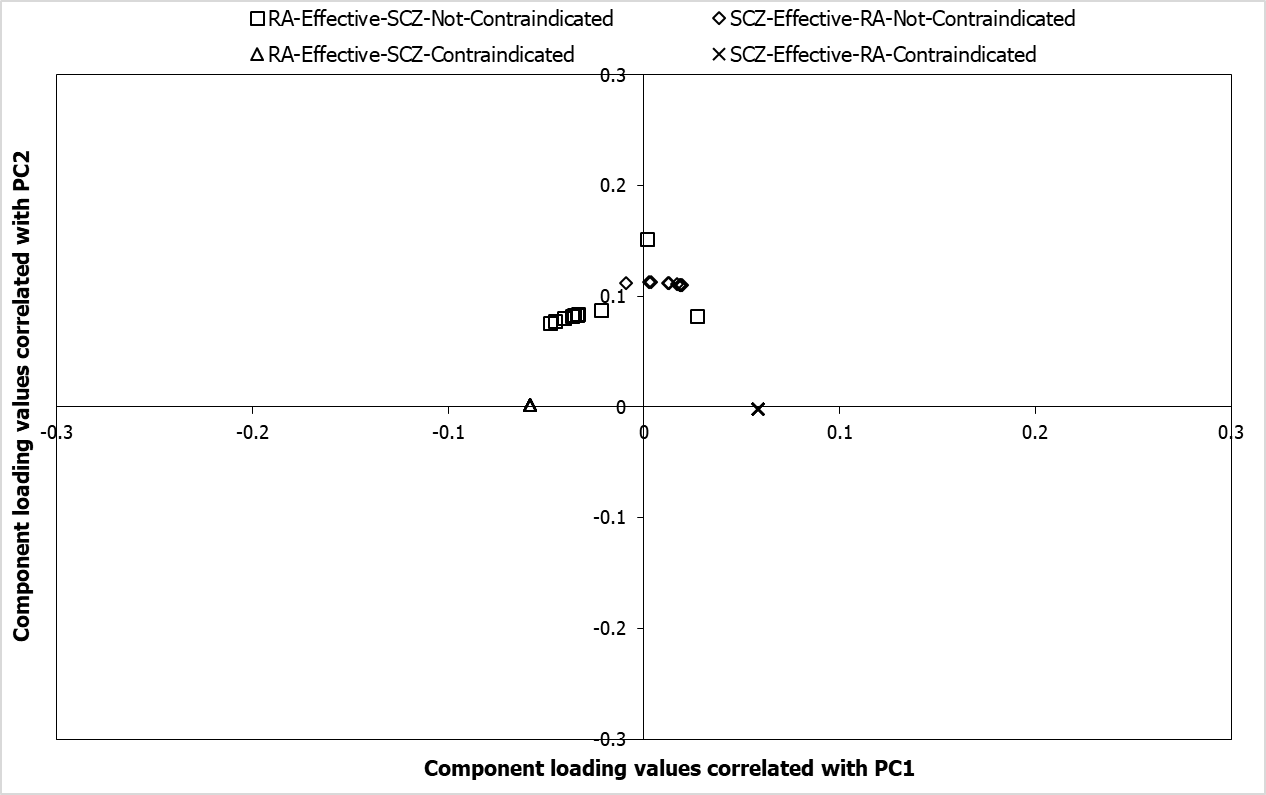
**Figure S6: Pathways associated with the target networks of rheumatoid and schizophrenia drugs.** Component loading values of 303 pathways associated with the drug target networks (DTNs) of rheumatoid arthritis (RA) and schizophrenia (SCZ) corresponding to PC1 and PC2 have been plotted along the X and Y axes respectively. PCA was performed with the p-values of enrichment of the pathways significantly associated (p-value < 0.05) with the DTNs of RA and SCZ. These values were transformed to –log_10_P values, which were then assembled into a data matrix containing pathways as rows and DTNs as columns. Unit variance scaling was applied across this matrix. Single value decomposition (SVD) with imputation was used to extract the principal components (PCs). The component loading values shown in the figure correspond to component scores of 4 DTNs along PC1 and PC2 that explain 66.3% and 28.5% of the total variance respectively. The top-10 pathways that appeared to be highly related to each of the 4 DTNs, which were obtained after computing the Euclidean distance between the component loading values and the component scores, are shown as square-shaped data points for the DTN of drugs effective in RA and not contraindicated in SCZ, diamond-shaped data points for the DTN of drugs effective in SCZ and not contraindicated in RA, triangle-shaped data points for the DTN of drugs effective in RA and contraindicated in SCZ and cross-mark-shaped data points for the DTN of drugs effective in SCZ and contraindicated in RA.

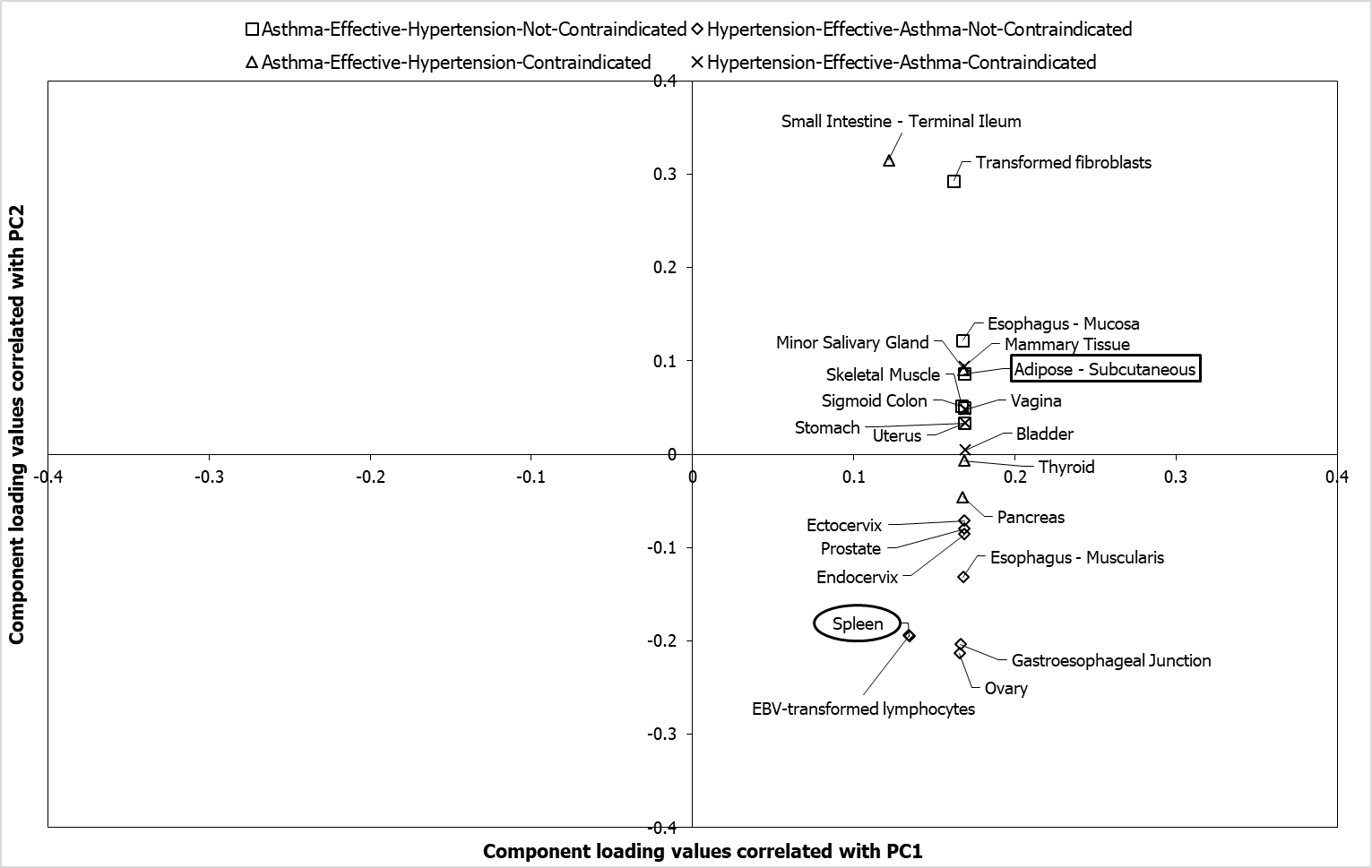
**Figure S7: Tissues associated with the target networks of asthma and hypertension drugs.** Component loading values of 38 tissues associated with the drug target networks (DTNs) of asthma and hypertension to PC1 and PC2 have been plotted along the X and Y axes respectively. PCA was performed with the p-values of enrichment of the tissues significantly associated (p-value < 0.05) with the DTNs of asthma and hypertension. These values were transformed to –log_10_P values, which were then assembled into a data matrix containing tissues as rows and DTNs as columns. Unit variance scaling was applied across this matrix. Single value decomposition (SVD) with imputation was used to extract the principal components (PCs). The component loading values shown in the figure correspond to component scores of 4 DTNs along PC1 and PC2 that explain 97% and 2.4% of the total variance respectively. The tissues that were exclusively associated with each of the 4 DTNs among the top-ten tissues that were identified to be highly related to the DTNs, after computing the Euclidean distance between the component loading values and the component scores, are shown as square-shaped data points for the DTN of drugs effective in asthma and not contraindicated in hypertension, diamond-shaped data points for the DTN of drugs effective in hypertension and not contraindicated in asthma, triangle-shaped data points for the DTN of drugs effective in asthma and contraindicated in hypertension and cross-mark-shaped data points for the DTN of drugs effective in hypertension and contraindicated in asthma. The tissues shown in circular and rectangular boxes were also identified to be highly specific to asthma and hypertension respectively by TSEA-DB (due to a significant enrichment of asthma/hypertension-associated variants).

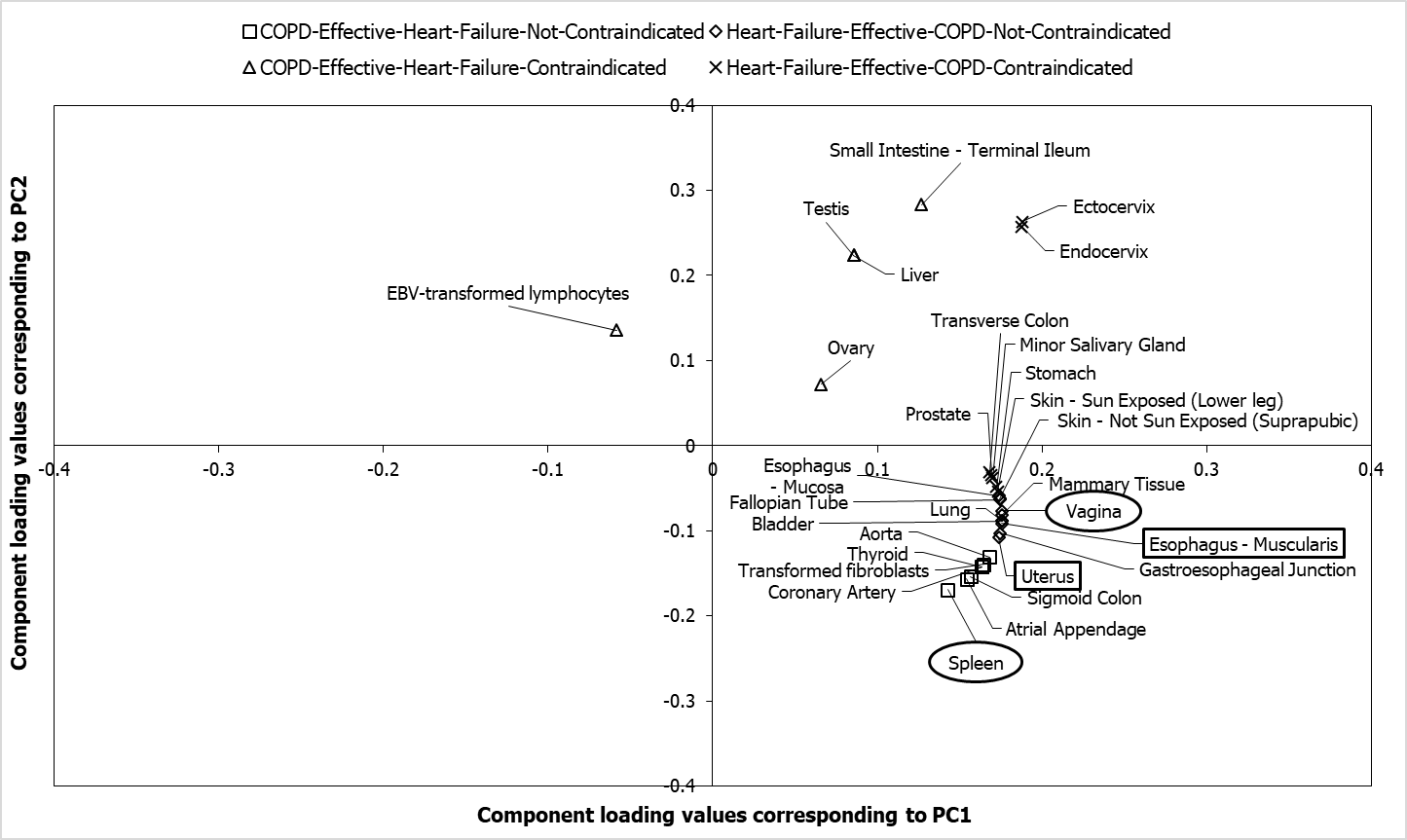
**Figure S8: Tissues associated with the target networks of chronic obstructive pulmonary disease and heart failure drugs.** Component loading values of 38 tissues associated with the drug target networks (DTNs) of chronic obstructive pulmonary disease (COPD) and heart failure to PC1 and PC2 have been plotted along the X and Y axes respectively. PCA was performed with the p-values of enrichment of the tissues significantly associated (p-value < 0.05) with the DTNs of COPD and heart failure. These values were transformed to –log_10_P values, which were then assembled into a data matrix containing tissues as rows and DTNs as columns. Unit variance scaling was applied across this matrix. Single value decomposition (SVD) with imputation was used to extract the principal components (PCs). The component loading values shown in the figure correspond to component scores of 4 DTNs along PC1 and PC2 that explain 79% and 14% of the total variance respectively. The tissues that were exclusively associated with each of the 4 DTNs among the top-ten tissues that were identified to be highly related to the DTNs, after computing the Euclidean distance between the component loading values and the component scores, are shown as square-shaped data points for the DTN of drugs effective in COPD and not contraindicated in heart failure, diamond-shaped data points for the DTN of drugs effective in heart failure and not contraindicated in COPD, triangle-shaped data points for the DTN of drugs effective in COPD and contraindicated in heart failure and cross-mark-shaped data points for the DTN of drugs effective in heart failure and contraindicated in COPD. The tissues shown in circular and rectangular boxes were also identified to be highly specific to COPD and heart failure respectively by TSEA-DB (due to a significant enrichment of COPD/heart failure-associated variants).

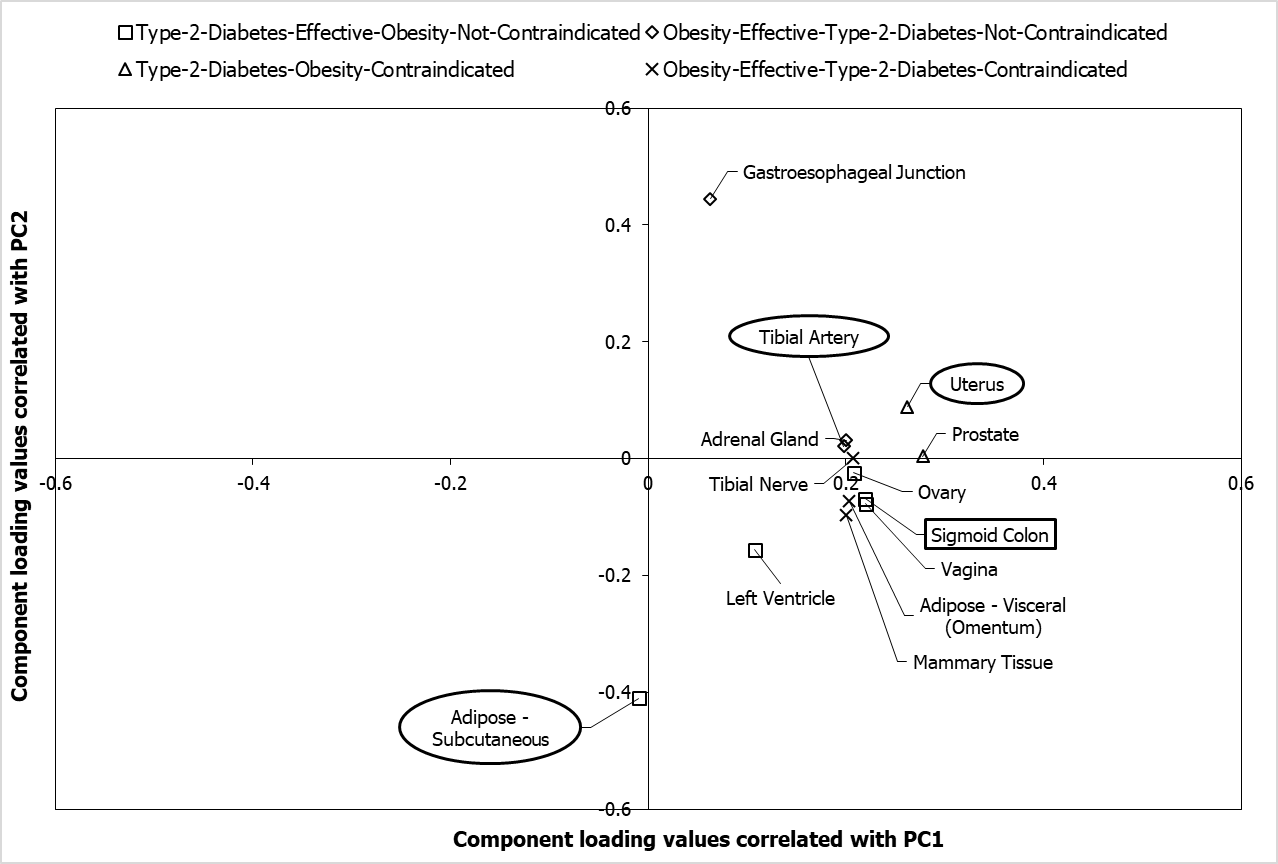
**Figure S9: Tissues associated with the target networks of type 2 diabetes and obesity drugs.** Component loading values of 26 tissues associated with the drug target networks (DTNs) of type 2 diabetes and obesity to PC1 and PC2 have been plotted along the X and Y axes respectively. PCA was performed with the p-values of enrichment of the tissues significantly associated (p-value < 0.05) with the DTNs of type 2 diabetes and obesity. These values were transformed to –log_10_P values, which were then assembled into a data matrix containing tissues as rows and DTNs as columns. Unit variance scaling was applied across this matrix. Single value decomposition (SVD) with imputation was used to extract the principal components (PCs). The component loading values shown in the figure correspond to component scores of 4 DTNs along PC1 and PC2 that explain 68% and 21% of the total variance respectively. The tissues that were exclusively associated with each of the 4 DTNs among the top-ten tissues that were identified to be highly related to the DTNs, after computing the Euclidean distance between the component loading values and the component scores, are shown as square-shaped data points for the DTN of drugs effective in type 2 diabetes and not contraindicated in obesity, diamond-shaped data points for the DTN of drugs effective in obesity and not contraindicated in type 2 diabetes, triangle-shaped data points for the DTN of drugs effective in type 2 diabetes and contraindicated in obesity and cross-mark-shaped data points for the DTN of drugs effective in obesity and contraindicated in type 2 diabetes. The tissues shown in circular and rectangular boxes were also identified to be highly specific to type 2 diabetes and obesity respectively by TSEA-DB (due to a significant enrichment of type 2 diabetes/obesity-associated variants).

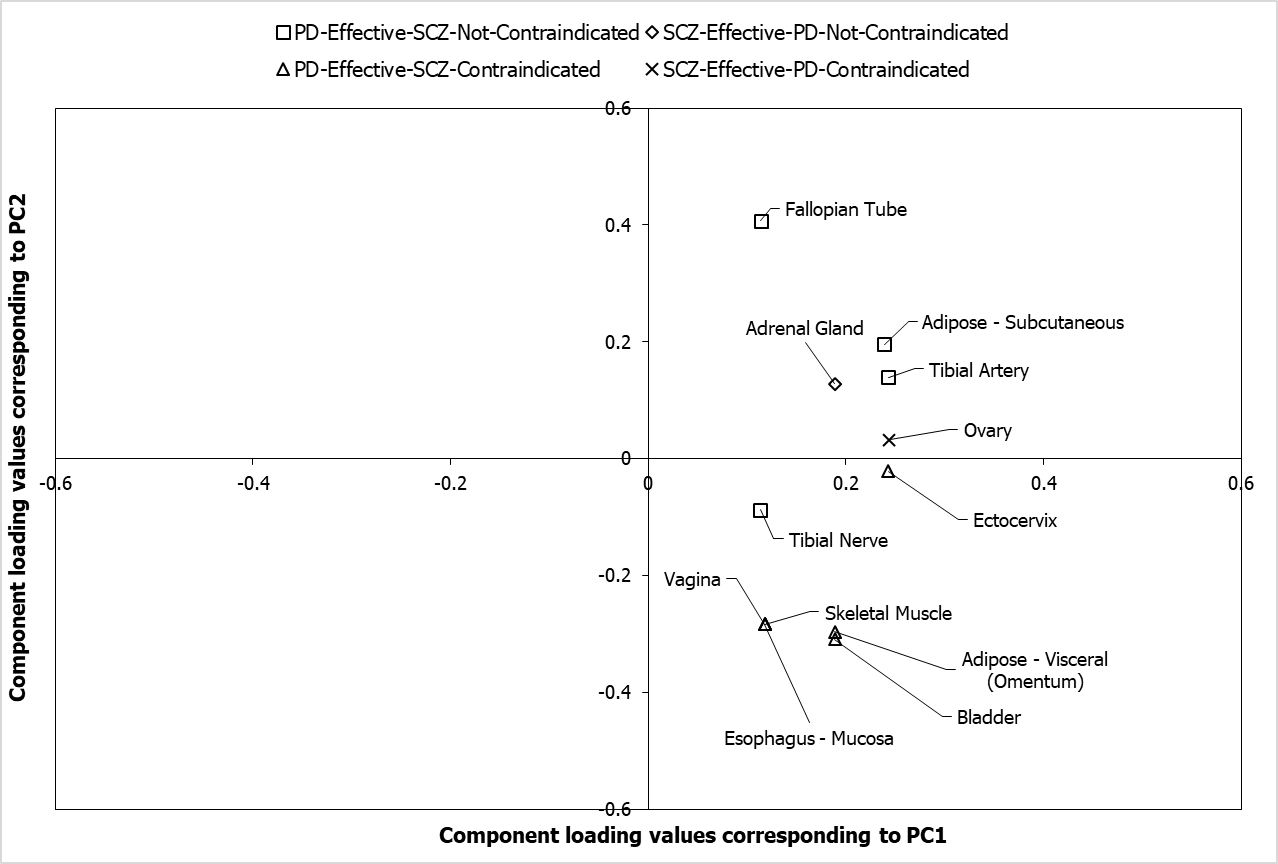
**Figure S10: Tissues associated with the target networks of Parkinson’s disease and schizophrenia drugs.** Component loading values of 24 tissues associated with the drug target networks (DTNs) of Parkinson’s disease (PD) and schizophrenia (SCZ) to PC1 and PC2 have been plotted along the X and Y axes respectively. PCA was performed with the p-values of enrichment of the tissues significantly associated (p-value < 0.05) with the DTNs of PD and SCZ. These values were transformed to –log_10_P values, which were then assembled into a data matrix containing tissues as rows and DTNs as columns. Unit variance scaling was applied across this matrix. Single value decomposition (SVD) with imputation was used to extract the principal components (PCs). The component loading values shown in the figure correspond to component scores of 4 DTNs along PC1 and PC2 that explain 93% and 4% of the total variance respectively. The tissues that were exclusively associated with each of the 4 DTNs among the top-ten tissues that were identified to be highly related to the DTNs, after computing the Euclidean distance between the component loading values and the component scores, are shown as square-shaped data points for the DTN of drugs effective in PD and not contraindicated in SCZ, diamond-shaped data points for the DTN of drugs effective in SCZ and not contraindicated in PD, triangle-shaped data points for the DTN of drugs effective in PD and contraindicated in SCZ and cross-mark-shaped data points for the DTN of drugs effective in SCZ and contraindicated in PD. The tissues shown in circular and rectangular boxes were also identified to be highly specific to PD and SCZ respectively by TSEA-DB (due to a significant enrichment of PD/SCZ-associated variants).

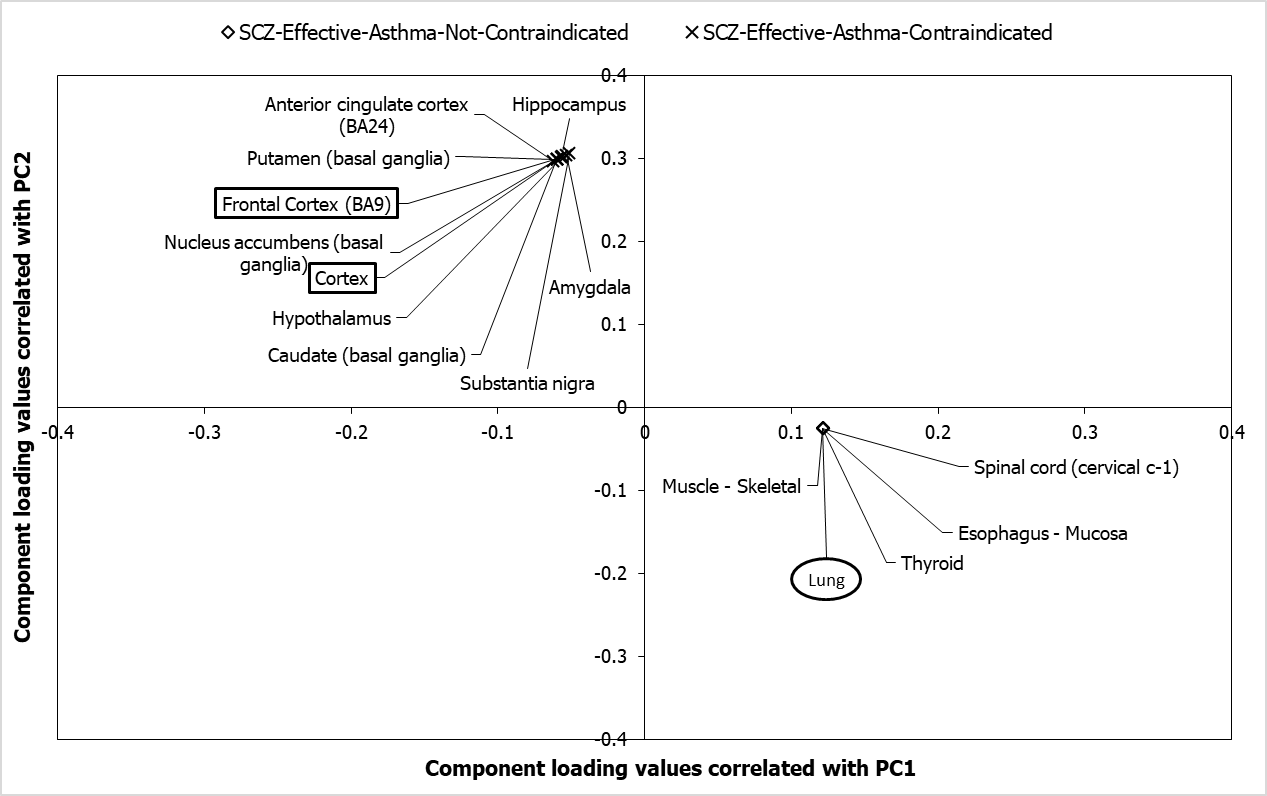
**Figure S11: Tissues associated with the target networks of asthma and schizophrenia drugs.** Component loading values of 53 tissues associated with the drug target networks (DTNs) of asthma and schizophrenia (SCZ) to PC1 and PC2 have been plotted along the X and Y axes respectively. PCA was performed with the p-values of enrichment of the tissues significantly associated (p-value < 0.05) with the DTNs of asthma and SCZ. These values were transformed to –log_10_P values, which were then assembled into a data matrix containing tissues as rows and DTNs as columns. Unit variance scaling was applied across this matrix. Single value decomposition (SVD) with imputation was used to extract the principal components (PCs). The component loading values shown in the figure correspond to component scores of 4 DTNs along PC1 and PC2 that explain 81% and 18.5% of the total variance respectively. The tissues that were exclusively associated with each of the 4 DTNs among the top-ten tissues that were identified to be highly related to the DTNs, after computing the Euclidean distance between the component loading values and the component scores, are shown as diamond-shaped data points for the DTN of drugs effective in SCZ and not contraindicated in asthma and cross-mark-shaped data points for the DTN of drugs effective in SCZ and contraindicated in asthma. The tissues shown in circular and rectangular boxes were also identified to be highly specific to asthma and SCZ respectively by TSEA-DB (due to a significant enrichment of asthma/SCZ-associated variants).

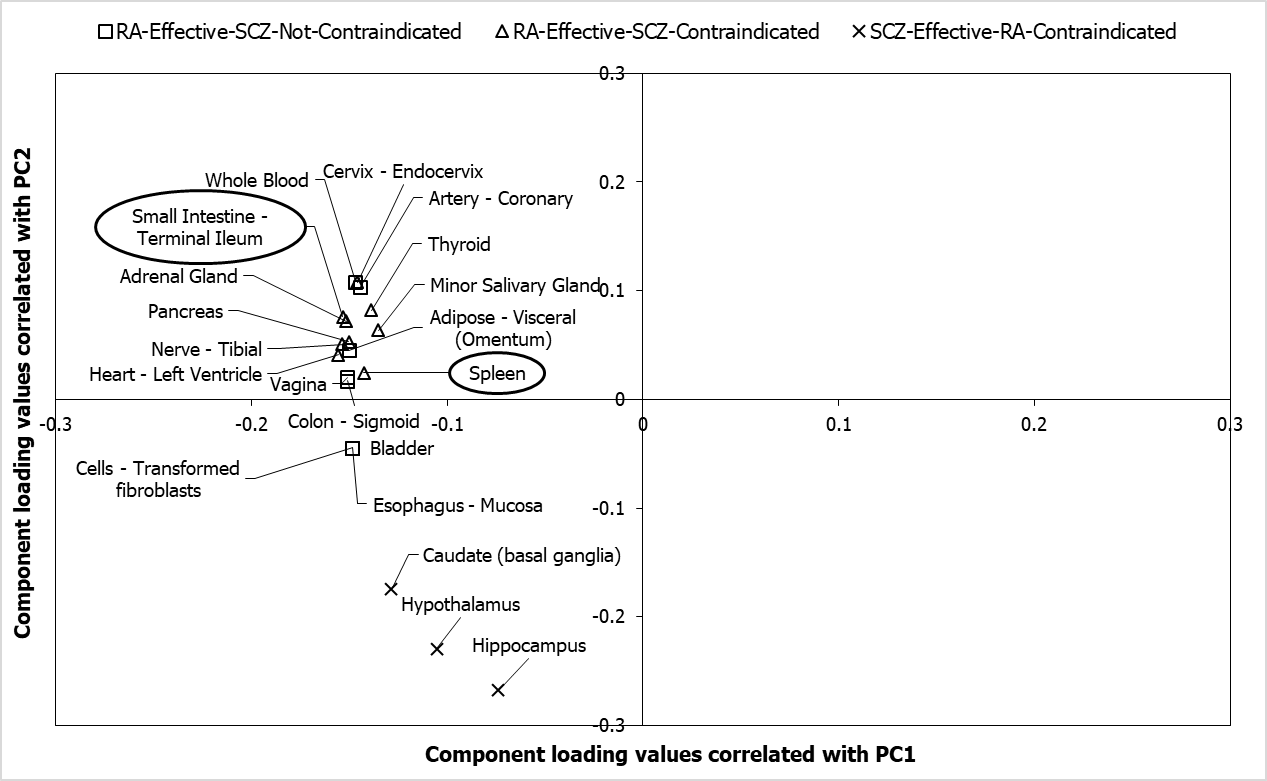
**Figure S12: Tissues associated with the target networks of rheumatoid arthritis and schizophrenia drugs.** Component loading values of 53 tissues associated with the drug target networks (DTNs) of rheumatoid arthritis (RA) and schizophrenia (SCZ) to PC1 and PC2 have been plotted along the X and Y axes respectively. PCA was performed with the p-values of enrichment of the tissues significantly associated (p-value < 0.05) with the DTNs of RA and SCZ. These values were transformed to –log_10_P values, which were then assembled into a data matrix containing tissues as rows and DTNs as columns. Unit variance scaling was applied across this matrix. Single value decomposition (SVD) with imputation was used to extract the principal components (PCs). The component loading values shown in the figure correspond to component scores of 4 DTNs along PC1 and PC2 that explain 75.3% and 19.8% of the total variance respectively. The tissues that were exclusively associated with each of the 4 DTNs among the top-ten tissues that were identified to be highly related to the DTNs, after computing the Euclidean distance between the component loading values and the component scores, are shown as square-shaped data points for the DTN of drugs effective in RA and not contraindicated in SCZ, triangle-shaped data points for the DTN of drugs effective in RA and contraindicated in SCZ and cross-mark-shaped data points for the DTN of drugs effective in SCZ and contraindicated in RA. The tissues shown in circular and rectangular boxes were also identified to be highly specific to RA and SCZ respectively by TSEA-DB (due to a significant enrichment of RA/SCZ-associated variants).

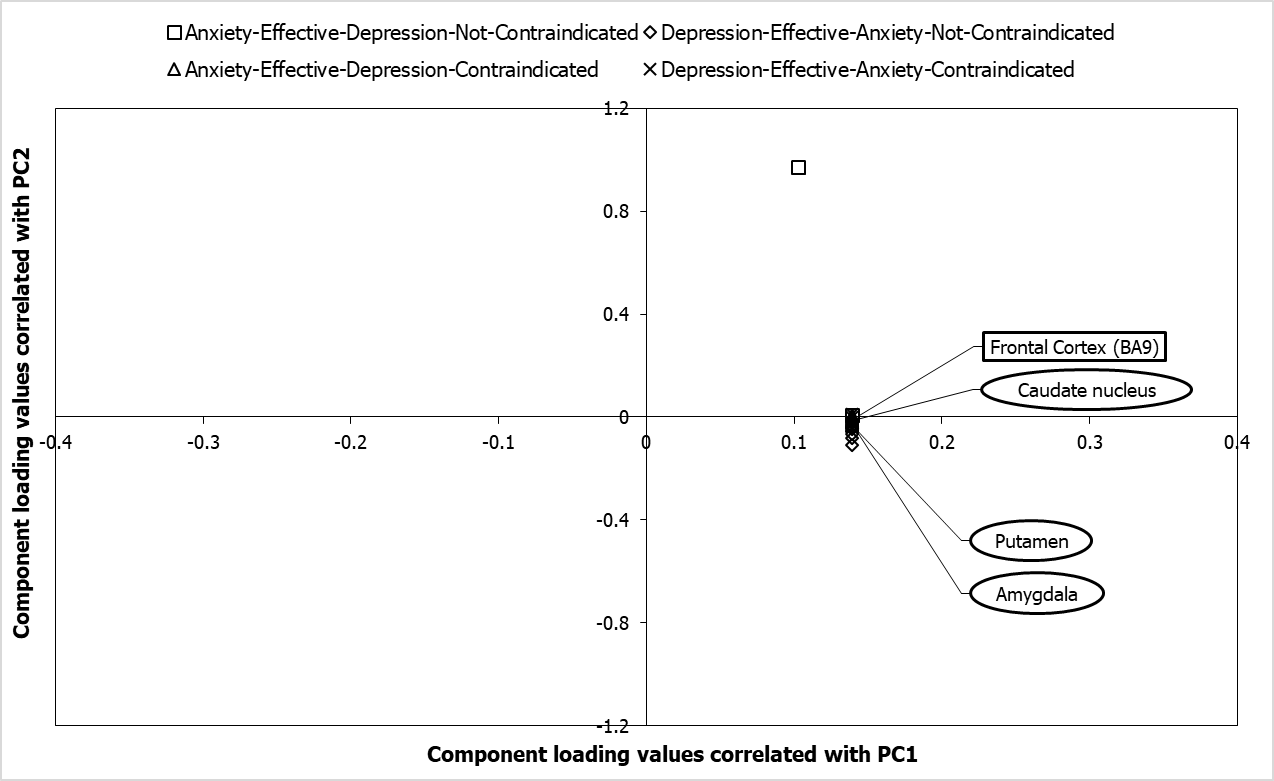

**Figure S13: Tissues associated with the target networks of anxiety and depression drugs.** Component loading values of 52 tissues associated with the drug target networks (DTNs) of anxiety and depression corresponding to PC1 and PC2 have been plotted along the X and Y axes respectively. PCA was performed with the p-values of enrichment of the tissues significantly associated (p-value < 0.05) with the DTNs of anxiety and depression. Note that all the genes that showed expression (transcripts per million (TPM) > 2) in the tissues were included for the enrichment analysis. The p-values were then log-transformed and assembled into a data matrix containing tissues as rows and DTNs as columns. Unit variance scaling was applied across this matrix. Single value decomposition (SVD) with imputation was used to extract the principal components (PCs). The component loading values shown in the figure correspond to component scores of 4 DTNs along PC1 and PC2 that explain 99.7% and 0.3% of the total variance respectively. The tissues that were exclusively associated with each of the 4 DTNs among the top-ten tissues that were identified to be highly related to the DTNs, after computing the Euclidean distance between the component loading values and the component scores, are shown as square-shaped data points for the DTN of drugs effective in anxiety and not contraindicated in depression, diamond-shaped data points for the DTN of drugs effective in depression and not contraindicated in anxiety, triangle-shaped data points for the DTN of drugs effective in anxiety and contraindicated in depression and cross-mark-shaped data points for the DTN of drugs effective in depression and contraindicated in anxiety. The tissues shown in circular and rectangular boxes were also identified to be highly specific to anxiety and depression respectively by TSEA-DB (due to a significant enrichment of anxiety/depression-associated variants).

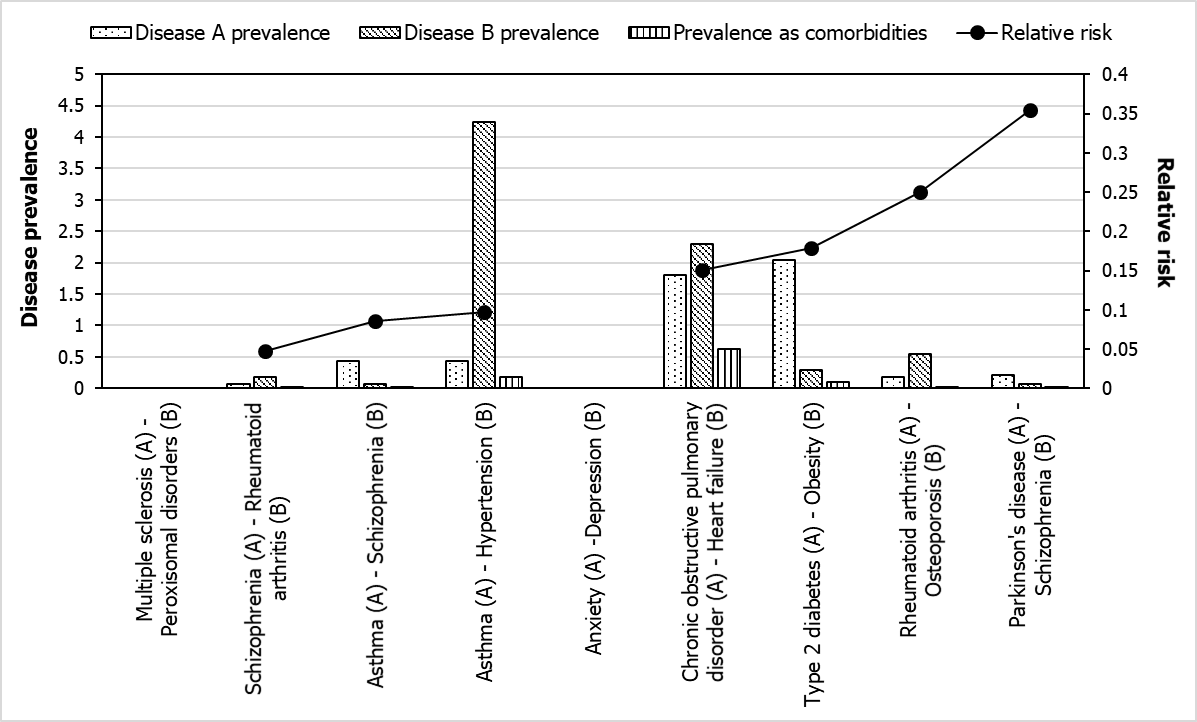

**Figure S14: Comparison of the relative risk of disease comorbidity with disease prevalence.** The graph shows the relationship of the relative risks of seven pairs of diseases (black data points) with the individual prevalence of the diseases and the prevalence of the diseases as comorbid conditions (bar graphs). The relative risks of Rheumatoid arthritis – Osteoporosis and Parkinson’s disease – Schizophrenia seem to have been overestimated due to the low prevalence of these diseases in the population.
